# A multiscale modeling - experimental framework reveals the interplay between drivers of *Drosophila* wing morphogenesis

**DOI:** 10.64898/2026.11.12.765773

**Authors:** Ethan Nowaski, Benjamin Speybroeck, Navaira Sherwani, Emerald Win, Zhiliang Xu, Weitao Chen, Jeremiah Zartman, Mark Alber

## Abstract

Robust organ formation depends on the coordinated regulation of cellular mechanics, yet how mechanobiological processes such as controlled cell-cell and cell-matrix adhesion, actomyosin contractility, membrane stiffness, and internal pressure interlink to generate an organ’s architecture remains poorly understood. Here, we systematically investigated underlying mechanobiology principles using *Drosophila* wing disc eversion, the metamorphic conversion of a larval epithelial sac into a functional wing. By integrating quantitative imaging experiments with a novel multiscale modeling environment we termed MORPHIOME, we extensively tested the roles of individual drivers of tissue morphogenesis, as well as their different combinations. Our analysis suggests the following multi-level sequence of morphogenetic processes defining early wing metamorphosis. First, the localized detachment of the basal Extra Cellular Matrix (ECM) at the dorsal-ventral compartment boundary initiates folding and boundary forces reorient and shear the wing disc pouch to guide the proximal ends of the pouch together. Then, patterned actomyosin contractility closes the fold. Also, cell rearrangements promote tissue elongation all through the process. In sum, these results show how organogenesis relies on a combination of subcellular force-generation mechanisms and tissue-level deformations. The developed integrated, data-calibrated nonlinear MORPHIOME framework can serve as a versatile general tool for investigating the mechanisms of tissue development and regeneration.

## Introduction

How can a single, relatively spherical cell embryo systematically transform itself into a coherent community of thousands, millions, or billions of cells that exhibit an increasingly complex multilayered architecture? While much progress has been made in sequencing genes and elucidating protein-protein interactions, our understanding of the multiscalar biophysical principles that define organogenesis remains incomplete. The remarkable process of organogenesis is driven by precise changes in cellular shape and number through the action of complex biochemical signaling networks acting under specific constraint processes^1,2^. Generating a predictive understanding of organogenesis requires the development of advanced computational modeling tools. Such predictive computational models can serve as design-based engineering frameworks for designing and testing targeted treatments of cancer, birth defects, and impaired wound healing ^3,4^.

A particularly dramatic example of how complex biological morphologies develop is the metamorphosis of the fruit fly (*Drosophila melanogaster*) wing disc into a fully functional wing. As a prime biological model system with unparalleled genetic tools and ease of optical imaging and manipulation, *Drosophila* enables the investigation of how living structures are generated across genetic, cellular, and tissue scales^4,5^. The wing disc is a sac-like epithelial organ composed of two opposed epithelial layers: the pseudostratified disc proper and the squamous peripodial epithelium. The disc proper is further subdivided into the pouch, hinge, notum, which exhibit morphogenetic behaviors^5–7^. The basal surface of the wing disc epithelia is associated with a basement membrane (BM), a specialized form of extracellular matrix (ECM). The BM is composed primarily of collagen IV, laminin, nidogen, and perlecan and provides structural and mechanical support to the tissue^8^. The pouch consists of tall, narrow, and polarized epithelial cells, with their apical surface facing the lumen of the sac and their basal surface contacting the BM^8^. This work primarily focuses on the morphogenetic changes occurring in the pouch region of the wing disc, as it dramatically changes shape from a dome shape into a bilayer during eversion and gives rise to the adult wing blade^9^. The pouch is subdivided into four major spatial domains that arise from the orthogonal anterior-posterior (AP) and dorsal-ventral (DV) compartment boundaries^10^. Finally, the pouch connects to the hinge and notum, which will form part of the thorax and wing connective tissue in the adult. As development progresses, patterned signaling along the anterior-posterior (AP) and dorsal-ventral (DV) axes establishes positional information within the wing disc. The morphogens Decapentaplegic (Dpp) and Wingless (Wg) help pattern the future wing blade, hinge, and notum regions^11–14^. Concurrently, signaling pathways, including Hedgehog, Notch, and Hippo, integrate spatial morphogen cues with growth control to coordinate tissue size and shape^15,16^.

By the late third instar, the wing disc exhibits pronounced curvature and mechanical tension^17,18^, setting the stage for the metamorphic remodeling that occurs during eversion^19–21^. The larval-to-pupal transition is initiated by a spike of the hormone ecdysone secreted from the prothoracic glands, which signals a cascade of key cellular and tissue-level changes in the organ. In this process, the proximal tips of the pouch converge, the pouch rotates approximately 90 degrees, then folds along the DV axis and elongates to form a bilayer^22–24^. This process shows significant changes in cell height and curvature (Fig 1. C, D, E). Following eversion, differentiation and vein patterning progress to generate the mature wing morphology over several days^25,26^.

**Figure 1.**
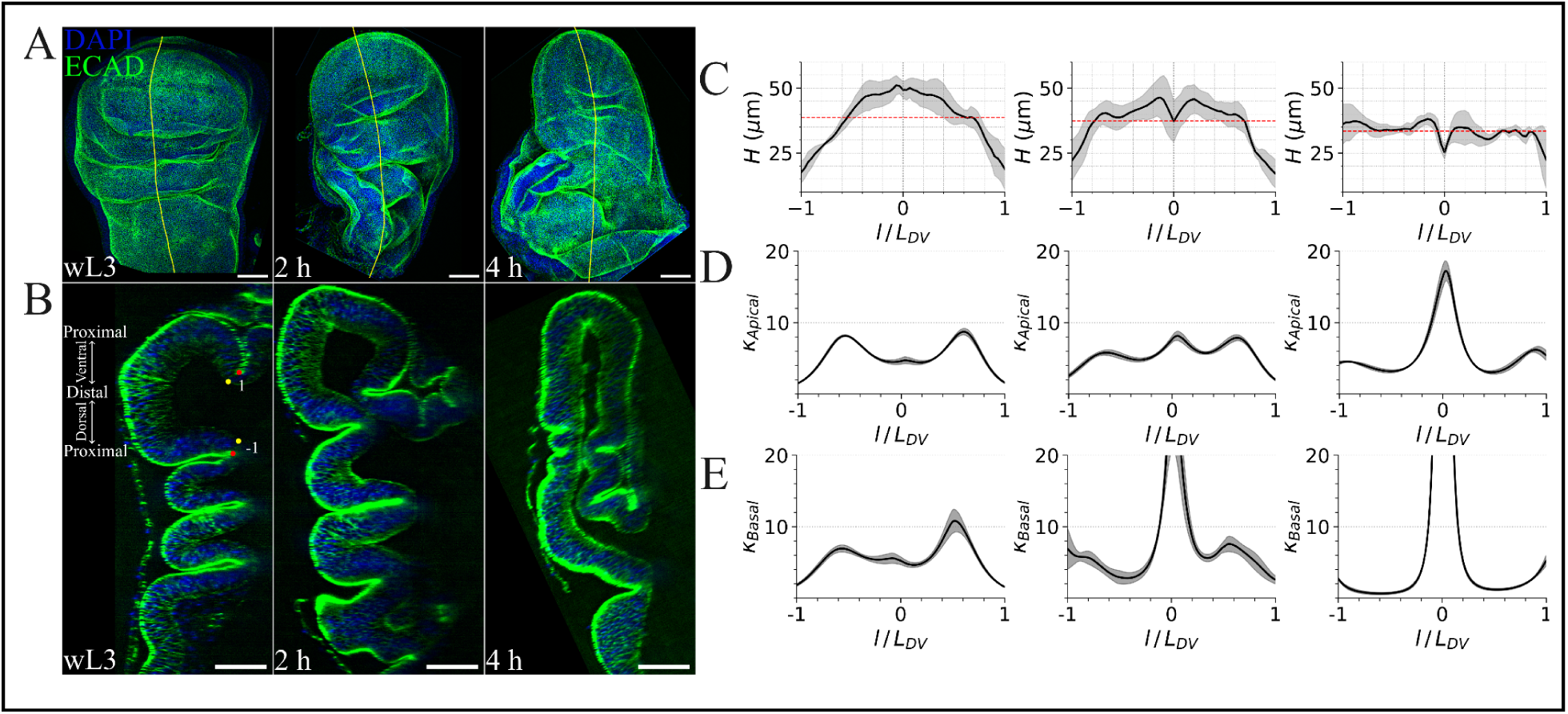
**Dynamic changes of tissue shape and structure during eversion**. **A.** Representative images of the wing disc at wL3, 2 h APF, and 4 h APF. **B.** Representative cross-sectional images along the dorsal-ventral (DV) axis. **C.** Tissue height profiles along the DV axis of each developmental stage. Solid lines represent the mean and shaded regions represent ± SD. **D.** Local apical curvature profiles along the DV axis. **E.** Local basal curvature profiles along the DV axis. For curvature profiles, solid lines represent the mean and shaded regions represent ± SEM. n=6 for wL3 and 2 h APF wing discs, and n=3 for 4 h APF wing discs. Scale bars, 50 µm.

Despite its critical role in the developmental process, eversion is an understudied phase of development compared to the other stages of wing growth. Eversion occurs during a relatively brief developmental window and involves rapid tissue remodeling, making it difficult to experimentally capture with high spatiotemporal resolution^5,6,27^. The rapid rotation, folding, and elongation generate complex deformations that are difficult to quantify using conventional image analysis and modeling approaches^18,24,28^. Further, eversion requires coordinated mechanical and biochemical interactions between the pouch, hinge, peripodial membrane, and ECM, further complicating mechanistic interpretation of tissue-scale behaviors^7,29,30^. Therefore, unraveling the mechanisms of governing this developmental stage is essential to understanding the principles of tissue morphogenesis and robust organ growth. To propose mechanical mechanisms driving these shape changes, we evaluated candidate forces involving contractility, cell rearrangements, cell-ECM interactions, and external boundary dynamics to determine which mechanisms, alone or in combination, are sufficient to reproduce the observed transformations.

An increasing number of computational mechanical, physics-informed computational modeling frameworks have been developed to study tissue development and growth. These span from continuum descriptions of bulk tissue to discrete, cell-based representations^1,31,32^. Within the context of the *Drosophila* wing imaginal disc, continuum formulations, built upon the finite element methods, have recently been applied to studying tissue deformation and bending of the disc epithelium^28,33–35^. Discrete, cell-based models, such as the one used in this paper, provide detailed descriptions of the processes involving individual cells, cell-cell adhesion and cell-ECM adhesion. These are customarily grouped into lattice-based and off-lattice models. A popular framework within the lattice-based approaches is Cellular Potts models, which represent each cell as a cluster of lattice sites whose shape is updated by an energy-based Monte-Carlo algorithm^36,37^. Such models have been used to study morphogenesis, cell aggregation and sorting, thrombus formation, and other biological problems^38,39^. Off-lattice models use several representations of the individual cell depending on the desired scale of resolution: a single particle in agent-based models^40^, a polygon made up of a small collection of nodes in vertex models^41,42^, and particles (nodes) defining the membrane and other cell structures in fine detail in subcellular element (SCE) models^43,44^. Since the latter represents a cell as a group of nodes connected by springs, it is capable of reproducing biologically relevant cell mechanical properties, shapes, and structures when embedded within a tissue^18,45,46^. This general modeling approach has been previously biologically calibrated and used for studying the earlier larval stages of the *Drosophila* wing disc cross-section along the anterior-posterior axis^18,46^.

In this paper, we describe a novel pseudo-3D off-lattice nonlinear **M**ultiscale **OR**gan-level **PH**ysics-**I**nf**O**rmed **M**odeling **E**nvironment (MORPHIOME) of the *Drosophila* wing disc cross-section for the dorsal-ventral (DV) axis (see Fig. 2). Computational simulations are used to suggest potential biophysical drivers of *Drosophila* wing disc eversion, and test whether hypothesized mechanisms are sufficient to cause the shape changes seen in time sequences of experimental images of eversion.

**Figure 2.**
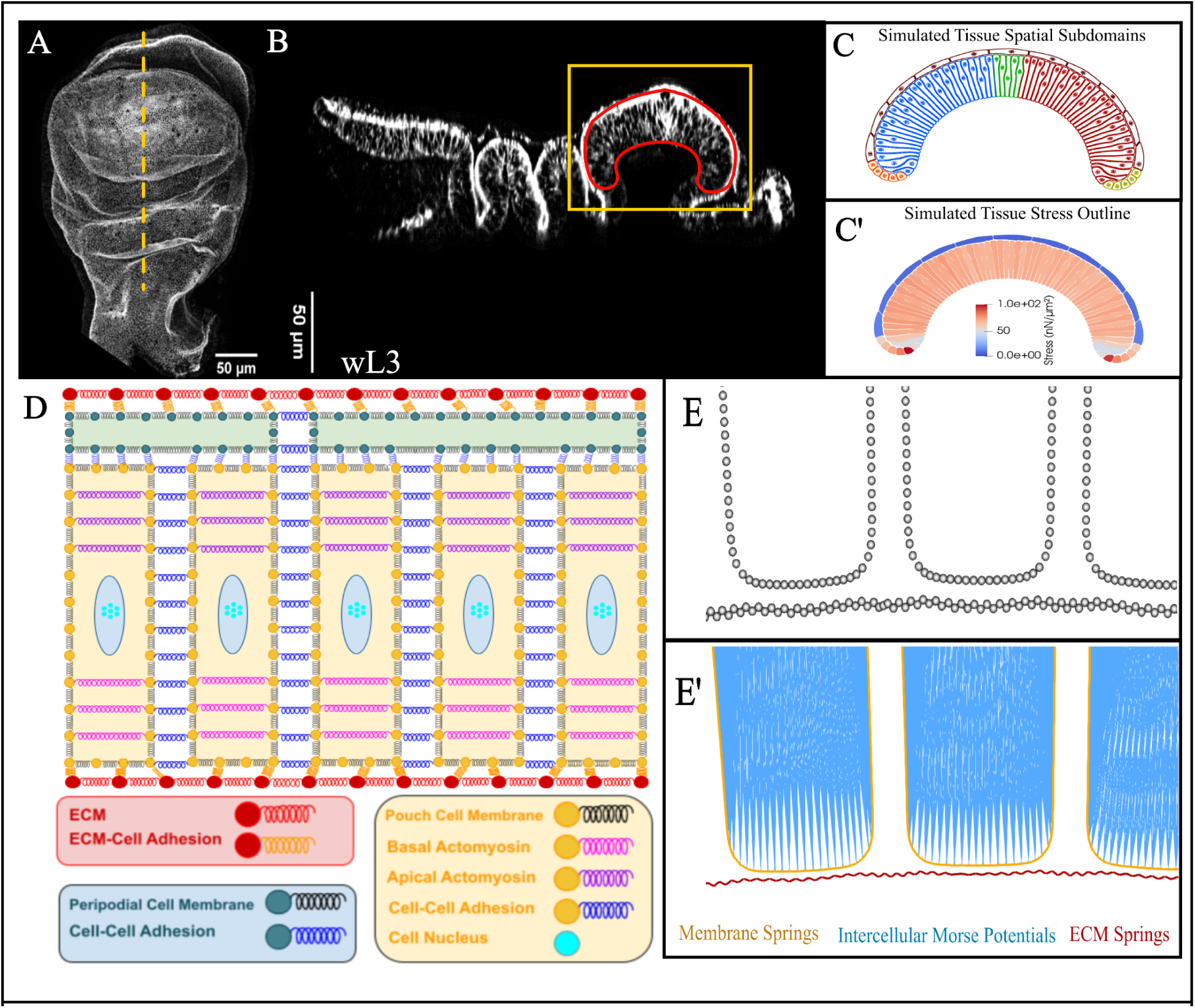
Description of the MORPHIOME framework for simulating wing disc eversion. A, **B**. Experimental images of the wing disc pouch at wL3 demonstrating the location of the DV cross-section and cross-section itself. **C**. Simulated tissue of the wing disc represented by nodes and color coded by spatial subdomains. The dorsal proximal region (DPR) is colored blue, the ventral proximal region (VPR) red, the dorsal-ventral boundary (DVB) green, the dorsal boundary cells orange, the ventral boundary cells yellow, and the squamous peripodial cells brown. **C’.** Simulated tissue of the wing disc DV cross-section represented by nodes and edges. As a simplification, the boundary of the pouch proper is connected with cuboidal cells that stand-in for the more complicated tissues connecting the pouch-interacting peripodial cells and pouch cells. Stress (in *nN*/µ*m*^2^) is calculated as the accumulated forces acting on a node at a given time point. **D.** Diagram of the MORPHIOME with subcellular nodes connected by springs representing the ECM (red) and cell (yellow/pouch and green/peripodial) structures. **E.** Zoomed-in visualization of the nodes showing the pouch cell and the basal ECM nodes. **E’.** The same visualization with physical interactions within the cells and ECM color-coded by interaction type.

The pseudo-3D MORPHIOME, described in this paper, couples three distinct spatial scales:

● **Micro-scale (Subcellular)** level comprises individual nodes connected through linear springs and other nonlinear energy potentials;
● **Meso-scale (Cellular)** level defines groups of nodes forming different cellular structures such as the columnar pouch cells, squamous peripodial cells, and the ECM;
● **Macro-scale (Tissue/Organ)** level encompasses the tissue along the dorsal-ventral axis cross-section of the wing disc, combining the eversion pouch domain with the rest of the tissue bounded by ECM and taking into account application of dynamic forces on the boundary of the domain.

Notably, linear models are generally only applicable for small deformations. Because wing disc eversion involves large deformations, spatial heterogeneities, and complex interacting forces, the nonlinear mechanical model is essential for producing outcomes that match experimental live-imaging data. By systematically isolating and perturbing model mechanisms against quantitative live-imaging data, apical/basal curvature profiles, and cell-height dynamics, the nonlinear MORPHIOME demonstrates that individual internal cellular mechanisms are insufficient on their own. Instead, successful eversion relies on the nonlinear coupling between localized subcellular force generators and macro-scale structural boundary forces.

To define and test the role of system-level interactions between specific morphogenetic processes, we developed and calibrated a new MORPHIOME to match the initial tissue shape, obtained as an equilibrium under a selected set of parameters, with experimental images at the beginning stage of eversion, wL3 (see Fig. 2B, C). Calibrated MORPHIOME was used to test the role of patterned actomyosin contractility, local ECM dynamics, active forces acting on the eversion domain boundary, and cell rearrangement in morphological changes leading to experimentally observed dynamical eversion shape formation. These potential mechanisms were tested, both individually as well as in combination with each other. We demonstrate that the complexity of wing disc eversion can be described as a “network” of concurrent morphogenetic events occurring downstream of the initial ecdysone pulse: (1) External connective tissue generates boundary forces and rotates the wing disc pouch by 90 degrees; (2) The formation of the folded bilayer depends on both basal actomyosin contractility and local delamination of the cells from the basal ECM at the DV boundary; and (3) Cell intercalation leading to local rearrangements contribute to the extension of the wing disc bilayer. In sum, this work provides new insights into the systems-level action of multiple concurrent morphogenetic processes shaping the pupal wing.

## Results

### Patterned basal actomyosin and membrane contraction produces robust fold formation but is insufficient to capture the intermediate dynamics of wing bilayer formation

Non-muscle Myosin II (NMII), an actin-based motor protein, generates contractile forces within the actomyosin cytoskeletal networks necessary for epithelial remodeling, including cell-shape changes, tissue bending, and morphogenesis^31,47–49^. For example, during *Drosophila* ventral furrow formation, pulsed actomyosin contractions drive apical constriction, and, during dorsal closure, pulsatile contractile forces contribute to directed tissue movement^19,50^. Therefore, we investigated whether spatially patterned actomyosin contractility is sufficient to generate the shape changes observed in experimental data. To vary actomyosin contractility *in silico*, we represented contractile forces as active, internal spring elements spanning the lateral membranes of the columnar pouch cells above and below nuclei (Fig. 3A, A’). In particular, we modeled varying levels of actomyosin-driven contractility by tuning spring constant values, which enabled us to test how patterned contractile forces can drive tissue-scale changes in the pouch morphology to generate the wing bilayer.

**Figure 3.**
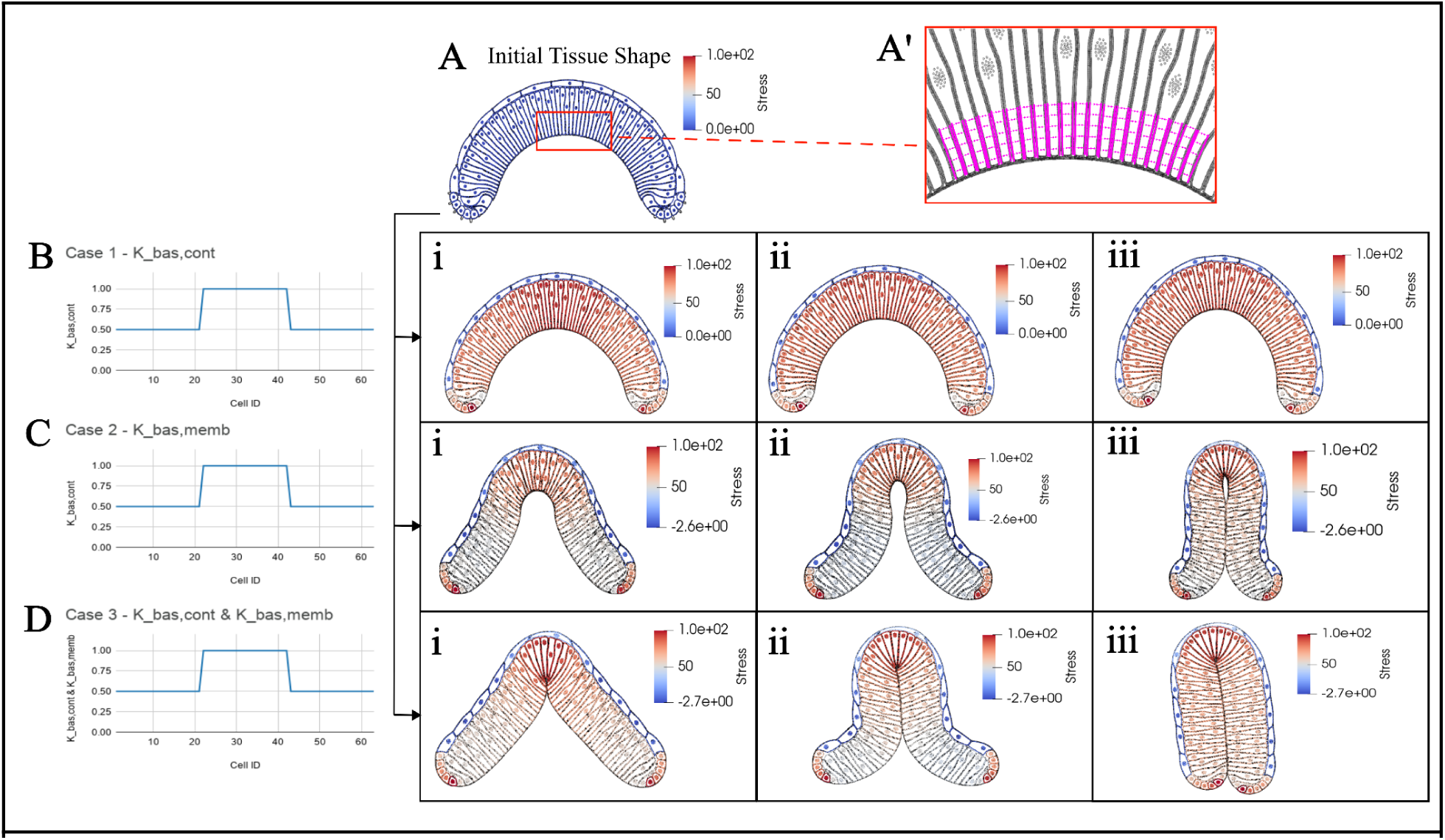
Medial-basal patterning of actomyosin contractility results in fold closure. The simulated tissue representing the pouch is divided into three subdomains: proximal dorsal, central (the dorsal ventral boundary region), and proximal ventral. This division is defined as spatially patterned basal membrane contractility. The color encodes the total stress magnitude (in *nN*/µ*m*^2^). **A.** The initial tissue shape in simulation reflects the initial shape in experimental data (see Fig.1B-ii). **A’.** Representation of the basolateral actomyosin contractile forces in magenta within the central subdomain of pouch cells. **B.** Simulated pouches with patterned basolateral actomyosin contractility fail to form an appropriate bilayered tissue shape. **C.** A simulated pouch with patterned basolateral membrane contractility generates a fold that is incompletely sealed at the DV boundary. **D.** A simulated pouch with coupled patterning of basal actomyosin and basal membrane contractility achieves a folded structure similar to experimental cross-sections observed at 4 h APF, but the direction of folding is opposite to experimental conditions.

Previous work published in Kumar et al. demonstrated the capability of the MORPHIOME computational framework to capture epithelial morphological change in the cross-section along the anterior-posterior axis of the wing disc pouch through patterning these contractile forces together with cell division^18^. Here, we developed a new model along the dorsal-ventral axis. We spatially divided the simulated pouch into thirds: the central dorsal-ventral boundary (DVB) region, which defines the distal region of the adult wing, flanked by two adjacent, more proximal regions of the wing disc, the dorsal proximal region (DPR) and ventral proximal region (VPR). We perturbed actomyosin contractility by decreasing the spring constant coefficients of the linear springs in the basolateral domain of the DPR and VPR relative to the DVB, to model higher contractility in the central subdomain. Such nonuniform actomyosin contractility is insufficient to generate the bilayer (Fig. 3B).

However, a specific persisting shape consistently formed at the site with higher basal actomyosin contractility, which was also observed during the progression found in experimental images of the wing bilayer formation (Fig. 1B). The cell’s wedge-like shape becomes more pronounced as the ratio of spring contractility increases (SI Fig. 2B-E).

Next, we applied the same nonuniform distribution to spring constants for springs connecting nodes representing the basal membrane (with globally uniform actomyosin contractility along the baso-lateral membrane) around the DVB. This simulated tissue developed into an incomplete bilayer structure, undergoing dorsal-ventral apposition but retaining a gap at the apex of the fold (Fig. 3C). As a follow-up iteration, we coupled both uneven basolateral contractility with uneven basal membrane contractility to generate a complete bilayer similar to experimental cross-sectional morphologies seen in the wing disc pouch at 4 h APF (Fig. 3D). The combination of both parameter perturbations represent a coupled biological process of actomyosin localized along both the basal membrane and the basolateral membrane, generating the final fold formation from a range of different initial tissue shape configurations, which indicates a level of robustness to variations in the initial shape. (Fig. S1).

Although the coupled basolateral and basal contractility of cells result in bilayer shape similar to 4 h APF, the intermediate simulated shapes deviate significantly from experimental observations. Specifically, experimental images from staged 2 h APF pupal wings suggest an “outside-in” closure mechanism. The proximal dorsal and ventral regions of the nascent wing bilayer make initial contact followed by the fold closing along the length of the central domain toward the DV boundary (Fig. 1B). In contrast, the simulation results in Fig. 3D exhibits a clear “distal-to-proximal” closure. Due to the spatial patterning of the contractility parameters, the center of the fold begins to close first, with the folding propagating down towards the tips. Consequently, the distal tips of the pouch are the last elements of the structure to meet and complete the fold in the simulation. This striking qualitative difference in the initiation and propagation of the folding event, while ultimately resulting in a shape similar to the endpoint at 4 h APF, suggests that this mechanism is insufficient to capture the entire folding dynamics observed experimentally.

### Reducing the cell-ECM adhesion and ECM stiffness in the center of the pouch, combined with boundary forces, initiates fold formation

To correctly replicate the temporal sequence of morphological changes occurring during eversion displayed in Fig. 1, we next focused on testing potential mechanisms that promote early contact between the proximal regions of the wing pouch. The ECM, an adhesive network of fibers surrounding cells, provides essential mechanical support and biochemical cues to the overlying epithelium^44,51^. A local disruption, specifically the detachment of the ECM from a centrally located, small group of basal cell membranes, is necessary to initiate tissue folding in the wing disc at the wandering third instar larval stage (wL3)^52^. To visualize this interface and determine the spatial extent of detachment for our model, we imaged collagen IV and actin in wL3 discs (Fig. 4A). Inspection of experimental cross-sectional images reveals a localized separation between the basal surface of the pouch epithelium and the ECM near the DVB (Fig. 4A’) in agreement with a previous study^52^.

**Figure 4.**
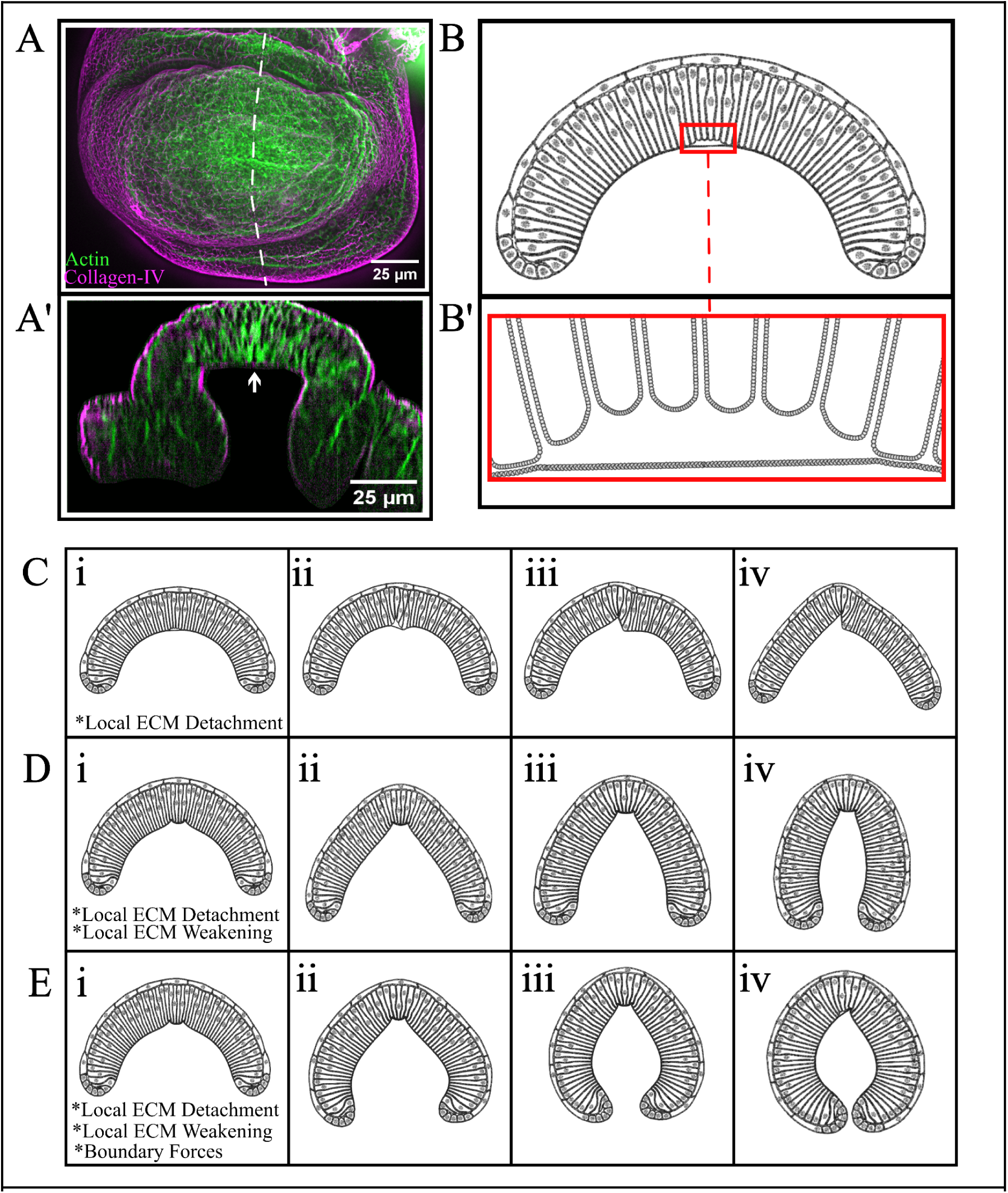
Outcomes of the detachment of the basal ECM from the basal cellular membranes of central pouch cells in model simulations. **A.** Experimental images of the wing disc pouch at wL3 with the detached basal ECM shown in the DV cross-section. **B.**

Thus, we defined the detachment domain as the 10% of pouch cells centered on the DVB. We reduced the adhesive spring force connecting the basal cell membrane to the ECM within this subdomain to model cell-ECM detachment (Fig. 4B, B’). Gradually reducing the intensity of basal cell-ECM adhesion in the central subdomain produced no obvious disruptions to the tissue’s geometric structure or equilibrium shape until the force was reduced entirely to zero (Fig. 4 and SI Fig. 3).

We observed a highly localized disruption in the basal-center of the pouch that impacts the overall tissue geometry in a free boundary setting with cell-ECM detachment acting as the sole mechanical perturbation (Fig. 4C). While capturing the overall tissue folding, the unchanged ECM rigidity present in the model scenario generates a stiff kink in the area around the DVB that disrupts the smoothness in the final tissue shape. Therefore, we hypothesized that both ECM bending and stretching stiffness also decreases concurrently with the local detachment of cells from the ECM to initiate folding.

Reducing the ECM bending and stretching stiffness by half after detachment enables the simulated tissue to deform into a smooth and symmetric bilayer structure with an open space inside near DV boundary, similar to experimental observations at 2 h APF (Fig. 4D, Fig. 1, column ii). However, this final equilibrium shape still fails to capture the proximal tip convergence as observed *in vivo*. Closer evaluation of the model and resulting final simulated shape reveals that the local ECM detachment is insufficient to drive fold formation even at the early stages of eversion (Fig. 4D). However, the combination of both applied boundary forces with local ECM detachment and weakening at the DV boundary reproduces not only proximal tip convergence but also the overall tissue morphology observed *in vivo* at 2 h APF (Fig. 4E).

Additionally, we found that perturbing the location of the detached region critically impacts the initiation of the fold formation (SI Fig. 3B-F). Further, increasing the size or number of the detached subdomains compromises the initiation of the fold formation, resulting in a tissue shape appearing more rectangular than folded (SI Fig. 3G-J). These results support the essential, but insufficient effect of localized ECM detachment for correct wing disc eversion.

### Cell rearrangements promote tissue elongation of the everting wing disc

We next examined the role of apical cell rearrangements (representing a T1 transition), which occurs during eversion (Fig. 5)^24^. The model includes a pseudo-3D component to represent cell intercalation events occurring in the two-dimensional cross-section as a net increase of cells into the DV cross-section.

**Figure 5.**
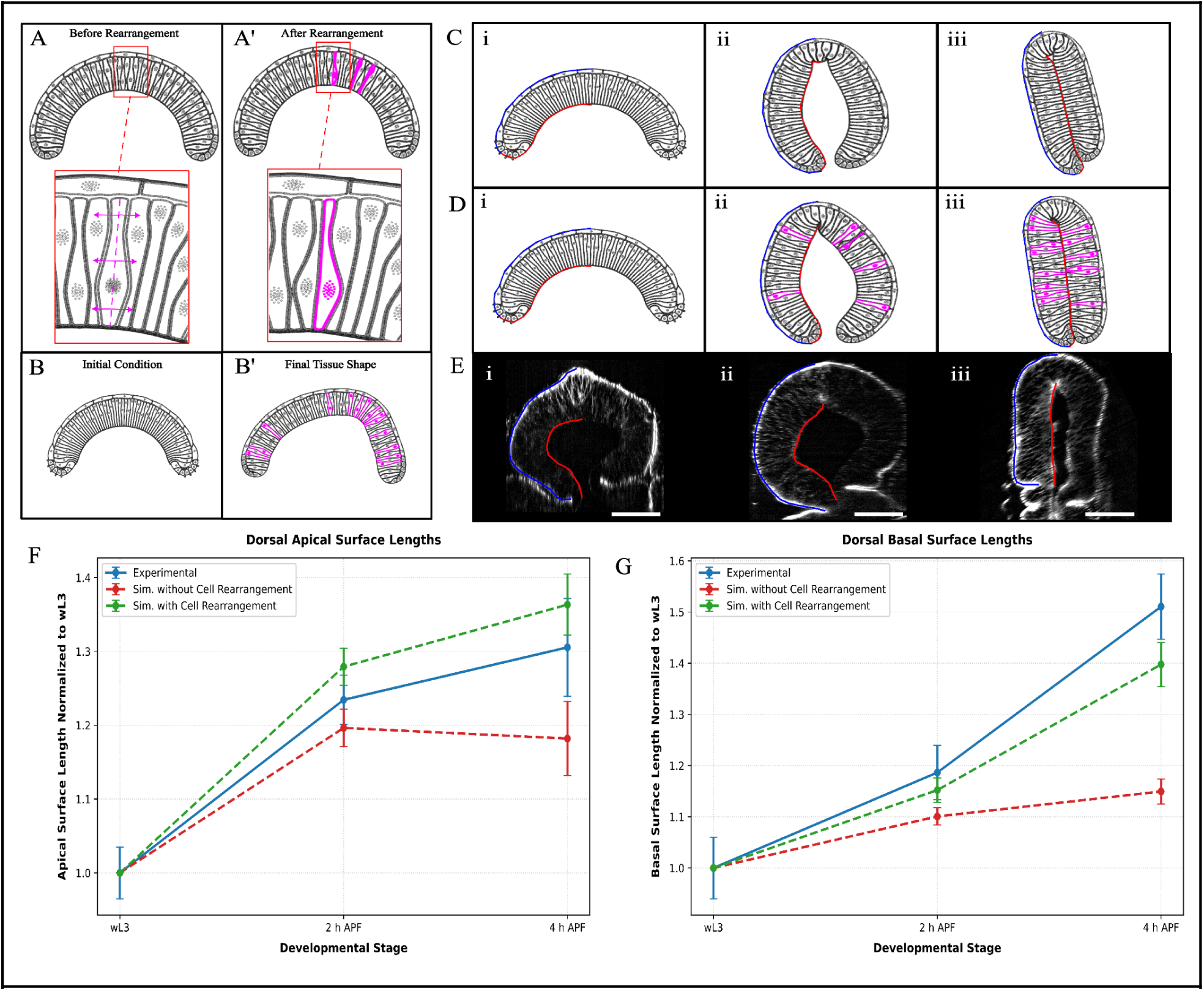
Cell rearrangement elongates the everting wing disc. **A.** Cell rearrangements are modeled by splitting cells along the vertical midline (zoom-in view) followed by volume relaxation in the plane of the cross-section. **A’.** Simulated cross-section after multiple cell intercalation events. **B.** Initial condition of tissue shape for the simulation with cell rearrangement. **B’.** Final tissue shape after doubling the number of pouch cells in the cross-sectional plane. **C.** Tissue shapes over time in a simulation combining inward boundary forces, basal ECM detachment, and basal actomyosin contraction, in the absence of cell rearrangements. **D.** Tissue shapes over time in a simulation incorporating the same components described in C. Stochastic cell rearrangements are shown in magenta. **E.** Representative experimental cross-sections showing the apical (blue) and basal (red) surface contours used for measurements within the dorsal compartments. **F.** Plot of dorsal apical lengths of experimental and simulated cross-sections at wL3, 2 h APF, and 4 h APF demonstrating that the pouch elongates along the DV axis due to cell rearrangements. Lengths are calculated by tracing the apical surface for experimental cross-sections in Fig. 1B and simulated cross-sections in 5C and 5D. A sample size of n=3 is used for each plotted line with standard deviation indicated by the vertical bars around data points. **G.** Plot of dorsal basal surface lengths of the cross-sections using the same methods as described in F.

These cell intercalation events are implemented by stochastically splitting existing pouch cells into new cells, which then exert lateral pressure on their neighbors, producing localized tissue expansion along the cross-section (Fig. 5A, A’). For instance, with cell rearrangements active, fourteen new columnar cells (∼20% increase) were intercalated by the end of the simulation (Fig. 5D). The local rates of cell rearrangements were calibrated using experimental data on tissue elongation.

Model simulations that only incorporate cell rearrangements fail to qualitatively match 4 h APF pouches (Fig. 5B’). Conversely, simulated pouches that incorporate other proposed mechanisms without cell rearrangements are sufficient to generate tissue morphology similar to experimental observations throughout the eversion process (Fig. 5C). Both simulation conditions, with and without rearrangements, qualitatively captured the intermediary and final experimental tissue shape cross-sections. Measuring the lengths from the DVB to the dorsal fold revealed that between wL3 and 2 h APF the pouch elongates by ∼25% along the apical surface and ∼20% along the basal surface (Fig. 5E, F, G). And at 4 h APF, the apical surface stabilizes, only increasing by approximately 5%, while the basal surface extends by another 30%. Overall, *in vivo* apical and basal surfaces elongate by 25-35% larger at 4 h APF compared to wL3. Both simulation scenarios generally match the experimental elongation trend, though we calculate the difference between the length of the simulated pouches and the *in vivo* pouch surfaces to measure the error, as formulated in SI Eqs. (2) and (3). With this metric, we find that activating cell rearrangements enables the pouch to more closely match the magnitude of elongation, resulting in a ∼41% decrease in error for dorsal apical and basal surface lengths across each time point (Fig. 5F, G). Furthermore, variability of elongation across different samples in experimental wing discs can be matched through tuning the rate of cell rearrangement.

### Morphogenesis of the everting wing disc is coordinated by different biophysical mechanisms interacting across both temporal and spatial scales

We refined and combined all four proposed biophysical mechanisms: patterned basal ECM detachment, delayed actomyosin contractility, cell rearrangement, and boundary forces to determine whether the system, acting as a whole, is sufficient to appropriately reproduce the developmental process. Together these results indicate that a combination of local internal mechanisms and global boundary forces transmitted from neighboring cells underlies the mechanistic robustness of wing disc eversion.

Relative to the model iteration of contractility in Fig. 3, the spatial patterning was modified across the distinct developmental stages to appropriately reflect the entire development process. We extracted spatial curvature profiles (Fig. 1D, E) and phosphorylated non-muscle Myosin II (pNMII) distributions during late third instar, wL3 (see SI Fig. 5 and SI Fig. 6), to achieve a biologically accurate initial simulated tissue shape for our comprehensive eversion simulation (Fig. 6B-i). At the onset of eversion (*T* = 0), basal ECM detachment and localized matrix weakening are initiated alongside stochastic cell rearrangements and dynamic boundary forces, while basal contractility is returned to a uniform baseline across the pouch which captures intermediate 2 h APF tissue morphology (Fig. 6B-ii).

**Figure 6.**
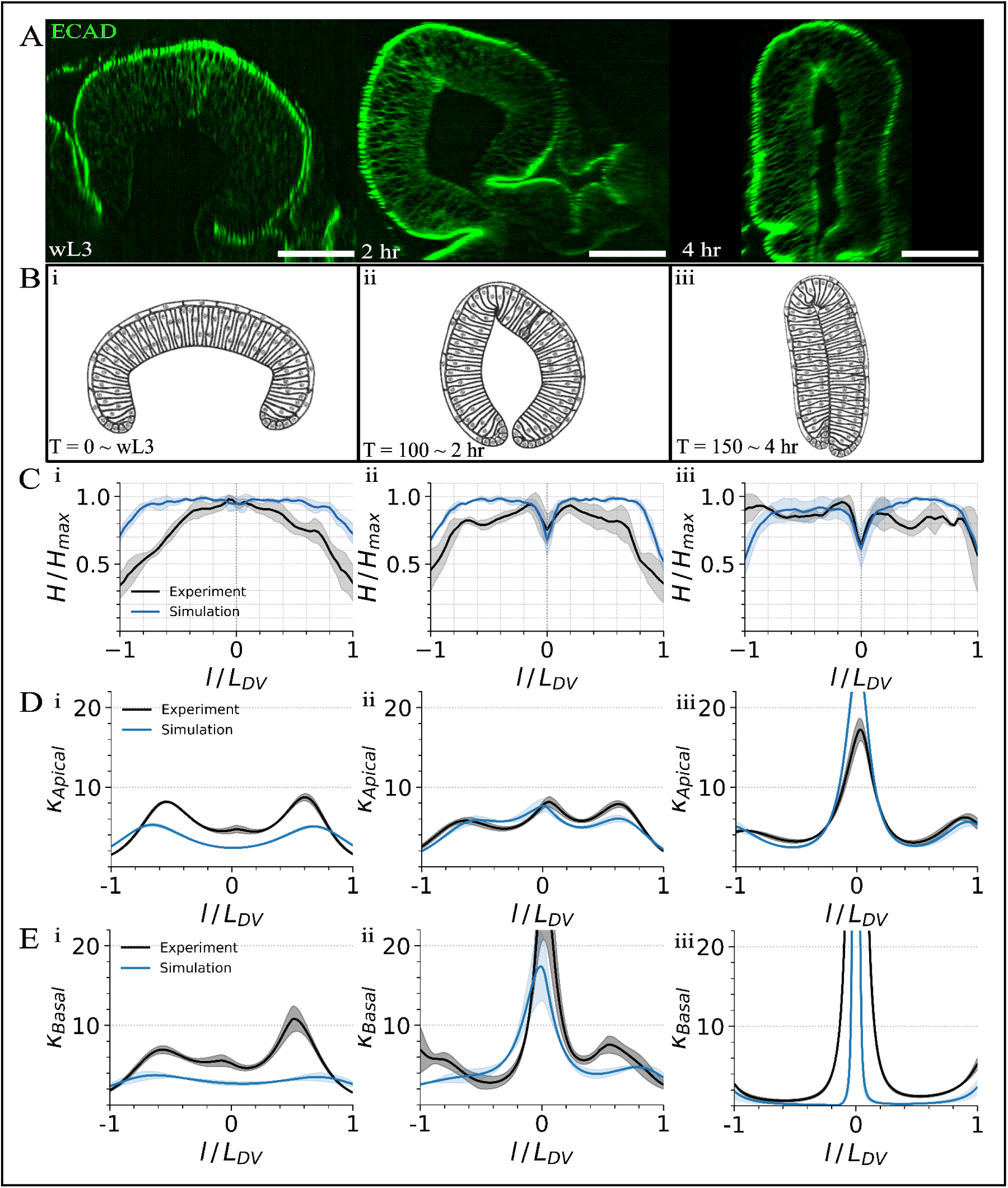
The nonlinear MORPHIOME simulation framework captures experimental tissue shapes throughout the early stages of eversion. Comparison between simulation results implementing basal ECM detachment, patterned basal actomyosin contractility, boundary forces, and cell rearrangement, and wing disc cross-sections at wL3, 2 h APF, and 4 h APF. **A.** Cross-sections of the wing disc pouch along the DV axis at different time points previously shown in Fig 1. **B.** Simulated wing discs achieved through implementing the biophysical mechanisms outlined in the study. **C.** Plots of the simulated pouch cell heights across the cross-section through wL3, 2 h APF, and 4 h APF normalized to the maximum cell height for each time point. Experimental tissue heights with the standard deviation between samples are blue and simulated tissue heights are red. **D.** Apical tissue curvature profiles for simulated cross-sections calculated with the same methods as in Fig. 1D overlaid with experimental apical curvature profiles. **E.** Basal tissue curvature profiles for simulated cross-sections calculated with the same methods as in Fig. 1E overlaid with experimental basal curvature profiles. n=6 for wL3 and 2 h APF wing discs, and n=3 for 4 h APF wing discs and for all simulation time points.

Shortly after the pouch achieves the 2 h APF intermediate tissue morphology (*T* = 100), basal actomyosin contractility is restricted to a narrow central domain corresponding to the DVB, reduced from the central third of columnar pouch cells down to the central tenth to match the localized zone of basal ECM detachment^52^.

Once activated, basal actomyosin contractility is patterned by relaxing the flanking dorsal and ventral proximal subdomains (DPR and VPR), creating a relative increase in central contractility that drives final fold closure (Fig. 6B-iii, SI Video 10). Our integrated model thus overcomes the limiting outcomes of individual morphogenetic processes and accurately reproduces *in vivo* shapes across the developmental stages of wing disc eversion.

Notably, our experimental - modeling framework demonstrates the morphogenetic robustness of wing disc eversion as the model produces similar morphogenetic outcomes when initiated with alternative dome-like shapes (SI Fig. 7). Our model also shows quantitatively similar cell height and curvature profiles (Fig. 6D, E). In sum, these qualitative and quantitative results show that the quantitative experiment - MORPHIOME framework maps the impacts of biophysical mechanisms driving eversion to morphogenetic outcomes (summarized in Fig. 7), describing key features of early *Drosophila* wing disc eversion.

**Figure 7.**
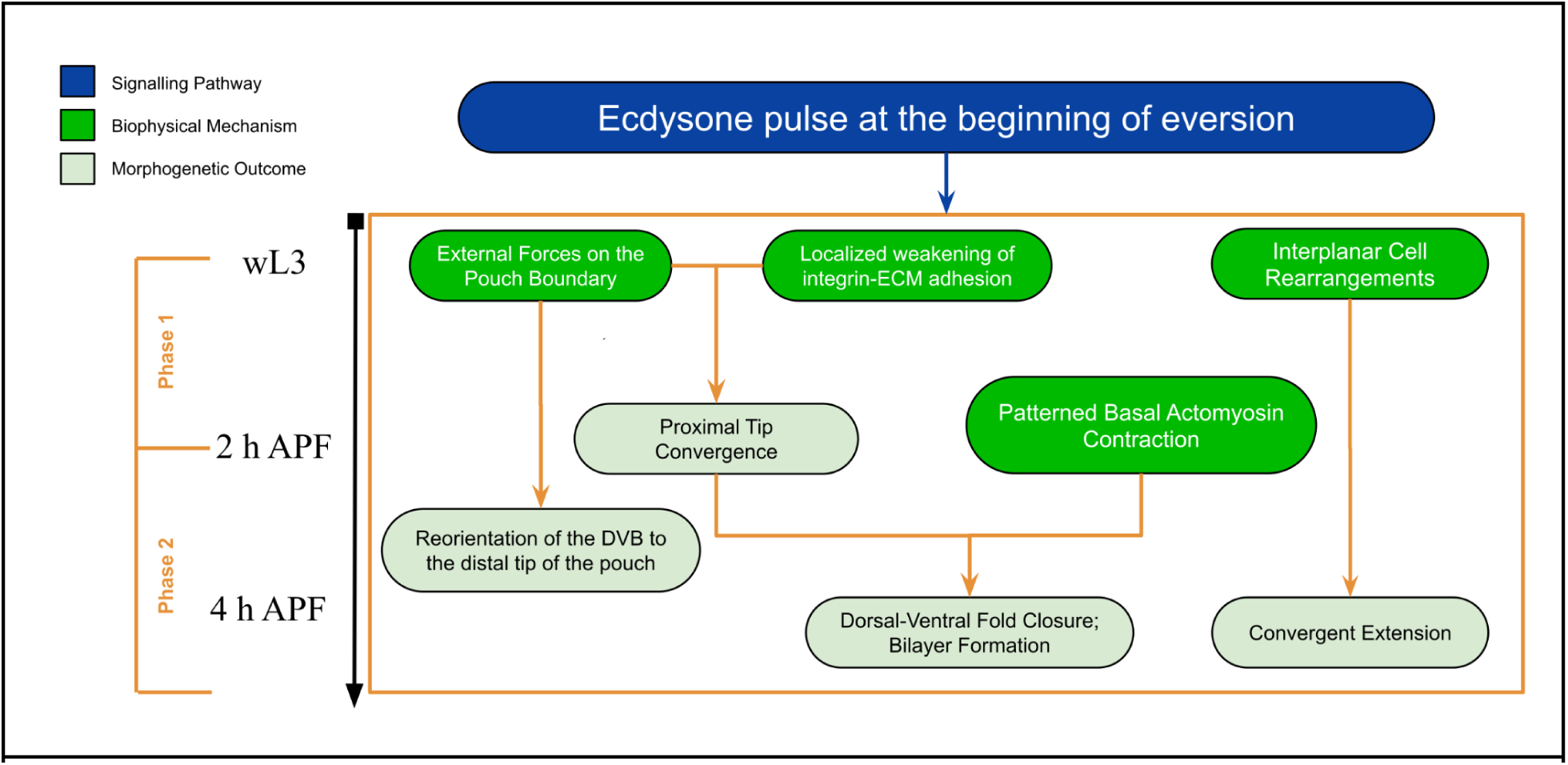
A systems-level decomposition of tissue deformation of the wing disc during eversion into separable cell-level morphogenetic processes. Phases of eversion and proposed corresponding active mechanisms of tissue deformation: (1) pouch rotation repositioning the DVB to the distal tip of the pouch, generated by boundary forces, likely from both the peripodial membrane and hinge tissue; (2) bilayer formation preceded by proximal tip convergence, from ECM detachment, boundary forces, plus basal-contractility patterning dynamics; (3) translation by convergent extension from cell rearrangements.

### Active boundary forces contribute to deformation and reorientation of the wing disc during eversion

Tracking ex vivo wing discs undergoing eversion reveals that the dorsal-proximal tip rotates about 90 degrees in-plane during eversion (SI Fig. 4A, B). Notably, none of the internal subcellular and cellular mechanisms previously tested using model simulations, were able to generate this rotational motion. To test whether external mechanical forces are necessary to drive DVB reorientation and curvature remodeling of the everting wing we applied a directional force vector, *E*_Ext_, to the proximal boundary cells, to simulate pressure from the surrounding neighboring boundary tissues. We found that the application of consistent boundary forces informed by the direction of experimentally observed unfolding external tissue dynamics were capable of matching the in-plane rotation observed in experimental samples.

A comprehensive description of the mechanical contributions of tissues surrounding the wing pouch remains incomplete. Previous studies have identified several remodeling events outside of the pouch that could influence the deformation described here^26,29,51^. Myosin-II dependent dynamics within the peripodial epithelium generate large-scale disc bending and permit expansion and retraction of the peripodial membrane during eversion^22^. However, the contribution of these forces to pouch rotation has not been quantitatively resolved in cross-sectional analysis. Similarly, recent studies have identified actomyosin and ECM-dependent remodeling within the hinge and notum, including basal ECM degradation^53^, and remodeling of the apical ECM during fold opening^54^. Nevertheless, how these neighboring tissue dynamics are mechanically coupled to pouch rotation and dorsal-ventral apposition remains unclear. Consistent with these observations, our modeling results (see Fig. 5C) demonstrate that externally imposed boundary forces are sufficient to produce pouch rotation and promote convergence of the proximal tips, which contributes to subsequent fold closure. These results support the hypothesis that forces transmitted from surrounding tissues contribute to multiple aspects of early pouch deformation, although the current model does not identify the biological origin of these forces or exclude contributions from pouch intrinsic mechanics. Together, previous studies establish that multiple regions surrounding the pouch are mechanically active during eversion, but leave unresolved whether early pouch rotation, tip convergence, and fold closure are driven primarily by forces generated within the pouch, transmitted from neighboring compartments, or coordinated between the two.

## Discussion

The morphogenesis of epithelial sheets relies on a coordinated hierarchy of processes that link subcellular mechanics with cellular dynamics and macro-scale tissue processes. In this paper, we developed a **M**ultiscale **OR**gan-level **PH**ysics-**I**nf**O**rmed **M**odeling **E**nvironment (MORPHIOME), and applied it to study organ-scale epithelial morphogenesis during *Drosophila* wing disc eversion. More specifically, we used the biologically calibrated computational modeling framework to test multiple hypothesized mechanisms of tissue folding and elongation. Wing disc eversion represents a dramatic morphogenetic transformation involving large-scale tissue deformations, spatial heterogeneities, and complex interacting forces.

First, experimental images of the wing disc pouch cross-section were analyzed to quantify morphological metrics including apical and basal membrane curvature and cell-height profiles (Fig. 1). By comparing these metrics of experimental imaging data and model simulations, we demonstrated that tissue deformation during eversion relies on the nonlinear coupling between localized sub-cellular force generators and macro-scale boundary forces (Fig. 2-6 as summarized in Fig. 7). The analysis of model simulations revealed that early wing disc eversion proceeds through two distinct morphogenetic phases:

● **Phase 1 (wL3 to 2 h APF):** Eversion is initiated by localized detachment and mechanical weakening of the basal ECM at the dorsal-ventral boundary (DVB). Concurrently, external boundary forces applied at the proximal edges drive proximal tip convergence (Fig. 4 and Fig. 6B-ii).
● **Phase 2 (2 to 4 h APF):** Spatially patterned actomyosin contractility in the basolateral region of DVB cells drives fold closure to form the wing bilayer (Fig. 3 and Fig. 6B-iii).

Across both phases, active cell rearrangements serve as a primary driver for tissue elongation in model simulations, to match the experimentally measured tissue size (Fig. 5F, G). Further, the robustness of these mechanisms were tested. First, it was shown that eversion morphogenetics were similar in simulations with different initial tissue shapes (SI Fig. 2 and SI Fig. 6). Second, when contractility is perturbed (such as in Fig. 3D), the pouch successfully folds across a range of initial tissue shapes (SI Fig. 1). However, perturbing the size and location of the detached region shifts and disrupts the precise fold location (see SI Fig. 3).

MORPHIOME lays a foundation for investigating complex epithelial morphogenetic programs and naturally translates to studies of organogenesis to identify biophysical mechanisms of congenital defects and as a design tool for synthetic multicellular engineering. Comprehensive exploration of the full multidimensional parameter space requires further developments to accelerate simulations and reduce computational expense. Additionally, future efforts are focused on developing an explicit 3D organ-level model at the subcellular element resolution^55,56^. Additional envisioned extensions include explicitly coupling of the mechanical model with both biochemical signaling and gene regulatory networks simulations^57^. Currently, the mechanical parameters are coarse-grained representations of downstream outputs of regulatory networks. Developing a predictive mechanochemical extension for MORPHIOME will enable systematic analysis of how developmental programs dynamically generate, coordinate, and adapt those forces during morphogenesis.

In summary, MORPHIOME is capable of providing key insights into the biophysical principles governing *Drosophila* wing disc morphogenesis and, more broadly, organogenesis as a whole. In particular, obtained results demonstrate that isolated, individual mechanical drivers fail to capture the full developmental process. Rather, only the dynamic, multiscale integration of ECM remodeling and detachment, delayed contractility patterning, cell rearrangements, and external tissue forces fully recapitulates the spatiotemporal changes in cell heights and curvature profiles observed *in vivo* (Fig. 6). Second, we infer how localized micro-scale force generators drive tissue-scale events. Third, coordination of multiple mechanobiological mechanisms across scales ensures organized, robust morphogenesis. Finally, these insights are relevant to the multiscale biomechanical coordination occurring across many other biological contexts ranging from ventral furrow formation and gastrulation, germband extension, dorsal closure and epidermal furrow formation in the fly to key developmental processes such as neural tube folding, optic cup formation, cortical folding in the brain in mammalian systems. As such, MORPHIOME provides an extensible and transferable environment enabling investigation of morphogenesis at multiple layers and scales^58–61^.

## Methods

### Computational formulation of MORPHIOME

**M**ultiscale **OR**gan-level **PH**ysics-**I**nf**O**rmed **M**odeling **E**nvironment (MORPHIOME) includes sets of nodes representing cell membranes, nuclei, and the surrounding extracellular matrix (ECM) (Fig. 2). These nodes are connected with one another via different types of springs. Their interactions are represented in the model through energy potentials. Node dynamics are described by a large system of coupled nonlinear Langevin ordinary differential equations evolving under overdamped viscous conditions since they represent cells in a tissue with limited fluctuations. Morphogenetic forces are calculated and balanced via nonlinear energy potentials, including Morse potentials, controlling volume exclusion and determining nuclear sizing, spring elasticity for membrane dynamics, and Lagrange multipliers for cellular volume conservation.

Beyond structural details, the MORPHIOME incorporates key biological processes, such as cell division and rearrangement, and mechanical forces including internal cellular pressure, nuclear positioning, actomyosin cytoskeletal activity, and adhesivity (Fig. 2D, E, E’). Previous work using MORPHIOME were calibrated with experimental data and have yielded key insights into the regulation of growth and tissue mechanics during the preceding larval stage of development; this demonstrates the importance of the ECM in maintaining tissue structure, the role of actomyosin forces in defining the basal curvature, and how rates of cell division impact overall tissue morphology^18,46,62^.

To study the biomechanical mechanisms driving the drastic changes in tissue morphology observed during wing disc eversion, we developed the MORPHIOME calibrated from experimental data. The MORPHIOME uses subcellular element (SCE) computational framework to simulate a cross-section of the *Drosophila* wing imaginal disc along the DV axis during the developmental process of eversion (Fig. 2)^18,46,62^. The SCE modeling framework is well established in the field of biophysics to study the impacts and emergent tissue level behavior produced from local cell and subcellular mechanical perturbations^43,63,64^. By coupling sub-cellular level dynamics with active boundary conditions that represent the dynamic constraints of the external tissue connected at the distal tips of the pouch, we report the novel MORPHIOME modeling framework capturing wing disc development from the late third instar, wL3, through the early stages of pupal development, 4 h APF.

The MORPHIOME presented in this paper represents the wing disc pouch cross-section with three distinct node types: cell membrane nodes, internal nucleus nodes, and extracellular matrix (ECM) nodes. Each node’s position and interaction with other nodes are governed by specific energy functions, and are evolved by Langevin small-particle dynamics within an overdamped environment; this is consistent with the high-viscosity environments of biological systems^39,43,44^. To accommodate this overdamping, a constant damping term calibrated from previous studies is applied to each equation of motion^46^. The modeling domain is initially populated with approximately 37,000 nodes, comprising roughly 30,000 membrane nodes, 5,000 ECM nodes, and 2,000 internal nucleus nodes. This high-resolution node distribution allows for a detailed coarse-grained representation of the molecular landscape, where individual nodes are grouped together to construct the larger cellular level structures of the pouch. The dynamics of these nodes are governed by Equations (1)-(3) as follows, respectively.

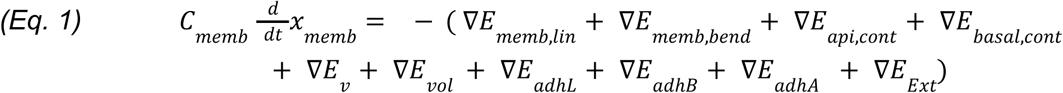

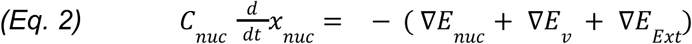

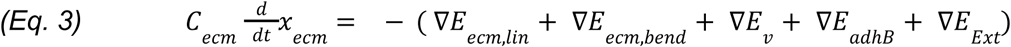

The right-hand sides of Equations (1)-(3) represent the net forces acting on the respective node types at a given time step. They are represented by the sum of negative gradients of the energy potentials. *C_memb_*,*C_nuc_*, *C_ecm_* are the damping coefficients for each node according to node type to represent the overdamped dynamics, replicating the viscous conditions surrounding the wing disc organ. Systems of equations (1)-(3) are solved computationally via gpu parallelization^65^. The energy potentials that define cellular mechanical properties and cell-cell and cell-ECM interactions are listed in Table 1, along with their biophysical meaning and specific formulation. Schematic representations of the nodes connected with each other representing the cell and tissue level structures are demonstrated in Fig. 2D with the Paraview visualization in Fig. 2E and E’^66^.

**Table 1.** Energy Potentials.

| Energy Potential | Type | Biophysical Meaning | Equation Representation |
|---|---|---|---|
| *E*<sub>*memb,lin*</sub> | Linear Spring | Elasticity of the cell membrane | $E_{linear} = \frac{1}{2}k_{linear}(\|\boldsymbol{x}_i - \boldsymbol{x}_j\| - L_0)^2$ |
| *E*<sub>*memb,bend*</sub> | Bending Spring | Bending elasticity of the cell membrane | $E_{memb,bend} = \frac{1}{2}k_{memb,bend}(\theta_{memb} - \theta_0)^2$ |
| *E*<sub>*api,cont*</sub> | Linear Spring | Apical constriction of actomyosin inside the cells above the cell nucleus | $E_{linear} = \frac{1}{2}k_{linear}(\|\boldsymbol{x}_i - \boldsymbol{x}_j\| - L_0)^2$ |
| *E*<sub>*basal,cont*</sub> | Linear Spring | Basal constriction of actomyosin inside the cells below the cell nucleus | $E_{linear} = \frac{1}{2}k_{linear}(\|\boldsymbol{x}_i - \boldsymbol{x}_j\| - L_0)^2$ |
| *E*<sub>*v*</sub> | Morse Potential | Volume Exclusion | $E_{morse} = U_{morse}exp(\frac{-\|\boldsymbol{x}_i-\boldsymbol{x}_j\|}{\xi_{morse}}) - W_{morse}exp(\frac{-\|\boldsymbol{x}_i-\boldsymbol{x}_j\|}{\gamma_{morse}})$ |
| *E*<sub>*vol*</sub> | Lagrange Multiplier | Volume conservation of the cell cytoplasm | $E_{Lagr} = k_{vol}(\Omega - \Omega_0)^2$ |
| *E*<sub>*adhL*</sub> | Linear Spring | Adhesion of cells to other cells via E-cadherin | $E_{linear} = \frac{1}{2}k_{linear}(\|\boldsymbol{x}_i - \boldsymbol{x}_j\| - L_0)^2$ |
| *E*<sub>*adhB*</sub> | Linear Spring | Adhesion of the ECM to cells mediated by Integrin | $E_{linear} = \frac{1}{2}k_{linear}(\|\boldsymbol{x}_i - \boldsymbol{x}_j\| - L_0)^2$ |
| *E*<sub>*adhA*</sub> | Linear Spring | Adhesion of the columnar pouch cells to apical peripodial cells | $E_{linear} = \frac{1}{2}k_{linear}(\|\boldsymbol{x}_i - \boldsymbol{x}_j\| - L_0)^2$ |
| *E*<sub>*ecm,lin*</sub> | Linear Spring | Elasticity of the Extracellular Matrix (ECM) | $E_{linear} = \frac{1}{2}k_{linear}(\|\boldsymbol{x}_i - \boldsymbol{x}_j\| - L_0)^2$ |
| *E*<sub>*ecm,bend*</sub> | Bending Spring | Bending elasticity of the Extracellular Matrix (ECM) | $E_{ecm,bend} = \frac{1}{2}k_{memb,bend}(1 - (cos(\theta_{memb} - \theta_0))$ |
| *E*<sub>*nuc*</sub> | Morse Potential | Size of the cell nucleus | $E_{morse} = U_{morse}exp(\frac{-\|\boldsymbol{x}_i-\boldsymbol{x}_j\|}{\xi_{morse}}) - W_{morse}exp(\frac{-\|\boldsymbol{x}_i-\boldsymbol{x}_j\|}{\gamma_{morse}})$ |
| *E*<sub>*ext*</sub> | Force Vector | Boundary Force applied by tissue outside the pouch | $E_{Ext} = F(t) * \hat{d}$ |

As described previously, nodes are grouped together to represent the distinct cell types in the wing disc pouch including columnar epithelial pouch cells, squamous peripodial cells, cuboidal boundary cells, as well as the surrounding ECM layers. While connective tissues extend beyond the pouch boundaries *in vivo* along the DV axis, our two-dimensional cross-sectional model is truncated at the proximal tips with cuboidal boundary cells. The inclusion of these cells ensures model stability and mechanical tissue integrity. We evaluated several boundary condition formulations to represent the pressure applied by the hinge, notum, and other external tissues connected to the proximal tips settling on a dynamic application of force vectors that mimics the non-local mechanical constraints observed across progressive stages of disc eversion.

Collectively, the combination of different types of cells constitute the integrated tissue-level representation of the wing disc pouch cross-section. Model baseline parameters were calibrated to ensure the initial equilibrium state recapitulates the characteristic curved, dome-like morphology observed at the onset of pupariation, wL3. By locally perturbing or patterning the mechanical properties at the node or cellular levels, such as introducing patterned basal cell contractility, we drive the system away from this initial steady state. The resulting transition toward a new global equilibrium reveals emergent morphogenetic behaviors. This framework allows us to systematically test hypothesized biological mechanisms, either in isolation or in tandem, identifying the necessary and sufficient biophysical conditions required for the wild-type wing disc to execute the complex geometric transformations observed during eversion.

### Sensitivity analysis

Due to detailed model representation of cells and ECM in the wing disc pouch cross-section, model simulations have a significant computational cost. A complete simulation runs with a time step size of 0.002 arbitrary units (AU) to maintain the numerical stability for a time of 50,000 totaling to 25,000,000 time steps in a complete simulation. An output file is printed after every 50,000 time steps resulting in 500 output files for each piece of data in a simulation. Given the sheer number of nonlinear force interactions calculated for each time step, Eqs. (1)-(3) are solved in tandem via GPU parallelization. Despite being updated for some of the newest versions of CUDA compatible with NVIDIA A100 GPU processors, a single simulation takes on average at least four and a half days (∼110 hours) to complete, making extensive sensitivity analysis of model parameter values and initial conditions in realistic time periods a challenge.

Instead, we focus on assessing how varying initial conditions affect the resulting morphogenetic outcomes associated with each proposed mechanism. For example, simulations testing basal ECM detachment were run with a variety of different rates of cell-ECM adhesivity before concluding that the most biologically accurate representation was to decrease the cell-ECM adhesion to zero in the central subdomain. We also choose the following parameters including sizes of the central subdomain, ratios of patterned basal actomyosin contractility, rates of cell rearrangement, and boundary force applications to perform sensitivity analysis. This is because these parameters directly impact wing disc eversion. The best perceived results from individual sensitivity analysis were combined in joint tests with other mechanisms. Additional joint mechanism sensitivity analysis was run with strategically chosen parameter values similar to Latin-Hypercube Sampling^46,67,68^.

### Initial tissue shape and structure generation

The pseudo-3D MORPHIOME initializes simulations using pre-configured datasets calibrated to match the experimentally observed morphology of the wing disc. While previous iterations of this framework primarily utilized a single initial condition from which to make mechanical perturbations, the current study employs a sequential initialization protocol where results from preceding simulations may serve as the starting configurations for subsequent simulations, matching up to experimental data at specific time points. For example, a MORPHIOME simulation with contractility patterning informed by pNMII spatial profiles taken sometime during the late third instar period generated a tissue shape similar to wL3 (SI Fig. 6). The simulation data at a suitable time step was extracted and then used as the initial configuration for the simulation in Fig. 6B. To facilitate this temporal continuity, we developed a custom MATLAB-based data parser (see Supplementary Resource folder) that converts simulation output files at a desired time step, containing node and cell profiles, into valid initial condition configuration files. This pipeline allows for the high-fidelity transfer of mechanical states across different developmental windows, providing a robust foundation for testing long-term morphogenetic transformations.

## Experimental Methods

### *Drosophila* stocks and husbandry

*Drosophila melanogaster* stocks were maintained on Fly Food B (LabExpress) at 25 °C under a 12 h light/dark cycle. Adult flies were transferred to fresh food vials daily to minimize larval overcrowding.

Fluorescent reporter lines included *E-cad::GFP; pnr-GAL4, Sqh::mCherry*, provided by the Yanlan Mao laboratory, and *y w; Vkg-GFP* (laboratory stock #161). E-cad::GFP corresponds to the endogenous *shg^*GFP knock-in previously described ^69^, whereas Sqh::mCherry is a functional genomic reporter expressed under native sqh regulatory sequences^19^; the combined *E-cad::GFP; pnr-GAL4, Sqh::mCherry* line has been reported previously^70^. The combined reporter line was used to visualize E-cadherin and non-muscle myosin II, whereas Vkg-GFP was used to visualize Collagen IV. The Vkg-GFP reporter is a GFP protein-trap line from the Carnegie/FlyTrap collection^71^. Experimental larvae were collected directly from these stocks. BDSC #84340 was crossed to BDSC #60584, and progeny of the appropriate genotype were selected for live imaging.

Male wandering third-instar (wL3) larvae were used for experiments. For live imaging, wing imaginal discs were dissected directly from wL3 larvae and placed into ex vivo culture. For fixed developmental time-course experiments, wL3 larvae were allowed to pupariate and aged to the indicated developmental stage. Zero hours after puparium formation (0 h APF) was defined as the white prepupal stage, identified by anterior spiracle eversion and cessation of locomotion.

### Fixed tissue preparation and immunohistochemistry

Male wandering third-instar larvae were collected and staged as described above. Prior to dissection, larvae were surface sterilized by sequential transfer through phosphate-buffered saline (PBS; Sigma-Aldrich, Cat. P5368), 70% isopropanol, and PBS. Animals were partially dissected in PBS to expose the wing imaginal discs while leaving them attached to the carcass, then fixed in 4% methanol-free formaldehyde in PBS for 20–30 min at room temperature. Samples were washed three times for 10–15 min in 0.03% PBT [PBS containing 0.03% Triton X-100(Sigma-Aldrich, Cat. T9284)] with gentle rocking.

Samples were blocked in 5% normal goat serum (NGS; Jackson ImmunoResearch, Cat. 005-000-121) for 30–60 min at room temperature. For *E-cad::GFP; pnr-GAL4, Sqh::mCherry* samples, phospho-myosin II was labeled by overnight incubation at 4 °C with rabbit anti-phospho-Myosin Light Chain 2 (Ser19) antibody (Cell Signaling Technology, Cat. 3671S; 1:50), followed by goat anti-rabbit IgG (H+L) Alexa Fluor 647 secondary antibody (Invitrogen, Cat. A-21244; 1:500) and DAPI (Sigma-Aldrich, Cat. D9542; 1:1000) for at least 2 h at room temperature.

For *vkg-GFP* samples, the same fixation, washing, and blocking conditions were used, followed by incubation with DAPI (1:1000) and Alexa Fluor Plus 750 phalloidin (Invitrogen; Cat. A30105 1:500) for at least 2 h at room temperature. Samples were then washed in 0.03% PBT, with the final wash continued overnight at 4 °C, and protected from light following fluorescent staining.

Wing imaginal discs were isolated and mounted in VECTASHIELD Antifade Mounting Medium (Vector Laboratories, Cat. H-1000-10). Mounting chambers were constructed on 24 × 60 mm #1.5 cover glass (VWR, Cat. 16004-312) using two layers of double-sided tape as spacers. Approximately 30 µL of mounting medium was added, and discs were oriented with the apical surface facing the base coverslip. A 22 × 22 mm #1.5 coverslip (VWR, Cat. 16004-302) was placed over the chamber and sealed with clear nail polish prior to imaging.

### Ex vivo culture and live imaging

Ex vivo culture medium was prepared based on previously described conditions for live imaging of Drosophila wing imaginal discs^6,22^. Shields and Sang M3 insect medium (HiMedia Laboratories, Cat. IM004) was prepared from powder at 39.8 g/L and supplemented with 0.5 g/L potassium bicarbonate (KHCO₃; Sigma-Aldrich, Cat. 60339), 2% fetal bovine serum (FBS; Gibco/Thermo Fisher Scientific, Cat. A5256701), and 0.5% penicillin-streptomycin (Sigma-Aldrich, Cat. P4333). The medium was adjusted to pH 6.7 with KOH, filter sterilized through a 0.22-µm filter, and stored at 4 °C.

20-Hydroxyecdysone (20E) (Sigma-Aldrich, Cat. H5142) was prepared as a 1 mg/mL stock in ethanol and stored at −80 °C. On the day of imaging, a 10 µg/mL intermediate solution was prepared in ethanol and diluted into culture medium to a final concentration of 0.1 µg/mL, resulting in a final ethanol concentration of 1% (v/v).

WL3 male larvae were collected and surface sterilized as described above. Wing imaginal discs were fully dissected and mounted using a modified version of the simple imaging chamber previously described^72^. Discs were oriented with their apical surfaces facing the glass bottom dish (Matsinami Glass, VWR Cat. 10810-056) in 20E-supplemented medium and gently restrained using a porous cell-culture insert (Millipore, Cat. PI8P01250) modified by removal of its support legs; additional culture medium was added within the insert and overlaid with light mineral oil (Sigma-Aldrich, Cat. M5310) to limit evaporation.

### Confocal microscopy and image acquisition

Fixed and live samples were imaged on a Nikon Ti2-E inverted microscope equipped with a Crest X-Light V3 spinning-disk confocal unit, Kinetix 22 back-illuminated sCMOS camera, and Lumencor Celesta light source. Image acquisition was controlled using NIS-Elements Advanced Research software. Unless otherwise noted, imaging was performed in standard spinning-disk confocal mode using a Nikon 40× Plan Fluor DIC H/N2 oil-immersion objective (NA 1.30, working distance 0.20 mm).

Fixed *E-cad::GFP; pnr-GAL4, Sqh::mCherry* and *vkg-GFP* samples were imaged as z-stacks with a 0.3-µm step size. Fluorescence channels were selected according to the reporters and stains described above. Acquisition settings were held constant across samples within experiments used for quantitative fluorescence-intensity comparisons.

For live imaging, E-cad::GFP; pnr-GAL4, Sqh::mCherry wing discs were imaged using the E-cadherin-GFP signal. Z-stacks were acquired at 1-µm intervals every 10 min for the duration of the time-lapse experiment. Samples were maintained at approximately 25 °C using a Tokai Hit stage-top incubation system and objective warming jacket. No supplemental gas flow was used.

## Supporting information

Code and data files for quantification metrics

## Acknowledgments

We thank S. Cini, D. Gazzo, and D. Geresu for discussions on the project and help reviewing the manuscript before submission. This work was supported by the Collaborative National Science Foundation grant NSF2424826 and NSF2424827 to all co-authors. J.Z. was supported in part by the National Institutes of Health under award number R35GM156615, the EMBRIO Institute (contract #2120200, a National Science Foundation Biology Integration Institute), and the Bioengineering & Life Sciences (BELS) Initiative (Notre Dame Morphogenetic Bioengineering Initiative) at the University of Notre Dame. E.W. and W.C. were supported in part by the National Science Foundation grant DMS2338630.

## Author Contributions

E.N. Prepared the original manuscript, and E.N. and B.S. led manuscript writing and revisions.

N.S. and E.W. contributed to writing and finalizing the manuscript.

B.S. designed and performed experiments and developed experimental and imaging methodologies

E.N. and Z.X. substantially extended the model and its computational implementation.

E.N. performed simulations.

B.S. E.N. N.S. and E.W. analyzed experimental data, with B.S. leading development of experimental quantification methodology.

W.C. Z.X. contributed to experimental and simulation data analysis, and edited the manuscript.

J.Z. and M.A. conceived, designed, analyzed, and interpreted results, supervised this study, and contributed to writing and editing the manuscript.

## Acknowledgement of AI Usage

The authors acknowledge that during the preparation of this manuscript they used Google Gemini, Claude, and ChatGPT to assist with data analysis and generating code for data visualization. Additionally, Gemini was used to help with text refinement and formatting. The authors critically reviewed, verified, and edited all generated outputs, and take full responsibility for the factual integrity, computational validity, and scientific conclusions presented in this work.

## Data availability

Due to data size constraints, the 3D confocal microscopy data are available on request from the corresponding authors. This includes all of the representative source data and analysis files for experiments within this manuscript. Source data are provided with this paper.

## Code availability

An open-source C + + and CUDA implementation of the MORPHIOME is available on GitHub (https://github.com/EthanNowaski/Episcale_CrossSectionalView). Source data are provided with this paper.

## Supplementary Information for

### Robustness of fold formation to variations in initial tissue shape

To assess the robustness of fold formation, we performed simulations starting from different initial tissue geometries while applying the same spatially patterned basal actomyosin contractility. Robust fold formation was observed across all initial tissue configurations, resulting in similar final tissue shapes despite differences in their initial geometries.

**SI Figure 1.**
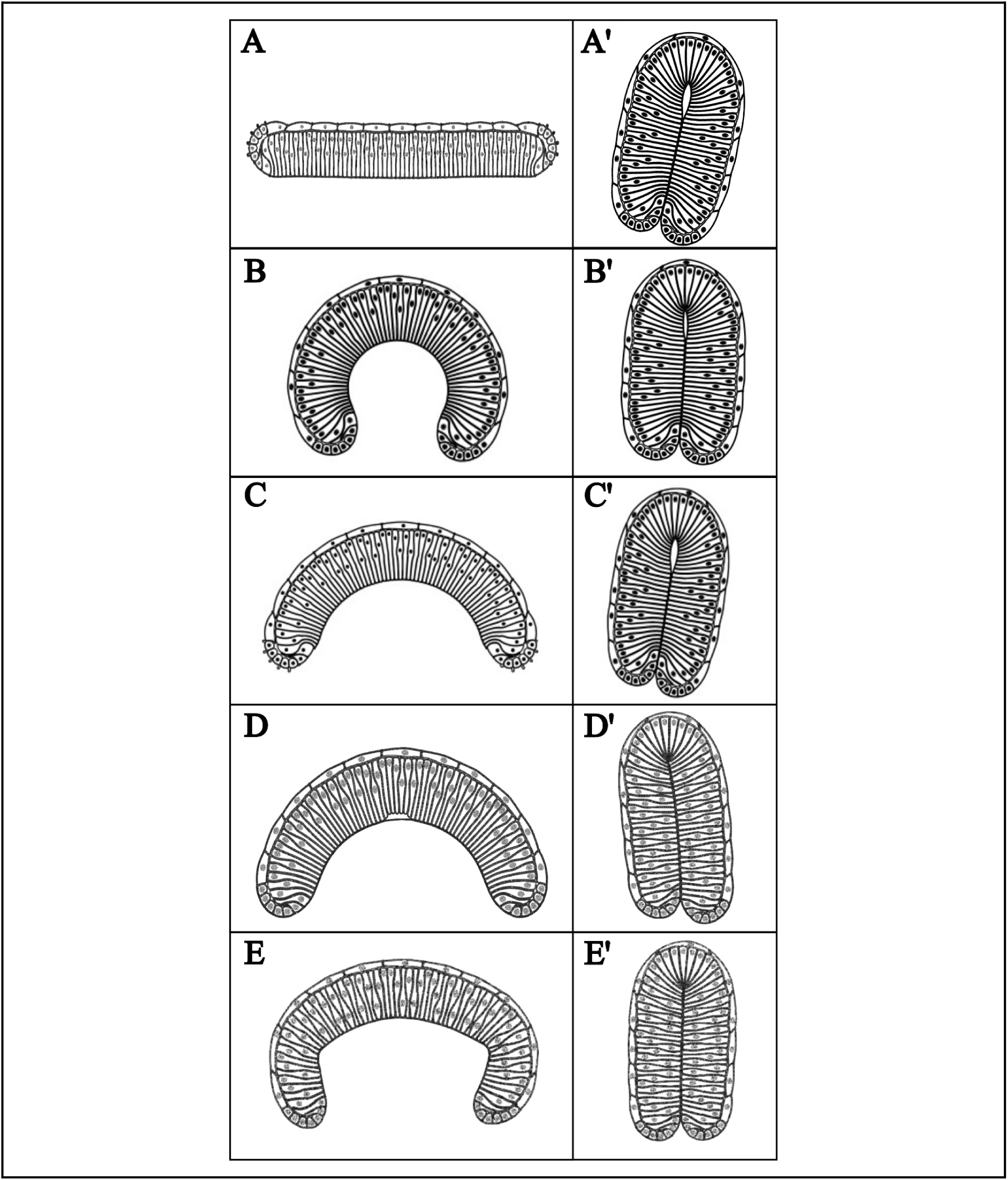
Patterned basal actomyosin contractility can generate fold formation from different initial tissue shapes. The basal actomyosin force is applied locally to the central pouch subdomain. For A-D the central pouch subdomain is defined as the central 21 cells in the pouch (∼33% of pouch cells). For E-G the central pouch subdomain is defined as the central 5 cells in the pouch (∼10% of pouch cells) corresponding to the DVB. **A**. Initial and final simulation results with a flat initial tissue shape. **B**. Initial and final simulation results with an overly curved initial tissue shape. **C.** Initial and final simulation results with the standard dome-like, curved initial tissue shape. **D**. Initial and final simulation results with the standard dome-like, curved initial tissue shape also with local basal cell-ECM detachment applied as an initial condition. **E.** Initial and final simulation results with the wL3 calibrated initial tissue shape from SI Fig. 6.

### Effects of basal actomyosin contractility and basal membrane contraction on final tissue morphology

We performed simulations with varying strengths of spatially patterned basal actomyosin contractility and basal membrane contraction, both independently and in combination, to investigate their effects on final tissue morphology. Within a moderate range of basal actomyosin contractility, increasing contractile strength resulted in more pronounced tissue bending. However, basal actomyosin contractility alone was insufficient to bring the two tissue ends together and produce the bilayer structure observed experimentally. Excessively strong basal actomyosin contractility instead resulted in a nearly flat tissue morphology, with a stiffened central region associated with elevated contractile forces. In contrast, increasing basal membrane contraction promoted more extensive tissue folding, ultimately leading to the formation of a bilayer structure resembling the experimentally observed morphology.

**SI Figure 2.**
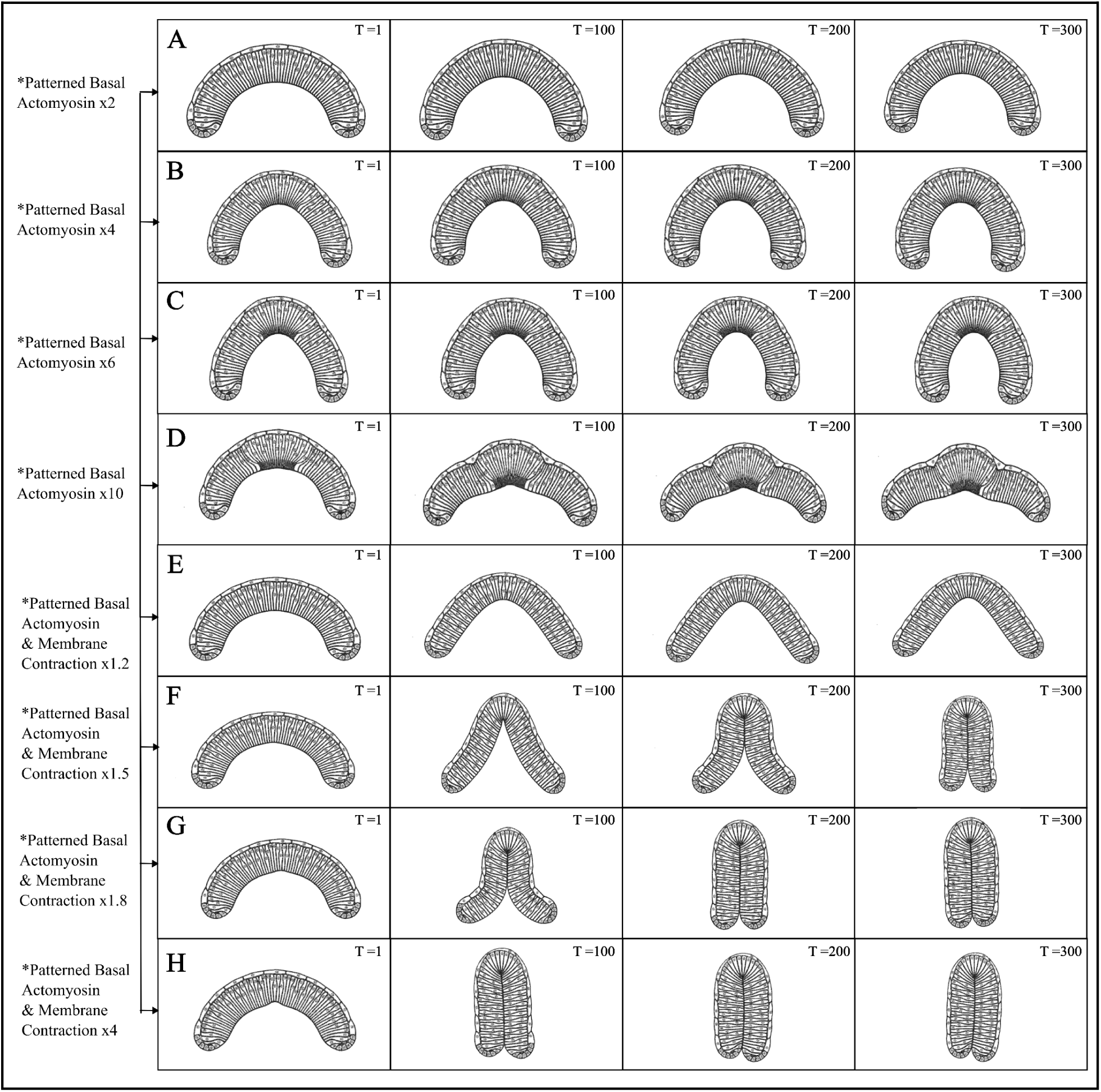
Additional cases of varying levels of contractility in the central pouch subdomain. Simulated tissue shapes and structures at different time points with basal actomyosin and basal membrane contraction patterned at different strengths in the central pouch subdomain. For A-D the central pouch subdomain is defined as the central 21 cells in the pouch (∼33% of pouch cells). For E-G the central pouch subdomain is defined as the central 5 cells in the pouch (∼10% of pouch cells) corresponding to the DVB. **A.** Simulation result with central basolateral actomyosin spring force x2. **B.** Simulation result with central basolateral actomyosin spring force x4. **C.** Simulation result with central basolateral actomyosin spring force x6. **D.** Simulation result with central basal actomyosin spring force x10. **E.** Simulation result with central basolateral actomyosin and basal membrane spring contractility x1.2. **F.** Simulation result with central basolateral actomyosin and basal membrane spring contractility x1.5. **G.** Simulation result with central basolateral actomyosin and basal membrane spring contractility x1.8. **H.** Simulation result with central basolateral actomyosin and basal membrane spring contractility x4.0.

### Effects of the location and spatial extent of basal cell-ECM detachment on final tissue morphology

We performed simulations with basal cell-ECM detachment occurring at different tissue locations and spanning over different numbers of cells. In all cases, tissue folding consistently occurred at the site of basal cell-ECM detachment, regardless of its location. Furthermore, the width of the resulting fold was determined by the spatial extent of the detachment region, with a larger number of detached cells producing a wider fold.

**SI Figure 3.**
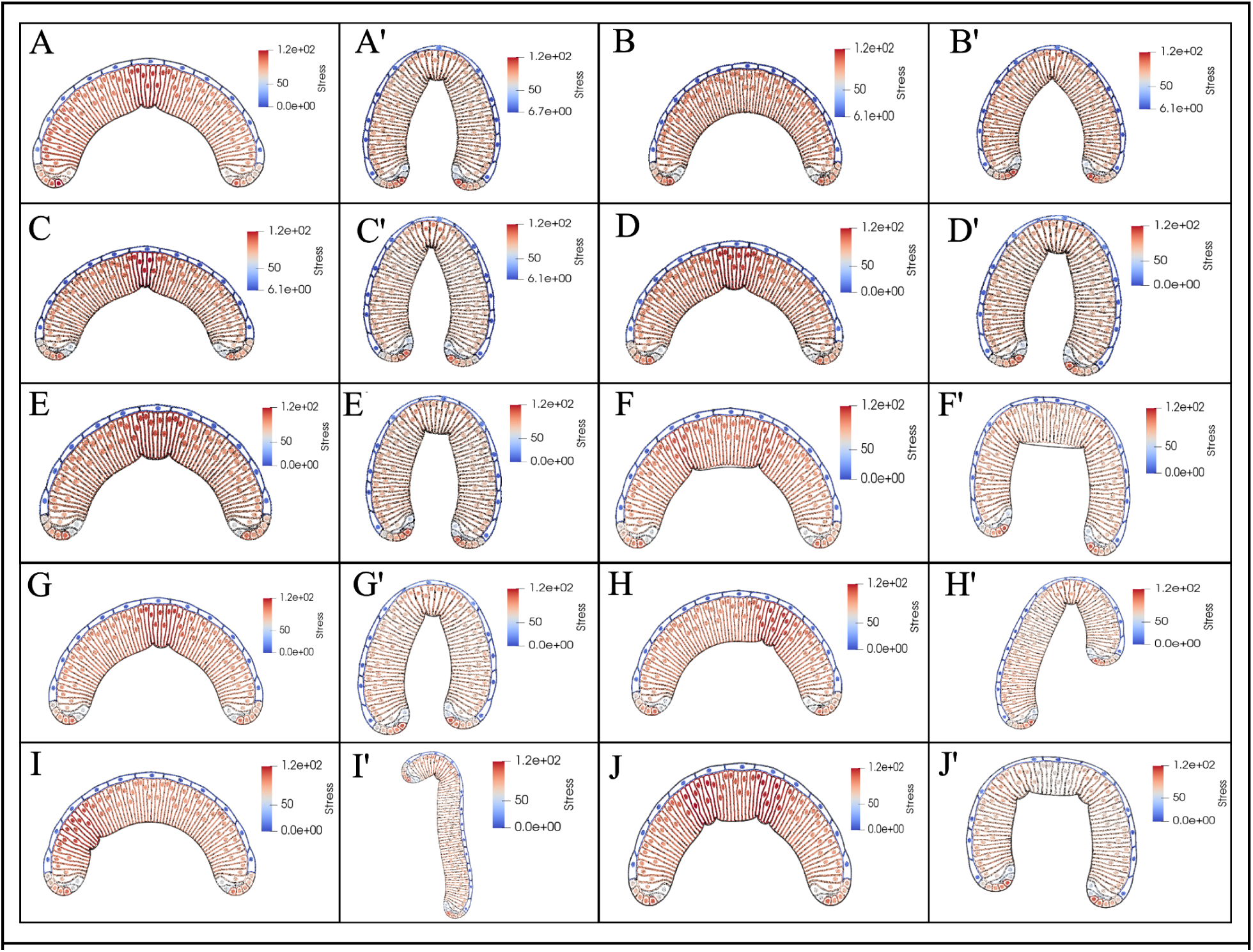
Perturbing the size and location of the detached basal ECM region in Simulations. Initial and final SCE simulation outputs with perturbations made to the locations and size of the region with cell-ECM detachment along the basal surface. Basal ECM linear and bending spring forces are also reduced by half in these regions. The color encodes the total stress magnitude (in nN/μm). **A, A’** Control simulation with the central subdomain of the detached basal ECM region involving 5 cells located in the center of the simulated pouch tissue. **B, B’** Simulation with the central subdomain of the detached basal ECM region involving 1 cell located in the center of the simulated pouch tissue. **C, C’** Simulation with 3 cells in the central detached ECM region. **D, D’** Simulation with 7 cells in the central detached ECM region. **E, E’** Simulation with 9 cells in the central detached ECM region. **F, F’** Simulation with 18 cells in the central detached ECM region. **G, G’** Simulation with the subdomain of detached basal cell-ECM region shifted over to the right 5 cells. **H, H’** Subdomain of detached basal cell-ECM region shifted to the right 10 cells. **I, I’** Subdomain of detached basal cell-ECM region shifted to the left 20 cells. **J, J’** Number of subdomains with detached basal cell-ECM increased to 2. Each region is made up of 5 cells and is shifted 10 cells to the left and right of the center, respectively.

**SI Figure 4.**
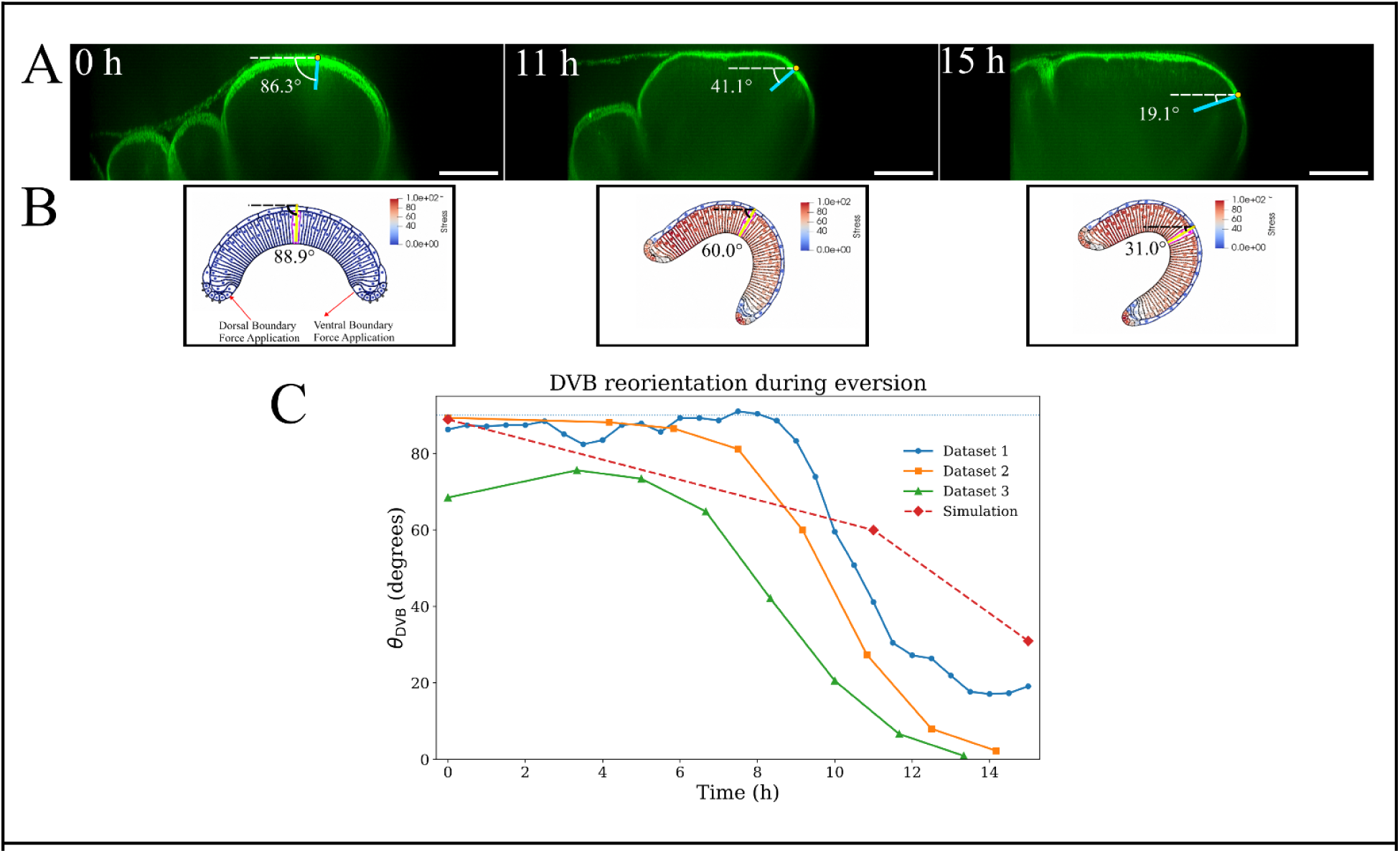
Dorsal-Ventral boundary reorientation during eversion is recapitulated by proximal boundary forces in the model. **A.** Representative E-cad::GFP cross sectional views of a wing disc pouch cultured ex vivo at 0, 11, and 15 h. Lines indicate the measured apical-to-basal orientation of the DVB relative to a fixed external imaging reference. **B.** DVB orientation (θ*_DVB_*) over the course of the time lapse. The DVB reoriented from approximately 86.3° to 19.1°, corresponding to a net angular reorientation of approximately 67.2°. **C.** Simulated pouch cross-sections at initial, intermediate, and final stages following application of proximal boundary forces at the dorsal and ventral pouch boundaries. The simulated tissue undergoes a θ*_DVB_* = 56. 4° reorientation, similar to the rotation of the *ex vivo* pouch. The θ*_DVB_* values for the simulation images in SI Fig. 4C were achieved by measuring the angle difference between three points: the origin at the simulation pivot point calculated as halfway between the DPB and VPB regions, a directly vertical point, and the midpoint between two central cells outlined in each image magenta.

### Description of Boundary Forces for Rotation

The cuboidal boundary cells on the dorsal proximal boundary (DPB) and the ventral proximal boundary (VPB) are categorized in the model as the boundary regions of the pouch affected by the unfolding dynamics of the external tissue^8,26^. The external boundary forces (*E_Ext_*) are applied upon these boundary cell subdomains to generate stresses which propagate through the rest of the tissue. The center of the pouch, used as the pivot point (P) for the rotation, is calculated at each step by taking half the distance between the DPB and VPB cells. This is indicated in SI Fig. 4Ci-iv by the origin of the θ*_DVB_* angle measurement. The boundary linear force vector with a constant magnitude acts tangentially to the boundary domain of the pouch in a clockwise direction around the central pivot point (See SI Fig. 6C), denoted as *E_Ext_*^54,58^. It is formulated as a kinematic follower force to ensure that the vector dynamically updates its directional orientation relative to the pivot point as the tissue rotates and deforms^40^. This force is applied uniformly to every node in each cell in the DPB and VPB boundary subdomains. As the pouch rotates, deforms slightly, and shifts throughout the simulation, the position of the pivot point and the direction of the force updates dynamically.

The equation for calculating the pivot point coordinates is given by SI Eq. 1 below:

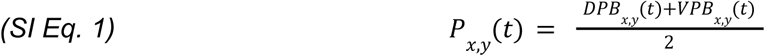

A comprehensive description of the mechanical contributions of tissues surrounding the wing pouch remains incomplete. Previous studies have identified several remodeling events outside of the pouch that could influence the deformation described here^8,26,51^. Myosin-II dependent dynamics within the peripodial epithelium generate large-scale disc bending and permit expansion and retraction of the peripodial membrane during eversion^22^. However, the contribution of these forces to pouch rotation has not been quantitatively resolved in cross-sectional analysis. Similarly, recent studies have identified actomyosin and ECM-dependent remodeling within the hinge and notum, including basal ECM degradation^53^, and remodeling of the apical ECM during fold opening^54^. Nevertheless, how these neighboring tissue dynamics are mechanically coupled to pouch rotation and dorsal-ventral apposition remains unclear. Consistent with these observations, our modeling results (See Fig. 6C) demonstrate that externally imposed boundary forces are sufficient to produce pouch rotation, and promote convergence of the proximal tips, which contributes to subsequent fold closure. These results support the hypothesis that forces transmitted from surrounding tissues contribute to multiple aspects of early pouch deformation, although the current model does not identify the biological origin of these forces or exclude contributions from pouch intrinsic mechanics. Together, previous studies establish that multiple regions surrounding the pouch are mechanically active during eversion, but leave unresolved whether early pouch rotation, tip convergence, and fold closure are driven primarily by forces generated within the pouch, transmitted from neighboring compartments, or coordinated between the two.

**SI Figure 5:**
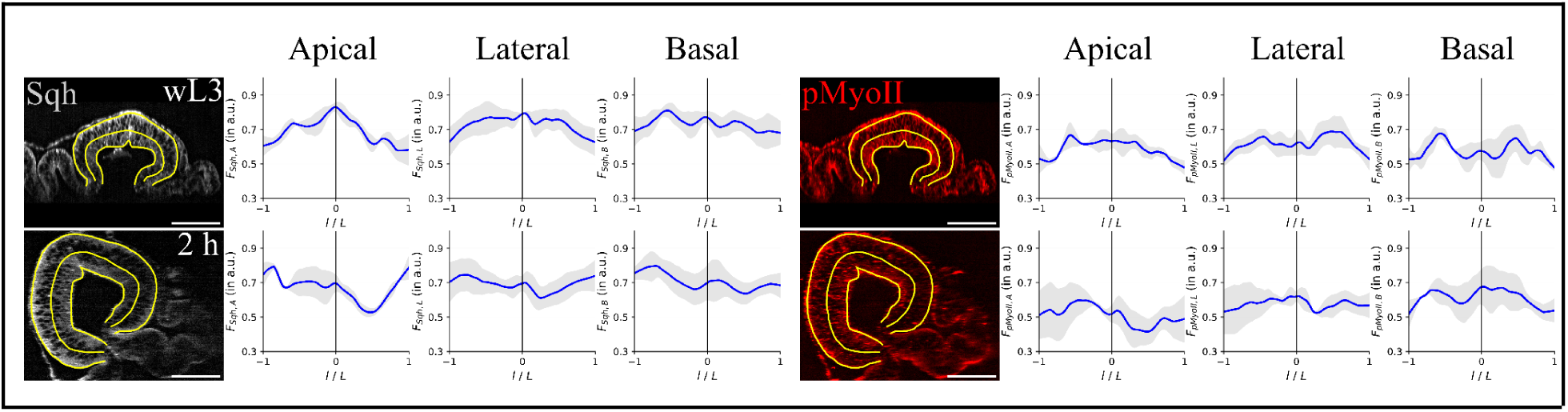
Spatial distribution of Sqh and phosphomyosin II during early wing disc eversion. Representative cross-sectional images of Sqh-mCherry (left) and phospho-myosin II (pNMII; right) in wing imaginal discs at wL3 and 2 h APF. Yellow lines indicate the apical, lateral, and basal contours used for fluorescence-intensity measurements. Corresponding normalized fluorescence-intensity profiles are shown for the apical, lateral, and basal regions. Blue lines indicate the mean and gray shading indicates ±SD across biological replicates (n=3 wind discs per developmental stage). Scale bars, 50 μm.

**SI Figure 6:**
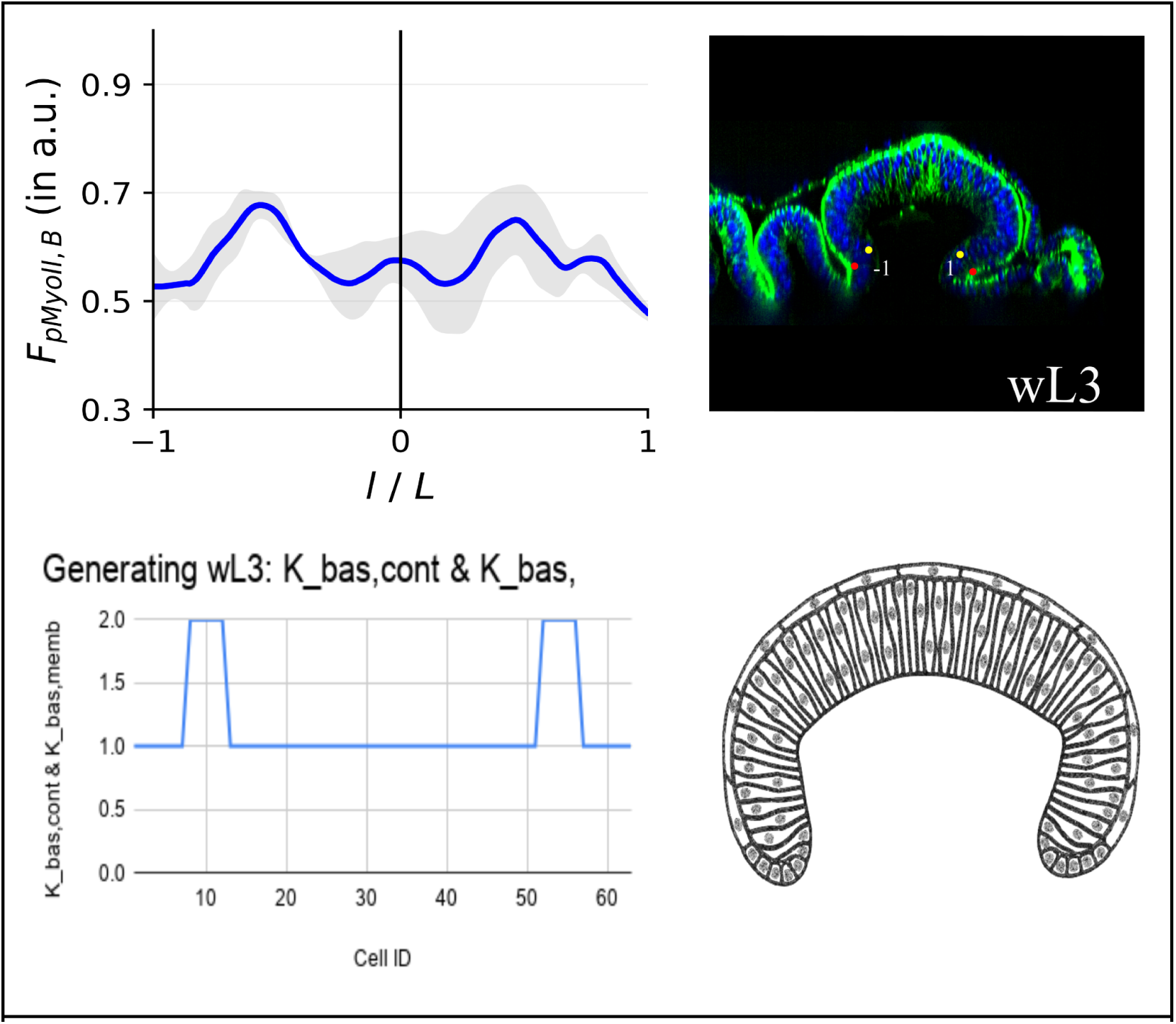
Generating wL3 initial simulation shape for. Fig. 6 **informed by pNMII data.** Spatial distribution of phospho-myosin II (top) in the basal regions of cells in wing imaginal discs at wL3 and the corresponding wing cross-section compared with the spatial distribution of basal actomyosin (K_bas,cont) and basal membrane (K_bas) contractility (bottom) for the simulated cross-section and the resulting model tissue shape used as the initial condition in Fig. 6.

**SI Figure 7:**
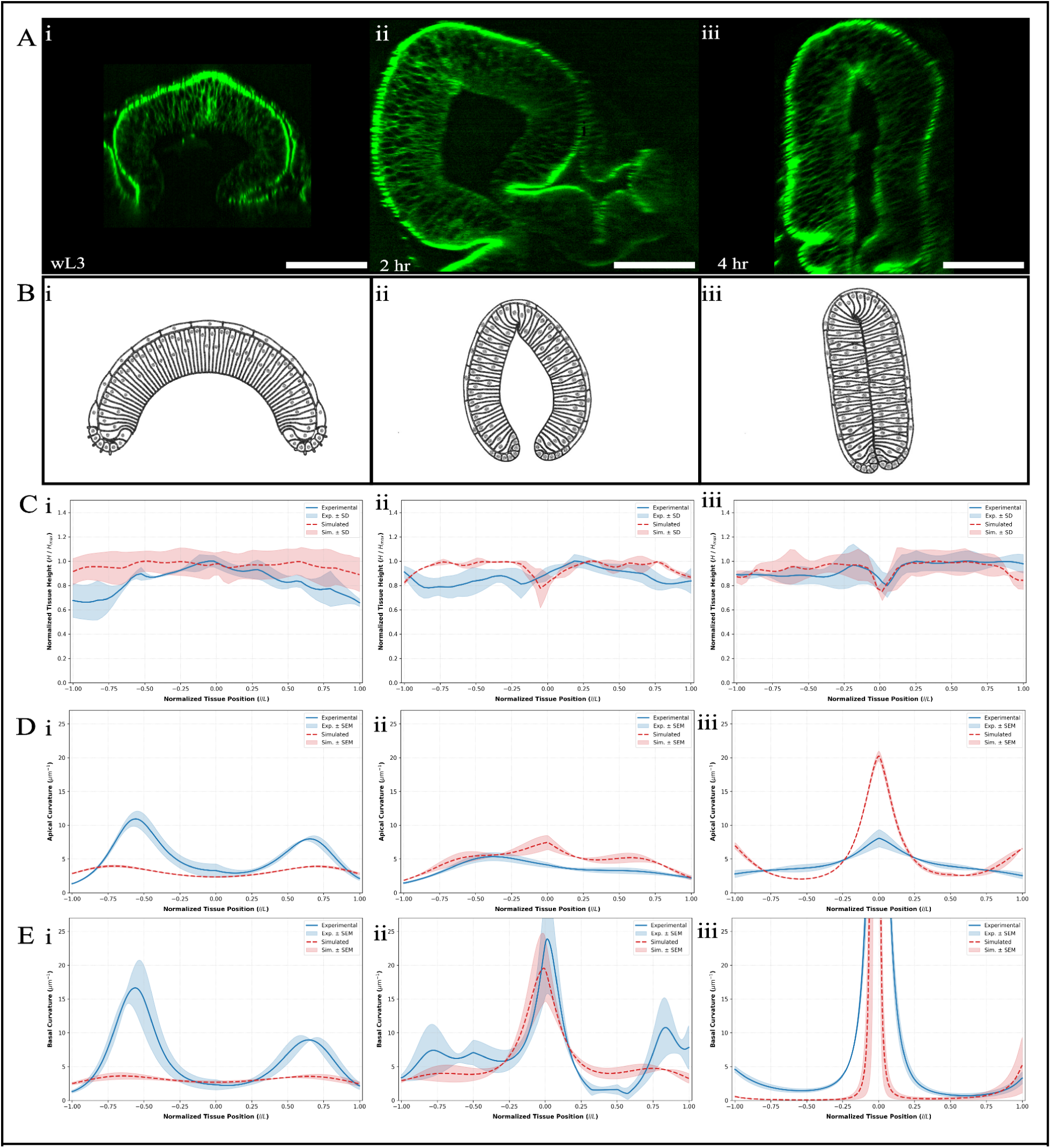
**Alternate eversion simulation (**Fig. 6B**) with dome-like initial tissue shape.** Comparison between simulation results implementing basal ECM detachment, patterned basal actomyosin contractility, boundary forces, and cell rearrangement, and wing disc cross-sections at wL3, 2 h APF, and 4 h APF. **A.** Cross-sections of the wing disc pouch along the DV axis at different time points previously shown in Fig 1. **B.** Simulated wing discs achieved through implementing the biophysical mechanisms outlined in the study. **C.** Plots of the simulated pouch cell heights across the cross-section through wL3, 2 h APF, and 4 h APF normalized to the cell height. Experimental tissue heights are blue and simulated tissue heights are red. **D.** Apical tissue curvature profiles for simulated cross-sections calculated with the same methods as in Fig 1D overlaid with experimental apical curvature profiles. **E.** Basal tissue curvature profiles for simulated cross-sections calculated with the same methods as in Fig 1E overlaid with experimental basal curvature profiles.

**SI Figure 8.**
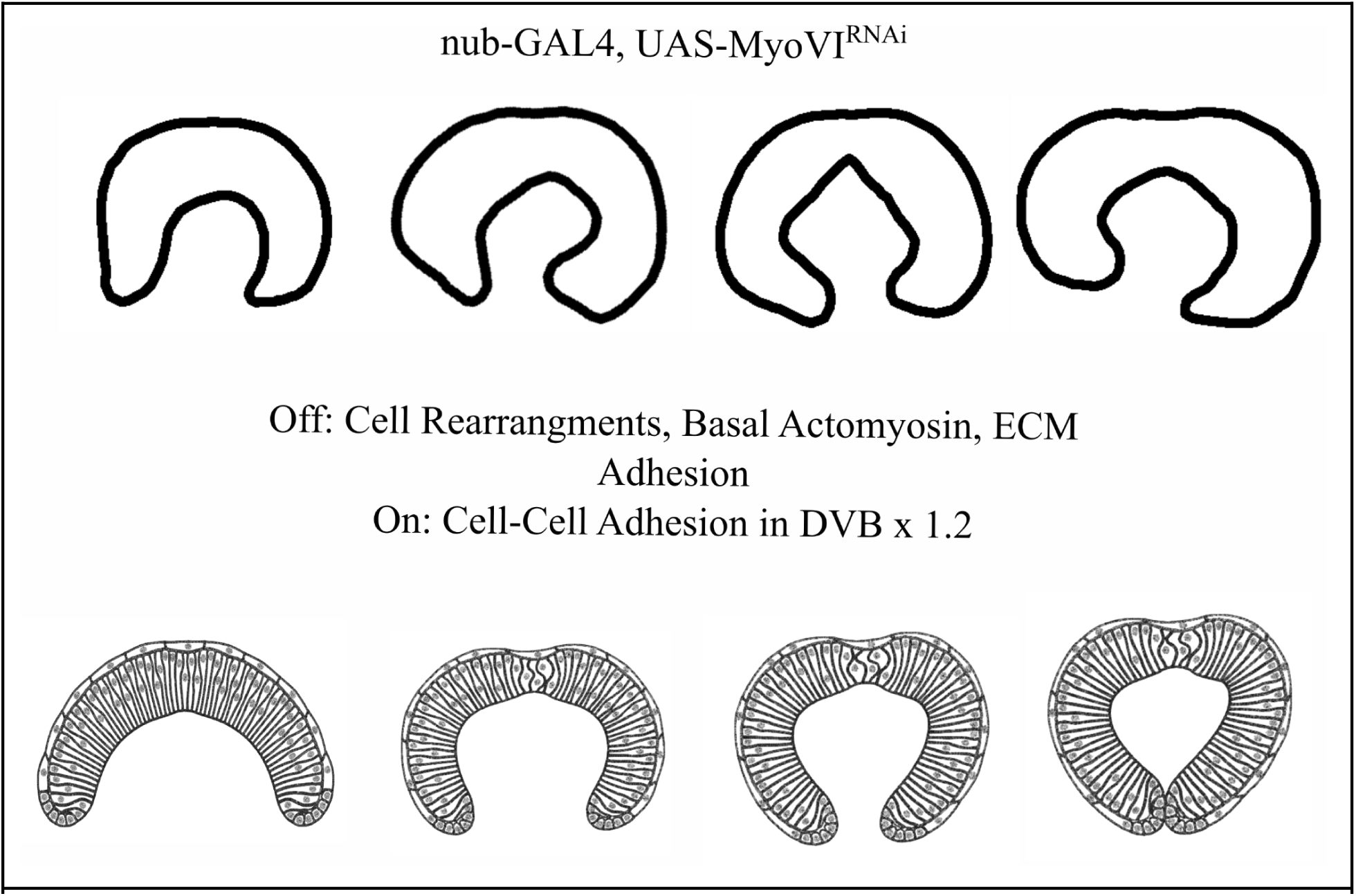
Simulation replicating mutant MyoVI knockdown. Experimental cross-sectional contours along the DV axis (**top**) depict *MyoVI* knockdown (*UAS-jar-RNAi*) wing pouches during eversion^24^. Simulated tissue profiles (**bottom**) replicate this defect by increasing lateral cell-cell adhesion spring forces (x1.2) in the central 5 DVB cells while omitting stochastic cell rearrangements, basal actomyosin contractility, and basal ECM detachment. Inward boundary forces on proximal pouch margins follow what was used in Fig. 4E, 5, and 6.

### Description of Quantification Methods

#### Image analysis and quantification

ND2 image stacks were opened in Fiji/ImageJ, rotated to a consistent orientation, and cropped to the region of interest prior to subsequent processing and analysis. Unless otherwise specified, quantitative measurements of tissue morphology and fluorescence intensity were performed without intensity-based preprocessing, including background subtraction, smoothing, or contrast adjustment. Additional processing described below was performed only for visualization and figure presentation.

For *E-cad::GFP; pnr-GAL4, Sqh::mCherry* samples, background subtraction was performed separately for each fluorescence channel using Fiji’s background-subtraction function without the sliding-paraboloid option. Processing parameters were selected separately for maximum-intensity projections and cross-sectional images. For maximum-intensity projections shown in Figure 1, background-subtraction radii of 150 and 50 pixels were used for DAPI and E-cadherin, respectively. For cross-sectional images shown in Figure 1, radii of 150 and 100 pixels were used for DAPI and E-cadherin, respectively. These processing steps were used only to improve visualization and were not applied to images used for quantitative morphology or fluorescence-intensity measurements.

For visualization of Vkg-GFP and phalloidin-labeled samples, image stacks were processed using an image-enhancement workflow in Fiji. A duplicate of the original stack was subjected to three-dimensional Gaussian blurring (σx = 6, σy = 6, σz = 3). The blurred stack was then subtracted from the original image stack (raw − blurred). The resulting image was background subtracted using a 25-pixel radius with the sliding-paraboloid option enabled. Display minimum and maximum intensity values were adjusted individually as needed for visualization; these adjustments affected only the displayed intensity range and did not alter pixel values used for quantitative analysis.

Sqh-mCherry and pNMyosin-II cross-sectional images used for visualization were background-subtracted with a 100-pixel radius, while raw images were used for quantification.

Live-imaging datasets used for quantitative analysis were not subjected to intensity-based preprocessing and were provided directly to the analysis pipeline following initial cropping and orientation.

#### Tissue height quantification

Tissue height was quantified from cross-sectional images of the E-cadherin channel using a custom Python pipeline. Cross-sections were generated in Fiji/ImageJ using a manually positioned segmented line oriented approximately along the dorsal–ventral axis near the anterior–posterior compartment boundary. Because the precise position of the compartment boundary could not be directly identified from the E-cadherin signal, line placement was determined visually from disc morphology and kept as consistent as possible between samples. Cross-sections were oriented consistently from the dorsal to the ventral side of the pouch prior to analysis.

Cross-sectional images were resampled to isotropic pixels prior to quantitative analysis. The calibrated pixel size of each dataset was verified, and the corresponding conversion factor was supplied to the analysis pipeline to convert distances from pixels to micrometers.

For each cross-section, points were manually placed along the apical and basal surfaces of the pouch. Parametric splines were fit through the annotated points without smoothing and interpolated to generate continuous representations of the apical and basal surfaces. The apical contour was interpolated to 400 points and the basal contour to 60 points. At each basal position, tissue height (*H*) was calculated as the minimum Euclidean distance between the basal point and the interpolated apical contour.

Height profiles were mapped to a normalized dorsal–ventral coordinate ranging from −1 to 1, with measurements consistently oriented from dorsal to ventral. Symmetry between the dorsal and ventral compartments was not assumed, and compartment orientation was retained throughout the analysis. Mean height and standard deviation were calculated across biological samples at each normalized position.

#### Tissue curvature quantification

Curvature was quantified from cross-sectional images of the E-cadherin channel using a custom Python pipeline. Cross-sections were generated and oriented as described above for tissue-height analysis, with profiles consistently annotated from the dorsal to the ventral side of the pouch. Apical and basal surfaces were analyzed separately.

For each surface, points were manually placed along the tissue contour and used to generate a continuous spline representation of the surface. Each contour was sampled at 300 positions and normalized by its total contour length to allow comparison between tissues of different sizes.

For curvature calculation, the normalized contour was fit with a parametric spline and evaluated at 300 positions. Normalized local curvature, κ, was calculated from the first and second derivatives of the fitted contour as:

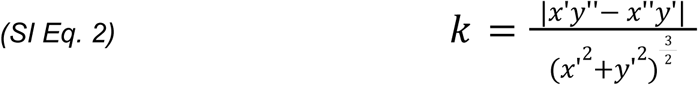

Curvature profiles were lightly smoothed using a Gaussian filter with a 3×3 kernel. Profiles were maintained in their original dorsal-to-ventral orientation, and symmetry between the dorsal and ventral compartments was not assumed. Curvature profiles were mapped to a normalized tissue coordinate ranging from −1 to 1, and mean curvature was calculated across biological samples at each position. Shaded regions in curvature represent ± SEM.

#### Fluorescence signal intensity quantification

Fluorescence intensity profiles of Sqh-mCherry and phospho-myosin II were quantified from cross-sectional images using Fiji/ImageJ and subsequently analyzed using a custom Python pipeline. Cross-sections were generated and oriented as described above for tissue height analysis. Fluorescence intensity was measured along manually defined apical, lateral, and basal contours for the pouch, with profiles consistently oriented from dorsal to ventral. Quantification was performed on images without intensity-based preprocessing or background subtraction, and acquisition settings were held constant across samples used for fluorescence-intensity comparisons.

Because the lengths of the measured contours varied between samples, each intensity profile was interpolated to 300 equally spaced positions along a normalized dorsal-ventral coordinate running from −1 to 1. Each biological sample was then independently normalized to its maximum fluorescence intensity. Thus, normalized values represent the relative spatial distribution of fluorescence within each sample rather than absolute differences in fluorescence intensity between samples or developmental stages.

Normalized profiles were smoothed independently using locally weighted scatterplot smoothing (LOWESS) with a smoothing fraction of 0.15 and no robustifying iterations (it = 0). The same smoothing parameters were applied to all samples and developmental stages. Mean normalized fluorescence intensity and sample standard deviation were subsequently calculated across biological replicates at each normalized position.

#### Apical and basal surface lengths quantification

Apical and basal surface lengths were quantified from experimental cross-sections and computational simulations to compare changes in tissue geometry across developmental stages. For experimental datasets at wL3, 2 h APF, and 4 h APF, the apical and basal surfaces were manually traced from the dorsal-ventral boundary at the center of the pouch cross-section to the dorsal proximal tip. Surface coordinates were extracted in Fiji/ImageJ, and smooth cubic splines were fit through manually placed control points along each contour. Absolute path lengths of the fitted apical and basal surfaces were then calculated.

Equivalent apical and basal contours were extracted from the corresponding portion of the simulated pouch. Experimental and simulated surface lengths were normalized to their respective wL3 baseline values to calculate fold changes across developmental stages. Experimental values are reported as the mean ± SD across biological replicates (*n* = 3 wing discs per developmental stage.) Surface-length measurements were subsequently used to calculate the error between simulated and experimental surface-length changes for calibration of the cell rearrangement model, as described below.

Agreement between simulated and experimental surface-length changes was quantified by calculating the absolute difference between the mean experimental and simulated surface lengths at each developmental stage:

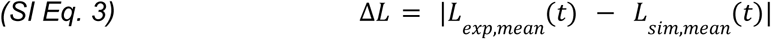

where *L_exp,mean_*(*t*) and *L_sim,mean_*(*t*) denote the mean experimental and simulated surface length, respectively, at developmental time *t*. To quantify improvement associated with active cell rearrangements, the surface-length error for the active rearrangement simulation was compared with that of the corresponding inactive rearrangement simulation:

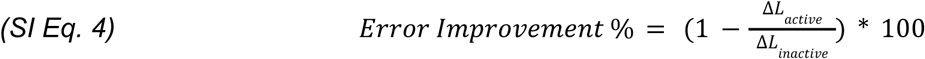

Here,

Δ*L_active_* and Δ*L_inactive_* represent the surface-length errors calculated using SI Eq. 3 for simulations with and without active cell rearrangements, respectively. Error improvement was calculated separately for apical and basal surface lengths at 2 h and 4 h APF.

**SI Table 1:** Improvement in model agreement following inclusion of cell rearrangements.

| Tissue Surface and Time Point | Normalized Experimental Value<br>*L*<sub>*exp,mean*</sub>(*t*) | Active Error<br>Δ*L*<sub>*active*</sub>(t) | Inactive Error<br>Δ*L*<sub>*inactive*</sub>(t) | Error Improvement Percentage |
|---|---|---|---|---|
| 2 h Apical Surface Lengths | 1.2344 | 0.0448 | 0.0381 | 59.94% |
| 2 h Basal Surface Lengths | 1.1865 | 0.0344 | 0.0858 | -17.85% |
| 4 h Apical Surface Lengths | 1.3055 | 0.0579 | 0.1235 | 53.12% |
| 4 h Basal Surface Lengths | 1.5106 | 0.1130 | 0.3611 | 68.71% |
| Average Improvement value referenced in main text |  |  |  | 40.98% |

#### Rotational Analysis

Dorsal-ventral boundary (DVB) reorientation was quantified from cross-sectional live-imaging time series using Fiji/ImageJ and a custom Python analysis pipeline. Cross-sectional images were resampled to isotropic pixels prior to ROI placement and angular quantification. For each selected timepoint, a circular ROI was manually positioned at the apical DVB landmark, and a straightline ROI was drawn through the DVB to approximate its apical-to-basal orientation. A fixed directed reference line was defined for each dataset and used as the common reference for all time points within that sample.

The center of the circular DVB ROI and endpoints of the corresponding straight-line ROI were extracted from the Fiji ROI files. The endpoints closest to the DVB landmark were designated as the apical endpoint and the opposite endpoint as the basal endpoint, thereby defining an apical-to-basal DVB vector. The DVB orientation angle, θ*_DVB_*, was calculated as the angle between the DVB vector and the directed reference vector:

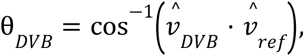

Where *v̂_DVB_* and *v̂_ref_* are unit vectors describing the apical-to-basal DVB orientation and reference direction, respectively. Angles therefore ranged from 0° to 180°.

Measurements were mapped to their corresponding positions in the original time-lapse sequence using the source-frame indices and the acquisition time interval. Absolute values were retained for each time point. Angular reorientation relative to the initial measurement was additionally calculated as

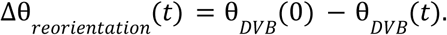

Individual biological replicates were analyzed independently and plotted as separate trajectories rather than averages into a single population curve.

DVB orientation in computational simulations was quantified using the same apical-to-basal vector and directed-reference definition applied to the experimental data. Selected simulation snapshots were measured independently and compared with experimental trajectories. Simulation snapshots were treated as discrete measurements; connecting lines between simulated points were used only as visual guides. Where simulation times had not been independently validated, their positions along the time axis were treated as illustrative and were not used to infer agreement in the timing or rate of DVB reorientation.

https://drive.google.com/file/d/18Qq--FingEKbftQAP4fE51NRXr-4a7cK/view?usp=sharing

**SI Video 1. Live imaging time lapse of wing disc eversion.**

https://drive.google.com/file/d/1Pug6-FIZ8PcEIuGAkcL6J-u41hhGHU_w/view?usp=sharing

**SI Video 2. Simulation Video for Fig. 3B**.

https://drive.google.com/file/d/1tqXI7Rbs_2Nkm9usdvW2Ktfgv2pzi5Qu/view?usp=sharing

**SI Video 3. Simulation Video for Fig. 3C**.

https://drive.google.com/file/d/1Emajt-nJVqBdWNmfJPKm267WQkOKdDXu/view?usp**= sharing**

**SI Video 4. Simulation Video for Fig. 3D**.

https://drive.google.com/file/d/12gtD-ABbxOm5Eq_NbbwsF9H6X4V3ZAga/view?usp=sharing

**SI Video 5. Simulation Video for Fig. 4C**.

https://drive.google.com/file/d/1QPYSXFnjKYtKiC56gEcMO1TNyWVlpiAL/view?usp=sharing

**SI Video 7. Simulation Video for Fig. 4E**.

https://drive.google.com/file/d/1LlD9Sq0wCHSPE4w88_PXRcosKK6vo2Yi/view?usp=sharing

**SI Video 8. Simulation Video for Fig. 5C**.

https://drive.google.com/file/d/1urVmez5qtafIZ85Jk1ciMSrLNTij9b8w/view?usp=sharing

**SI Video 9. Simulation Video for Fig. 5D**.

https://drive.google.com/file/d/1EXYkXDvdQFW00Dypr9WZuHAjvjwClwzH/view?usp=sharing

**SI Video 10. Simulation Video for Fig. 6**.

## References

1. Guillot C, Lecuit T. Mechanics of Epithelial Tissue Homeostasis and Morphogenesis. Science. 2013;340(6137):1185–1189. doi:10.1126/science.1235249

2. Heisenberg CP, Bellaïche Y. Forces in Tissue Morphogenesis and Patterning. Cell. 2013;153(5):948–962. doi:10.1016/j.cell.2013.05.008

3. Pickup MW, Mouw JK, Weaver VM. The extracellular matrix modulates the hallmarks of cancer. EMBO Rep. 2014;15(12):1243–1253. doi:10.15252/embr.201439246

4. Lee YS, Wysocki A, Warburton D, Tuan TL. Wound Healing in Development. Birth Defects Res Part C Embryo Today Rev. 2012;96(3):213–222. doi:10.1002/bdrc.21017

5. Dahmann C, Oates AC, Brand M. Boundary formation and maintenance in tissue development. Nat Rev Genet. 2011;12(1):43–55. doi:10.1038/nrg2902

6. Aldaz S, Escudero LM, Freeman M. Live imaging of Drosophila imaginal disc development. Proc Natl Acad Sci. 2010;107(32):14217–14222. doi:10.1073/pnas.1008623107

7. Tripathi BK, Irvine KD. The wing imaginal disc. Genetics. 2022;220(4):iyac020. doi:10.1093/genetics/iyac020

8. Pastor-Pareja JC, Xu T. Shaping Cells and Organs in Drosophila by Opposing Roles of Fat Body-Secreted Collagen IV and Perlecan. Dev Cell. 2011;21(2):245–256. doi:10.1016/j.devcel.2011.06.026

9. Hariharan IK. Organ size control: lessons from Drosophila. Dev Cell. 2015;34(3):255–265. doi:10.1016/j.devcel.2015.07.012

10. Garcia-Bellido A, Ripoll P, Morata G. Developmental Compartmentalisation of the Wing Disk of Drosophila. Nature New Biol. 1973;245(147):251–253. doi:10.1038/newbio245251a0

11. Nellen D, Burke R, Struhl G, Basler K. Direct and Long-Range Action of a DPP Morphogen Gradient. Cell. 1996;85(3):357–368. doi:10.1016/S0092-8674(00)81114-9

12. Parker J, Struhl G. Control of Drosophila wing size by morphogen range and hormonal gating. Proc Natl Acad Sci U S A. 2020;117(50):31935–31944. doi:10.1073/pnas.2018196117

13. Shaikh R, Larson NJ, Kam J, et al. Optimal performance objectives in the highly conserved bone morphogenetic protein signaling pathway. Npj Syst Biol Appl. 2024;10(1):103. doi:10.1038/s41540-024-00430-9

14. Zecca M, Basler K, Struhl G. Direct and Long-Range Action of a Wingless Morphogen Gradient. Cell. 1996;87(5):833–844. doi:10.1016/S0092-8674(00)81991-1

15. Baonza A, Garcia-Bellido A. Notch signaling directly controls cell proliferation in the Drosophila wing disc. Proc Natl Acad Sci. 2000;97(6):2609–2614. doi:10.1073/pnas.040576497

16. Pan Y, Alégot H, Rauskolb C, Irvine KD. The dynamics of Hippo signaling during Drosophila wing development. Dev Camb Engl. 2018;145(20):dev165712. doi:10.1242/dev.165712

17. Kumar N, Mim MS, Dowling A, Zartman JJ. Reverse engineering morphogenesis through Bayesian optimization of physics-based models. Npj Syst Biol Appl. 2024;10(1):49. doi:10.1038/s41540-024-00375-z

18. Kumar N, Rangel Ambriz J, Tsai K, et al. Balancing competing effects of tissue growth and cytoskeletal regulation during Drosophila wing disc development. Nat Commun. 2024;15(1):2477. doi:10.1038/s41467-024-46698-7

19. Martin AC, Kaschube M, Wieschaus EF. Pulsed contractions of an actin-myosin network drive apical constriction. Nature. 2009;457(7228):495–499. doi:10.1038/nature07522

20. Milner MJ. The eversion and differentiation of Drosophila melanogaster leg and wing imaginal discs cultured in vitro with an optimal concentration of beta-ecdysone. J Embryol Exp Morphol. 1977;37(1):105–117.

21. Uyehara CM, Leatham-Jensen M, McKay DJ. Opportunistic binding of EcR to open chromatin drives tissue-specific developmental responses. Proc Natl Acad Sci U S A. 2022;119(40):e2208935119. doi:10.1073/pnas.2208935119

22. Aldaz S, Escudero LM, Freeman M. Dual role of myosin II during Drosophila imaginal disc metamorphosis. Nat Commun. 2013;4(1):1761. doi:10.1038/ncomms2763

23. Diaz-de-la-Loza M del C, Ray RP, Ganguly PS, et al. Apical and Basal Matrix Remodeling Control Epithelial Morphogenesis. Dev Cell. 2018;46(1):23–39.e5. doi:10.1016/j.devcel.2018.06.006

24. Fuhrmann JF, Krishna A, Paijmans J, et al. Active shape programming drives Drosophila wing disc eversion. Sci Adv. 2024;10(32):eadp0860. doi:10.1126/sciadv.adp0860

25. de Celis JF. Pattern formation in the Drosophila wing: The development of the veins. BioEssays. 2003;25(5):443–451. doi:10.1002/bies.10258

26. Diaz de la Loza MC, Thompson BJ. Forces shaping the *Drosophila* wing. Mech Dev. 2017;144:23–32. doi:10.1016/j.mod.2016.10.003

27. Nienhaus U, Aegerter-Wilmsen T, Aegerter CM. In-vivo imaging of the Drosophila wing imaginal disc over time: novel insights on growth and boundary formation. PloS One. 2012;7(10):e47594. doi:10.1371/journal.pone.0047594

28. Harmansa S, Erlich A, Eloy C, Zurlo G, Lecuit T. Growth anisotropy of the extracellular matrix shapes a developing organ. Nat Commun. 2023;14(1):1220. doi:10.1038/s41467-023-36739-y

29. Pastor-Pareja JC, Grawe F, Martín-Blanco E, García-Bellido A. Invasive cell behavior during Drosophila imaginal disc eversion is mediated by the JNK signaling cascade. Dev Cell. 2004;7(3):387–399. doi:10.1016/j.devcel.2004.07.022

30. Proag A, Monier B, Suzanne M. Physical and functional cell-matrix uncoupling in a developing tissue under tension. Dev Camb Engl. 2019;146(11):dev172577. doi:10.1242/dev.172577

31. Chaturvedi R, Huang C, Kazmierczak B, et al. On multiscale approaches to three-dimensional modelling of morphogenesis. J R Soc Interface. 2005;2(3):237–253. doi:10.1098/rsif.2005.0033

32. Gsell S, Tlili S, Merkel M, Lenne PF. Marangoni-like tissue flows enhance symmetry breaking of embryonic organoids. Nat Phys. 2025;21(4):644–653. doi:10.1038/s41567-025-02802-2

33. Hannezo E, Prost J, Joanny JF. Theory of epithelial sheet morphology in three dimensions. Proc Natl Acad Sci U S A. 2014;111(1):27–32. doi:10.1073/pnas.1312076111

34. Trushko A, Di Meglio I, Merzouki A, et al. Buckling of an Epithelium Growing under Spherical Confinement. Dev Cell. 2020;54(5):655–668.e6. doi:10.1016/j.devcel.2020.07.019

35. Yang H, Roy G, Nguyen AQ, et al. MultiCell: geometric learning in multicellular development. Nat Methods. 2026;23(3):617–625. doi:10.1038/s41592-025-02983-x

36. Graner F, Glazier JA. Simulation of biological cell sorting using a two-dimensional extended Potts model. Phys Rev Lett. 1992;69(13):2013–2016. doi:10.1103/PhysRevLett.69.2013

37. Voss-Böhme A. Multi-Scale Modeling in Morphogenesis: A Critical Analysis of the Cellular Potts Model. PLOS ONE. 2012;7(9):e42852. doi:10.1371/journal.pone.0042852

38. Minsuk SB, Sego TJ, Umulis DM, Mullins MC, Glazier JA. Modeling epithelial deformation and cell rearrangement in response to external forces during Zebrafish epiboly. Npj Syst Biol Appl. 2026;12(1):63. doi:10.1038/s41540-026-00708-0

39. Xu Z, Lioi J, Mu J, et al. A Multiscale Model of Venous Thrombus Formation with Surface-Mediated Control of Blood Coagulation Cascade. Biophys J. 2010;98(9):1723–1732. doi:10.1016/j.bpj.2009.12.4331

40. Van Liedekerke P, Palm MM, Jagiella N, Drasdo D. Simulating tissue mechanics with agent-based models: concepts, perspectives and some novel results. Comput Part Mech. 2015;2(4):401–444. doi:10.1007/s40571-015-0082-3

41. Alt S, Ganguly P, Salbreux G. Vertex models: from cell mechanics to tissue morphogenesis. Philos Trans R Soc Lond B Biol Sci. 2017;372(1720):20150520. doi:10.1098/rstb.2015.0520

42. Bielmeier C, Alt S, Weichselberger V, et al. Interface Contractility between Differently Fated Cells Drives Cell Elimination and Cyst Formation. Curr Biol. 2016;26(5):563–574. doi:10.1016/j.cub.2015.12.063

43. Newman TJ. Modeling multicellular systems using subcellular elements. Math Biosci Eng MBE. 2005;2(3):613–624. doi:10.3934/mbe.2005.2.613

44. Sandersius SA, Weijer CJ, Newman TJ. Emergent cell and tissue dynamics from subcellular modeling of active biomechanical processes. Phys Biol. 2011;8(4):045007. doi:10.1088/1478-3975/8/4/045007

45. Michael C, Banwarth-Kuhn M, Rodriguez K, et al. Role of turgor-pressure induced boundary tension in the maintenance of the shoot apical meristem of Arabidopsis thaliana. J R Soc Interface. 2023;20(203):20230173. doi:10.1098/rsif.2023.0173

46. Nematbakhsh A, Levis M, Kumar N, Chen W, Zartman JJ, Alber M. Epithelial organ shape is generated by patterned actomyosin contractility and maintained by the extracellular matrix. PLOS Comput Biol. 2020;16(8):e1008105. doi:10.1371/journal.pcbi.1008105

47. Newell-Litwa KA, Horwitz R, Lamers ML. Non-muscle myosin II in disease: mechanisms and therapeutic opportunities. Dis Model Mech. 2015;8(12):1495–1515. doi:10.1242/dmm.022103

48. Weißenbruch K, Grewe J, Hippler M, et al. Distinct roles of nonmuscle myosin II isoforms for establishing tension and elasticity during cell morphodynamics. eLife. 10:e71888. doi:10.7554/eLife.71888

49. Li S, Liu ZY, Li H, et al. Basal actomyosin pulses expand epithelium coordinating cell flattening and tissue elongation. Nat Commun. 2024;15(1):3000. doi:10.1038/s41467-024-47236-1

50. Solon J, Kaya-Copur A, Colombelli J, Brunner D. Pulsed forces timed by a ratchet-like mechanism drive directed tissue movement during dorsal closure. Cell. 2009;137(7):1331–1342. doi:10.1016/j.cell.2009.03.050

51. Srivastava A, Pastor-Pareja JC, Igaki T, Pagliarini R, Xu T. Basement membrane remodeling is essential for Drosophila disc eversion and tumor invasion. Proc Natl Acad Sci U S A. 2007;104(8):2721–2726. doi:10.1073/pnas.0611666104

52. Valencia-Expósito A, Khalilgharibi N, Martínez-Abarca Millán A, Mao Y, Martín-Bermudo MD. Local weakening of cell-extracellular matrix adhesion triggers basal epithelial tissue folding. EMBO J. 2025;44(7):2002–2024. doi:10.1038/s44318-025-00384-6

53. Hinata C, Nakagawa H, Nonaka S, Nozaki K, Sato Y, Ohsawa S. Extrinsic MMPs drive epithelial shape change via basal extracellular matrix disassembly in the Drosophila wing disc. Commun Biol. Published online May 29, 2026. doi:10.1038/s42003-026-10376-9

54. Fuhrmann JF, Schimmenti VM, Cwikla G, et al. Apical extracellular matrix regulates fold morphogenesis in the Drosophila wing disc. bioRxiv. Preprint posted online September 6, 2025:2025.09.06.674631. doi:10.1101/2025.09.06.674631

55. Lu L, Jin P, Pang G, Zhang Z, Karniadakis GE. Learning nonlinear operators via DeepONet based on the universal approximation theorem of operators. Nat Mach Intell. 2021;3(3):218–229. doi:10.1038/s42256-021-00302-5

56. Peng GCY, Alber M, Buganza Tepole A, et al. Multiscale Modeling Meets Machine Learning: What Can We Learn? Arch Comput Methods Eng. 2021;28(3):1017–1037. doi:10.1007/s11831-020-09405-5

57. Ramezani A, Britton S, Zandi R, Alber M, Nematbakhsh A, Chen W. A multiscale chemical-mechanical model predicts impact of morphogen spreading on tissue growth. Npj Syst Biol Appl. 2023;9(1):16. doi:10.1038/s41540-023-00278-5

58. Bai J, Zeng X. Computational modeling and simulation of epithelial wound closure. Sci Rep. 2023;13:6265. doi:10.1038/s41598-023-33111-4

59. Mietke A. Tissue wrinkles foreshadow cancer. Nat Phys. 2025;21(2):183–184. doi:10.1038/s41567-024-02763-y

60. Velagala V, Zartman JJ. Pinching and pushing: fold formation in the Drosophila dorsal epidermis. Biophys J. 2021;120(19):4202–4213. doi:10.1016/j.bpj.2021.08.028

61. Zartman JJ, Shvartsman SY. Unit operations of tissue development: epithelial folding. Annu Rev Chem Biomol Eng. 2010;1:231–246. doi:10.1146/annurev-chembioeng-073009-100919

62. Nematbakhsh A, Sun W, Brodskiy PA, et al. Multi-scale computational study of the mechanical regulation of cell mitotic rounding in epithelia. PLoS Comput Biol. 2017;13(5):e1005533. doi:10.1371/journal.pcbi.1005533

63. Marin-Riera M, Brun-Usan M, Zimm R, Välikangas T, Salazar-Ciudad I. Computational modeling of development by epithelia, mesenchyme and their interactions: a unified model. Bioinformatics. 2016;32(2):219–225. doi:10.1093/bioinformatics/btv527

64. Jamali Y, Azimi M, Mofrad MRK. A Sub-Cellular Viscoelastic Model for Cell Population Mechanics. PLoS ONE. 2010;5(8):e12097. doi:10.1371/journal.pone.0012097

65. Micikevicius P. 3D finite difference computation on GPUs using CUDA. In: Proceedings of 2nd Workshop on General Purpose Processing on Graphics Processing Units. GPGPU-2. Association for Computing Machinery; 2009:79–84. doi:10.1145/1513895.1513905

66. Ahrens J, Geveci B, Law C. ParaView: An End-User Tool for Large-Data Visualization. In: Elsevier; 2005:717–731. doi:10.1016/B978-012387582-2/50038-1

67. Helton JC, Davis FJ. Latin hypercube sampling and the propagation of uncertainty in analyses of complex systems. Reliab Eng Syst Saf. 2003;81(1):23–69. doi:10.1016/S0951-8320(03)00058-9

68. McKay MD, Beckman RJ, Conover WJ. Comparison of Three Methods for Selecting Values of Input Variables in the Analysis of Output from a Computer Code. Technometrics. 1979;21(2):239–245. doi:10.1080/00401706.1979.10489755

69. Huang J, Zhou W, Dong W, Watson AM, Hong Y. Directed, efficient, and versatile modifications of the Drosophila genome by genomic engineering. Proc Natl Acad Sci. 2009;106(20):8284–8289. doi:10.1073/pnas.0900641106

70. Duda M, Kirkland NJ, Khalilgharibi N, et al. Polarization of Myosin II Refines Tissue Material Properties to Buffer Mechanical Stress. Dev Cell. 2019;48(2):245–260.e7. doi:10.1016/j.devcel.2018.12.020

71. Buszczak M, Paterno S, Lighthouse D, et al. The Carnegie Protein Trap Library: A Versatile Tool for Drosophila Developmental Studies. Genetics. 2007;175(3):1505–1531. doi:10.1534/genetics.106.065961

72. Gazzo DV, Zartman JJ. Calcium Imaging in Drosophila. In: Gorvin CM, ed. Calcium Signaling: Methods and Protocols. Springer US; 2025:257–271. doi:10.1007/978-1-0716-4164-4_19

