## Supplementary figures and images for "A multiscale modeling - experimental framework reveals the interplay between drivers of *Drosophila* wing morphogenesis"

### 0 hr ECAD Apical Surface Length.png

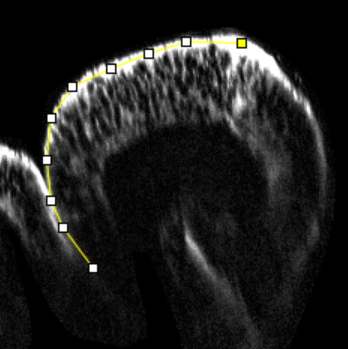

### 0 hr ECAD Basal Surface Length.png

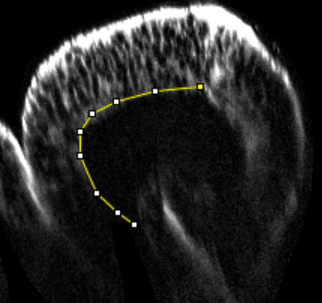

### 0hAPF Simulated Shape.jpg

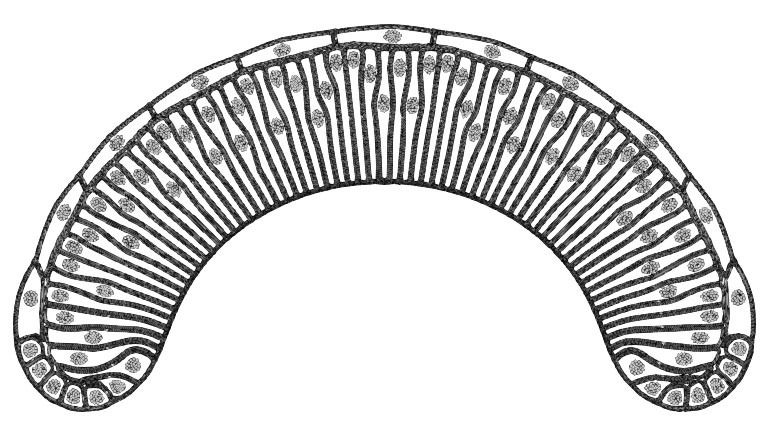

### 2 hr Apical Surface.jpeg

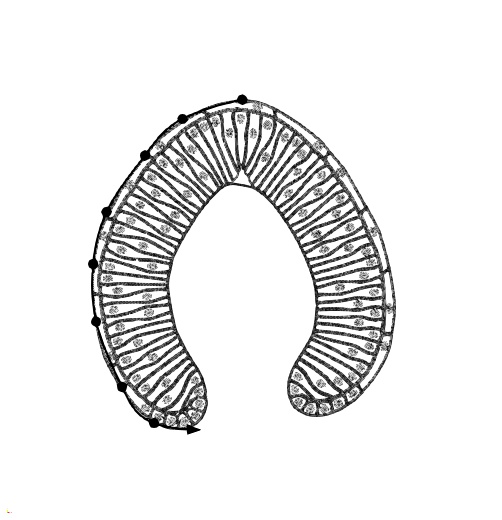

### 2 hr ECAD Apical Surface Length.jpg

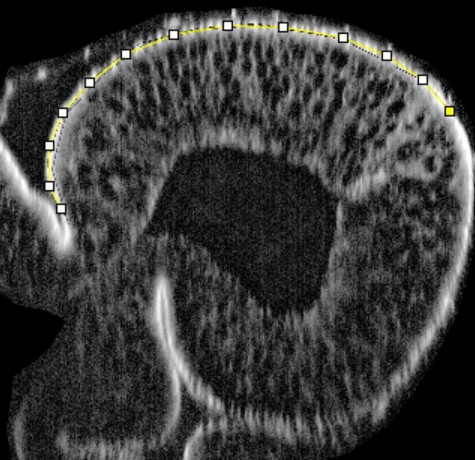

### 2 hr ECAD Basal Surface Length.png

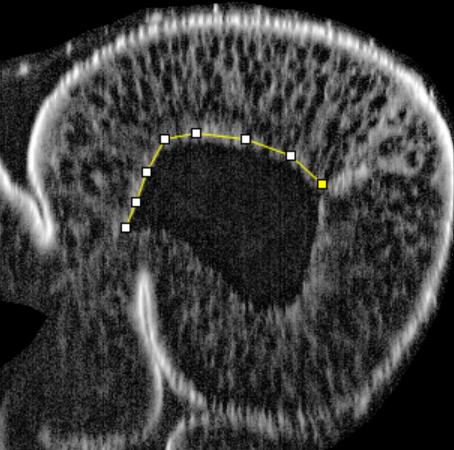

### 2h_height_overlay.png

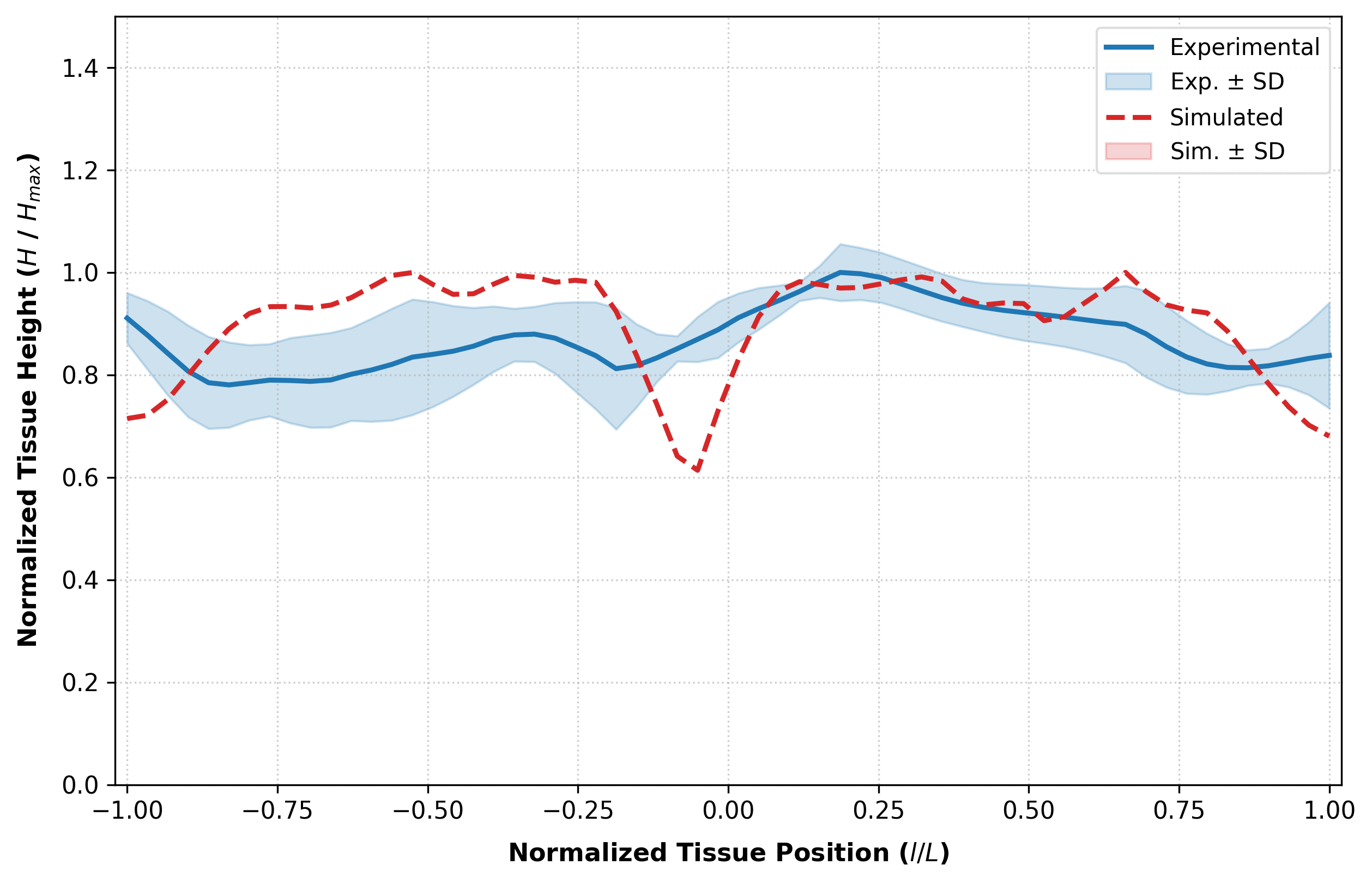

### 2hAPF Simulated Tissue Shape 2.jpg

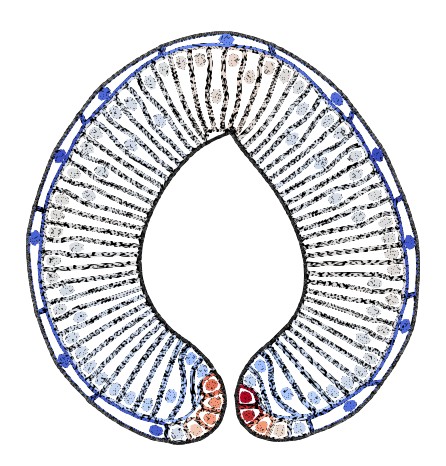

### 2hAPF Simulated Tissue Shape 2_overlay.jpg

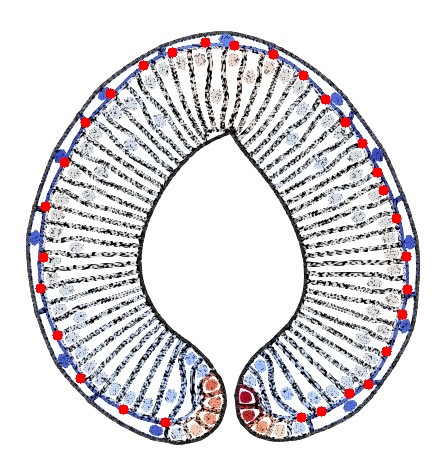

### 2hr_height_overlay.png

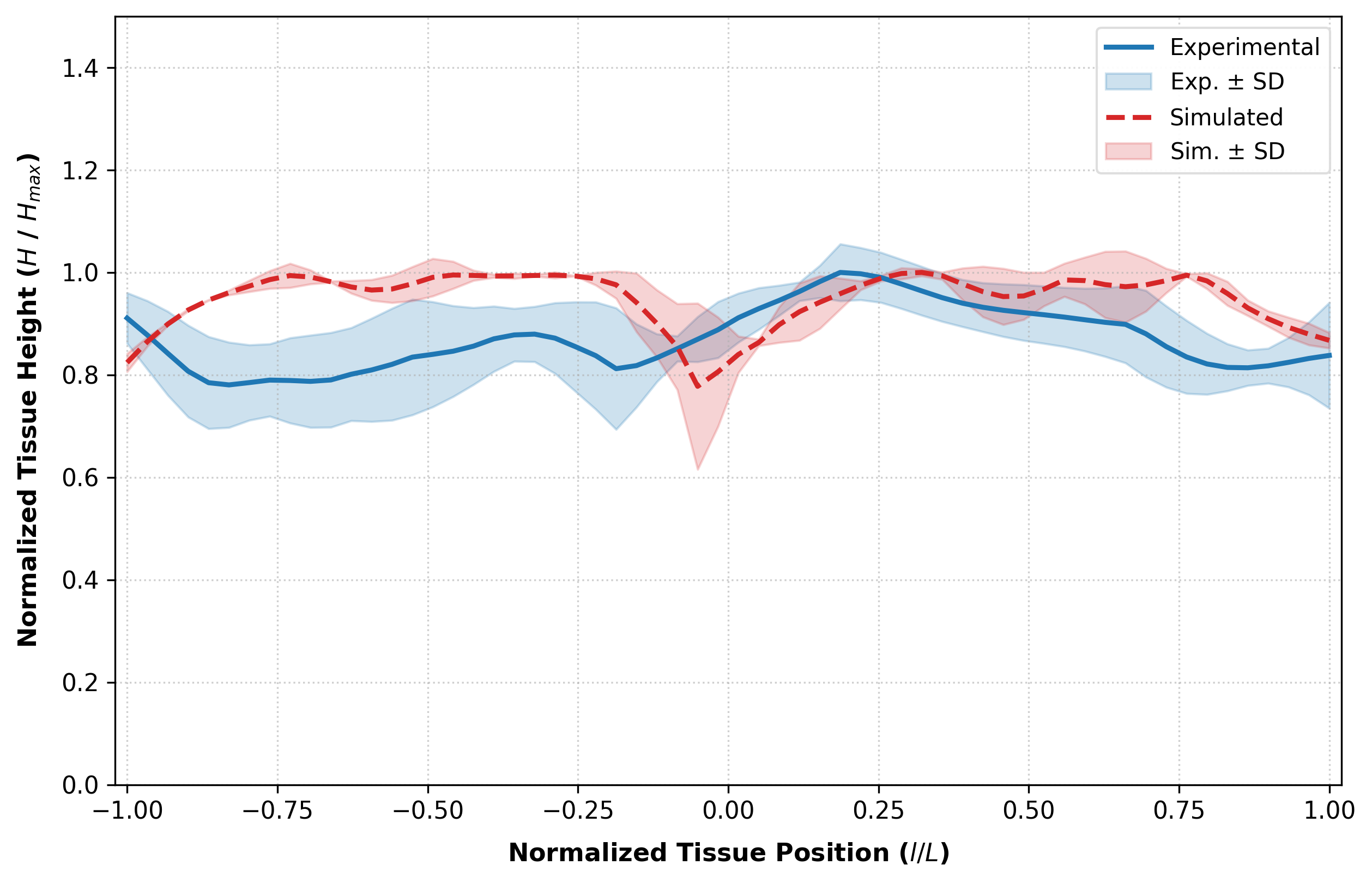

### 2hrSimulatedShape.jpg

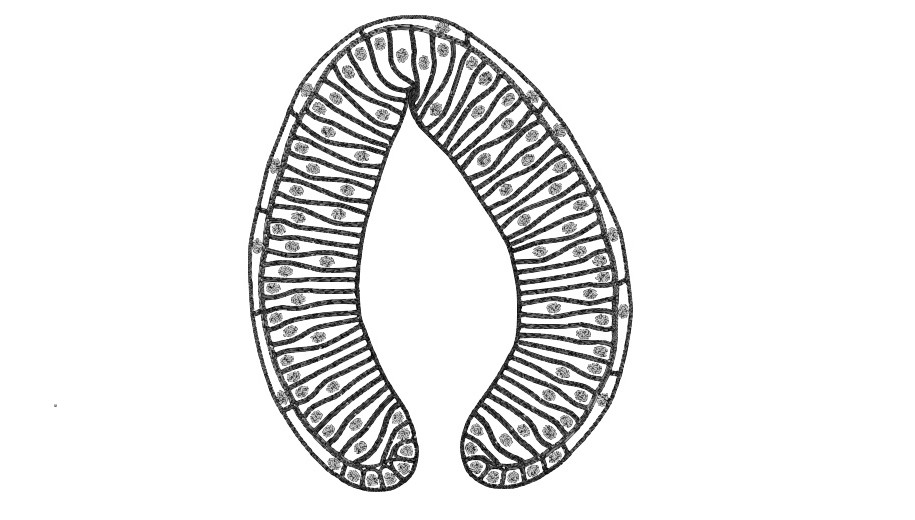

### 2hrSimulatedShape_overlay.jpg

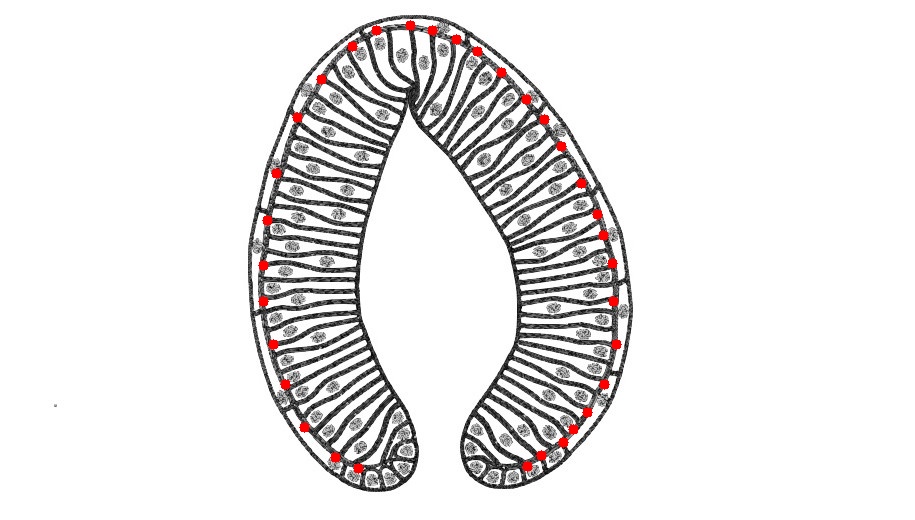

### 4 hr Apical Surface.jpeg

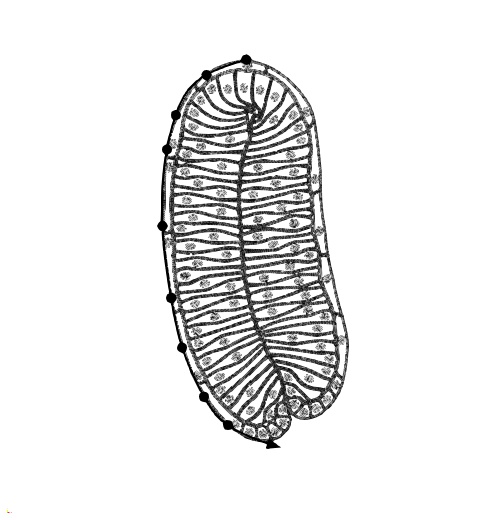

### 4 hr ECAD Apical Surface Length.png

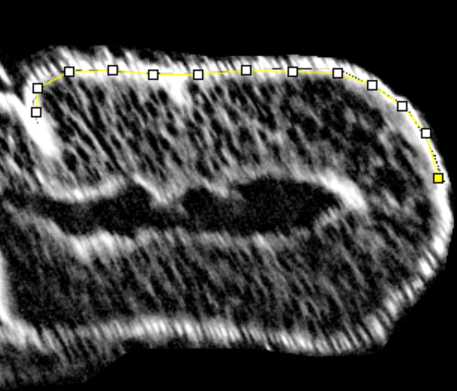

### 4 hr ECAD Basal Surface Length.png

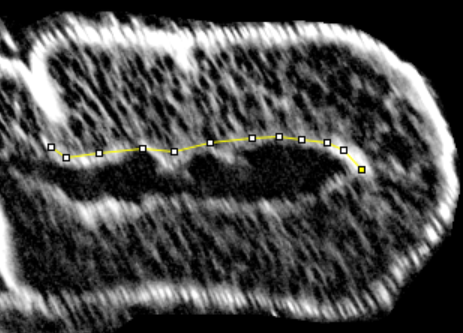

### 4h_height_overlay.png

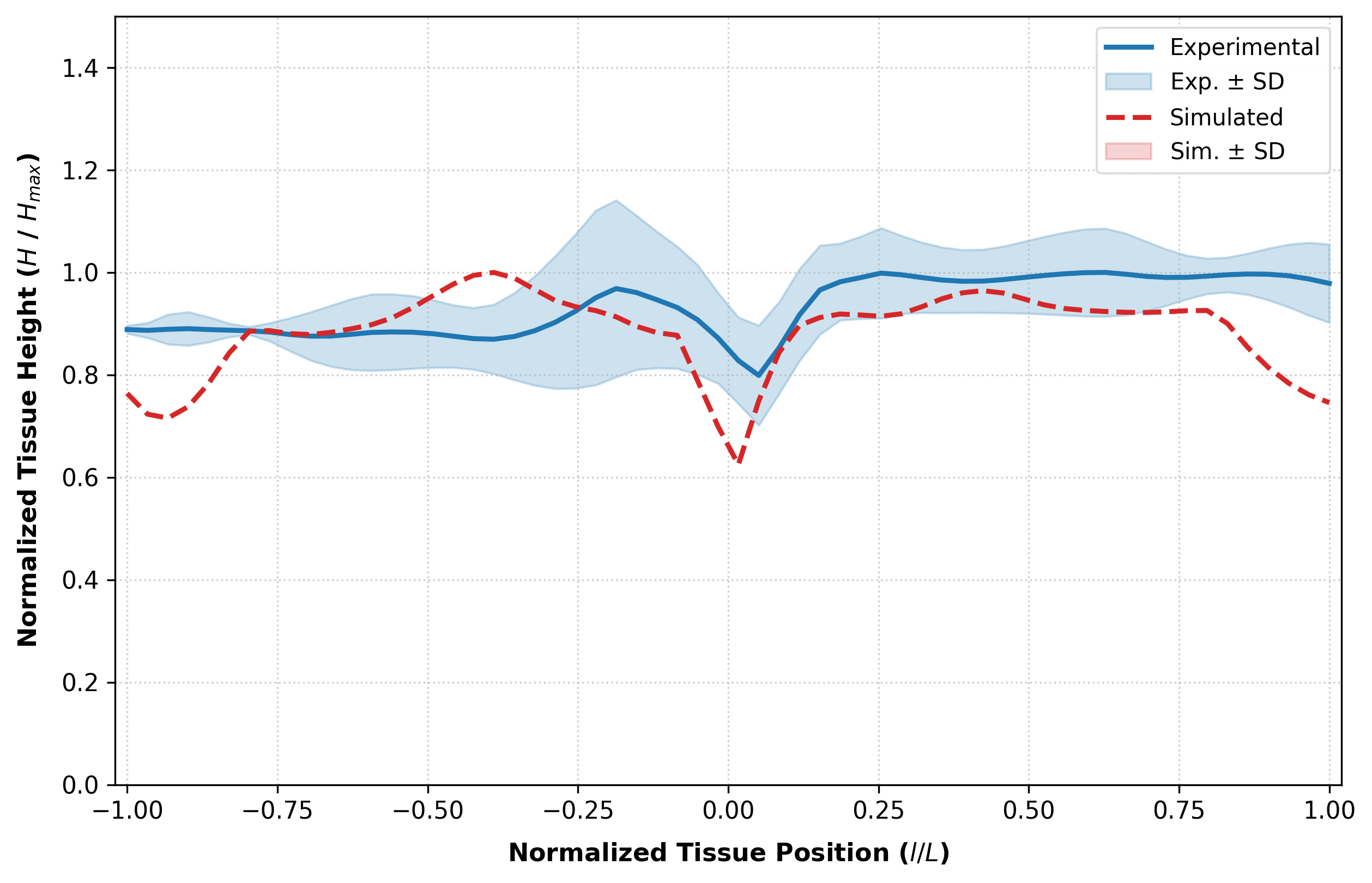

### 4hAPF Simulated Tissue Shape.jpg

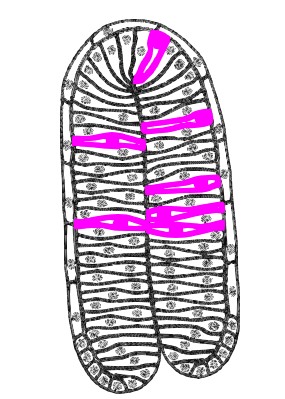

### 4hAPF Simulated Tissue Shape_overlay.jpg

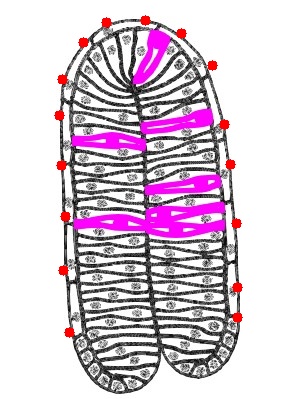

### 4hr_height_overlay.png

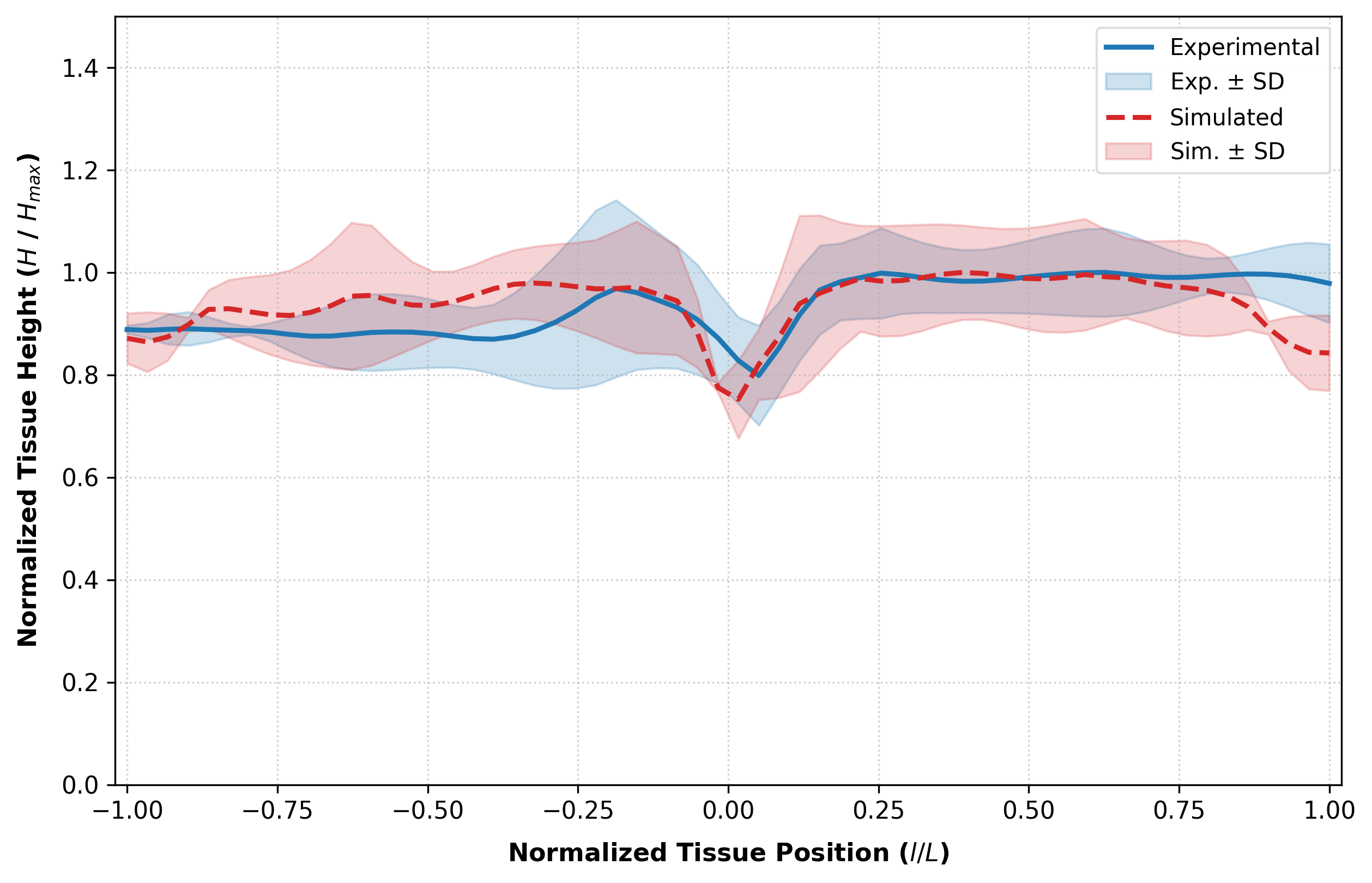

### 4hrSimulatedShape.jpg

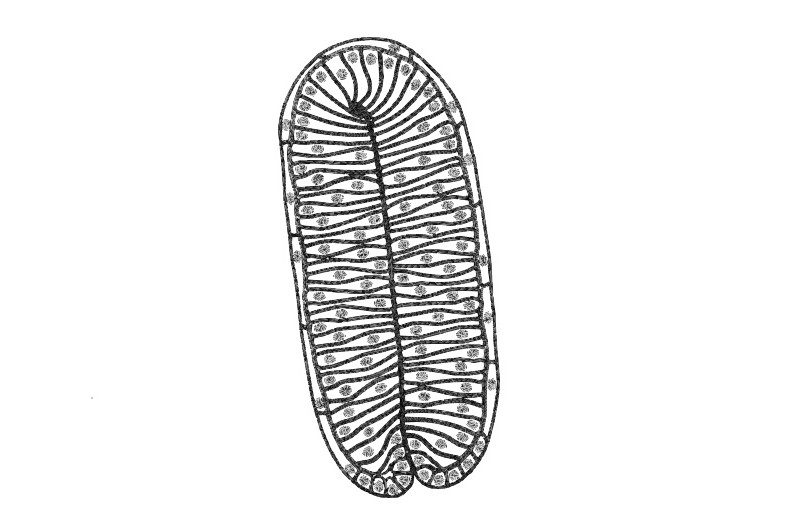

### 4hrSimulatedShape_overlay.jpg

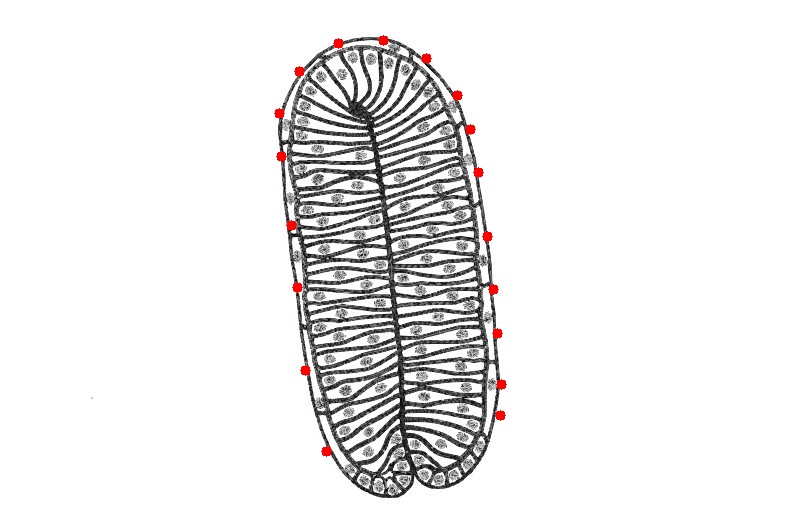

### alt_apical_2hr_curvature_overlay.png

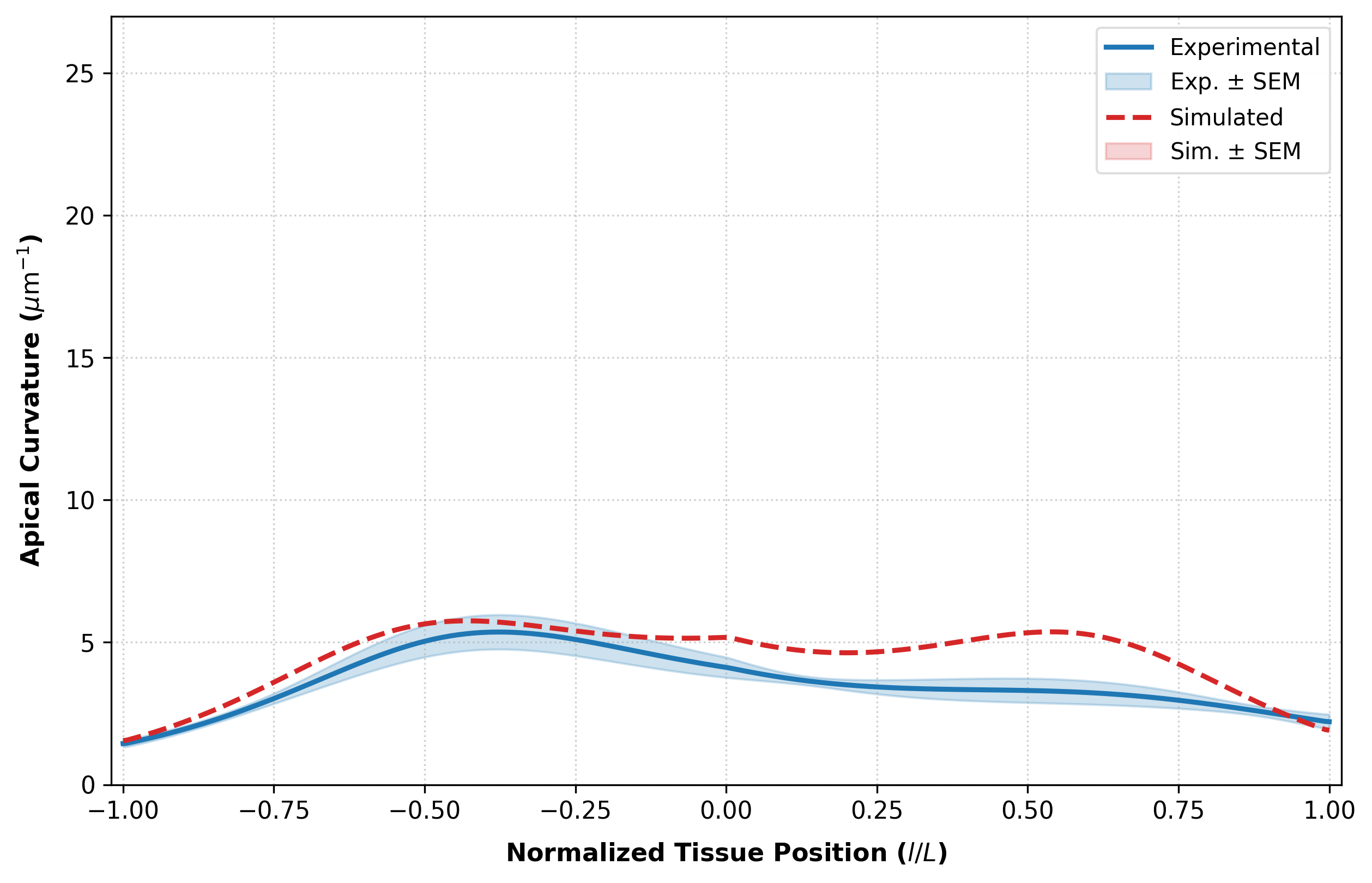

### alt_apical_4hr_curvature_overlay.png

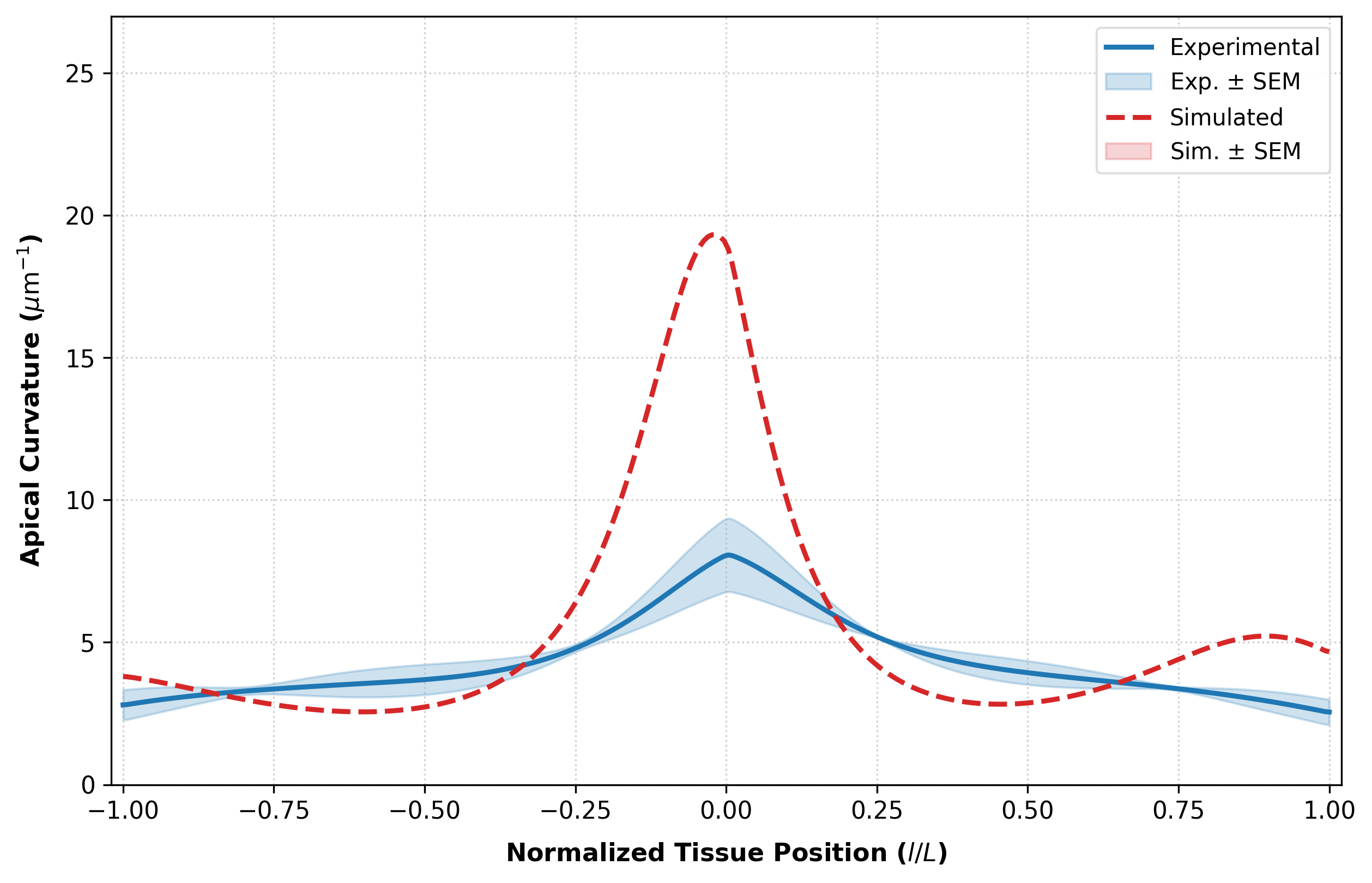

### alt_apical_wL3_curvature_overlay.png

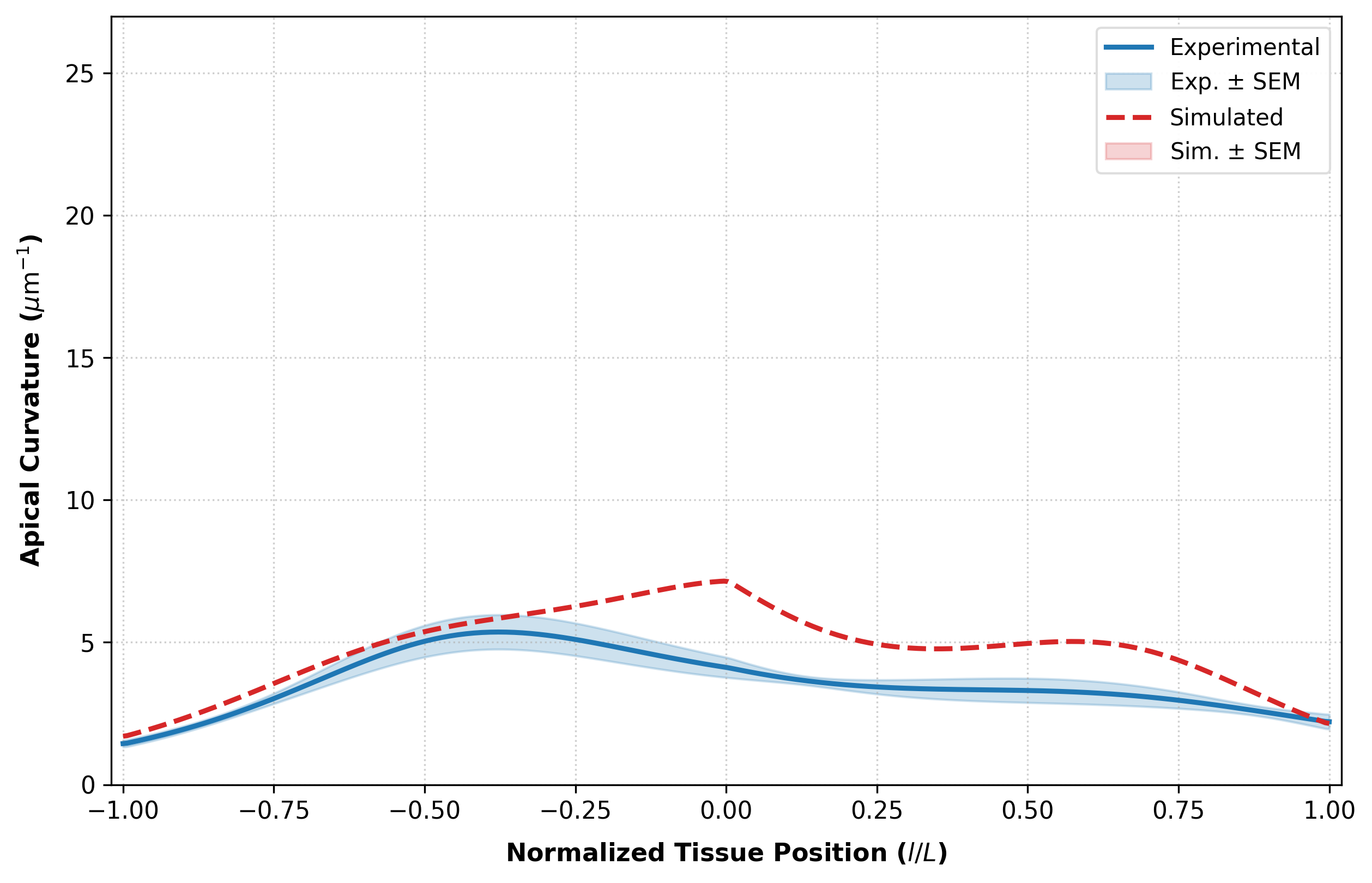

### AltwL3_2hrAPF.jpg

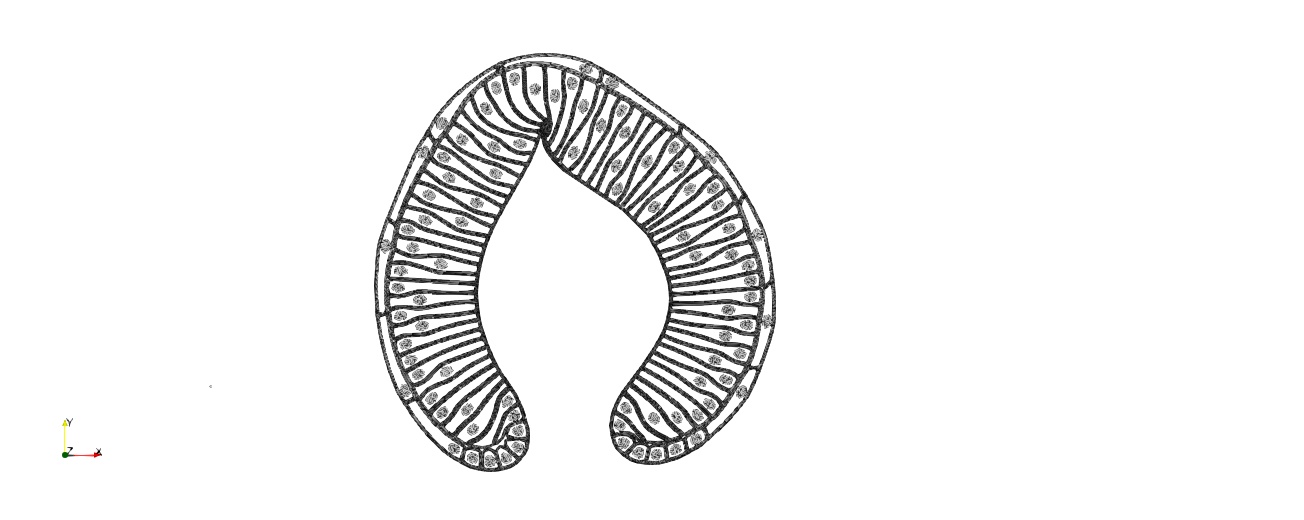

### AltwL3_2hrAPF_overlay.jpg

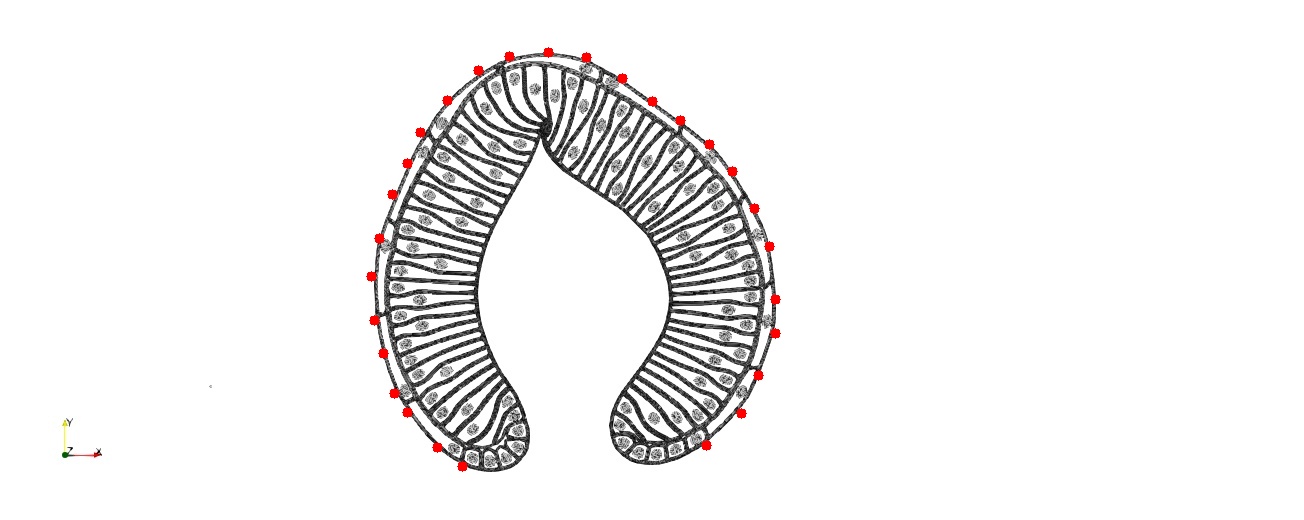

### AltwL3_2hrT100.jpg

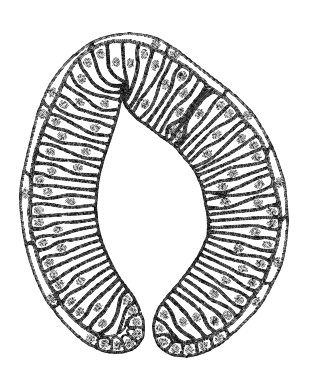

### AltwL3_4hrAPF.jpg

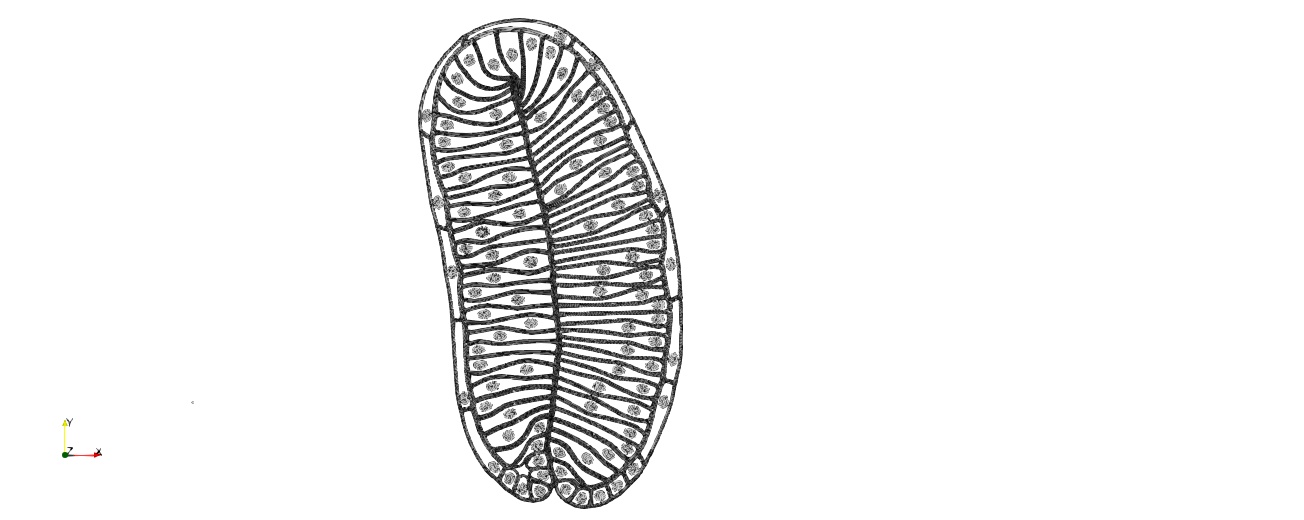

### altwL3_basal_2hr_curvature_overlay.png

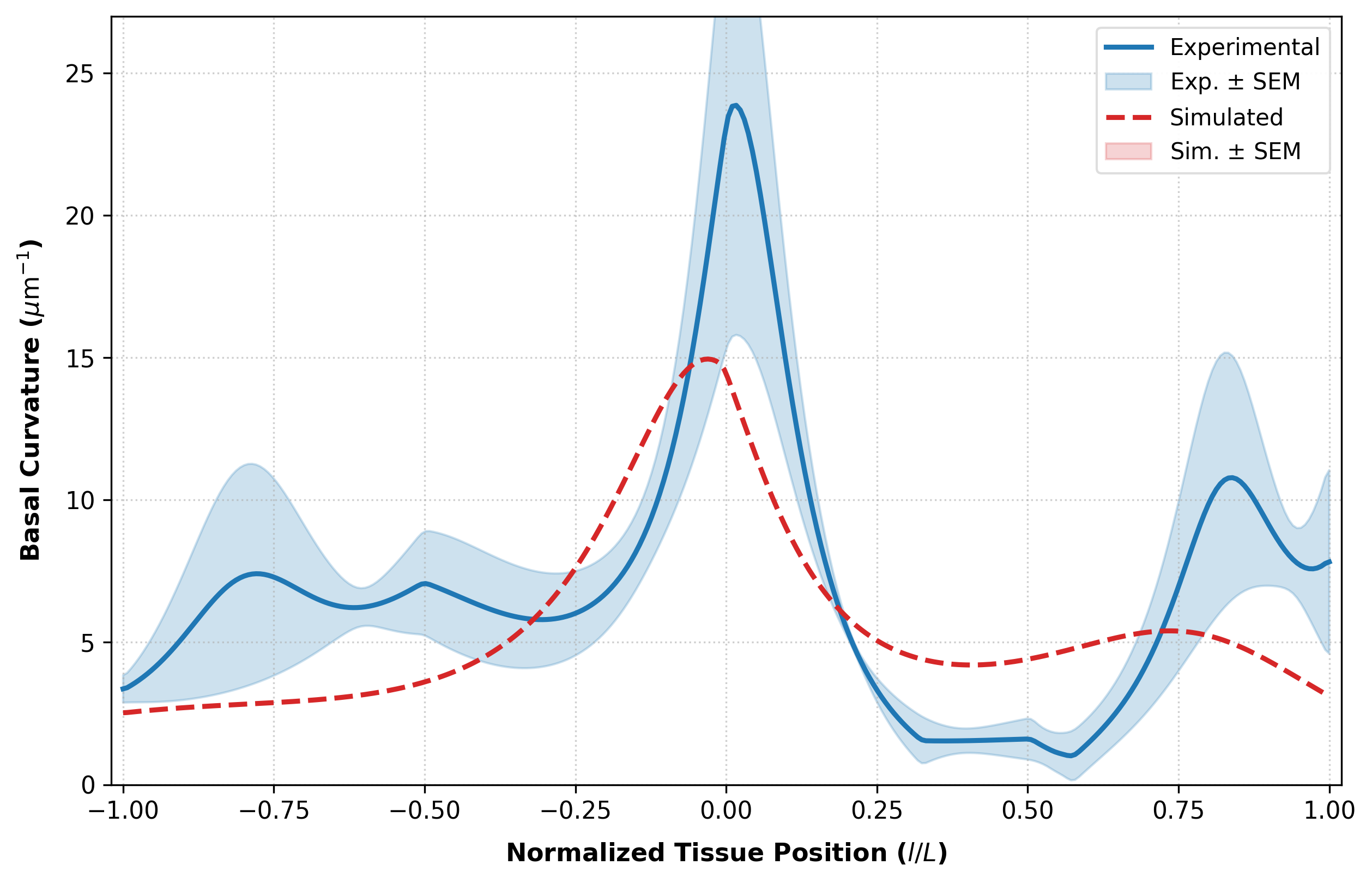

### altwL3_basal_4hr_curvature_overlay.png

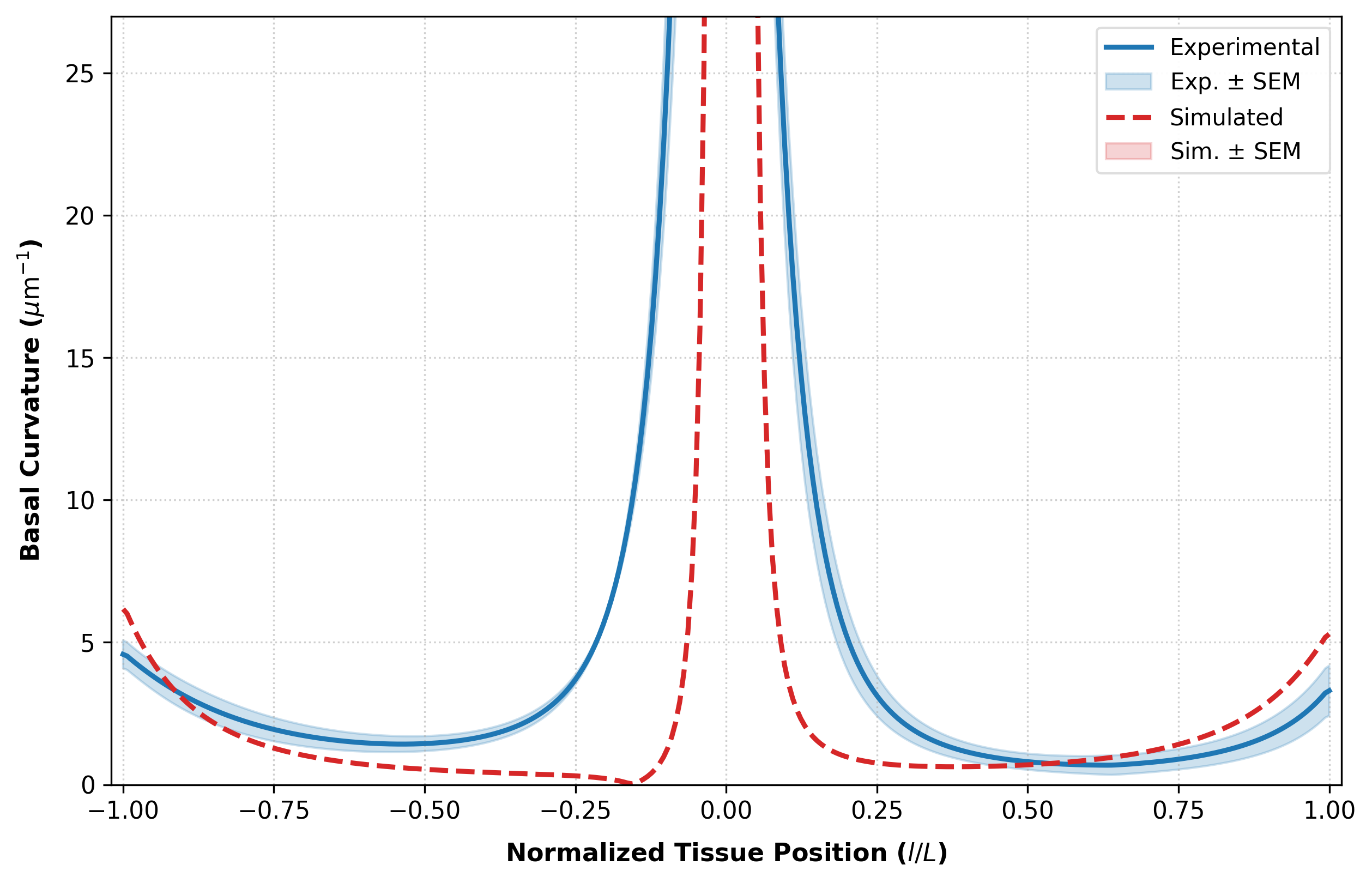

### altwL3_basal_wL3_curvature_overlay.png

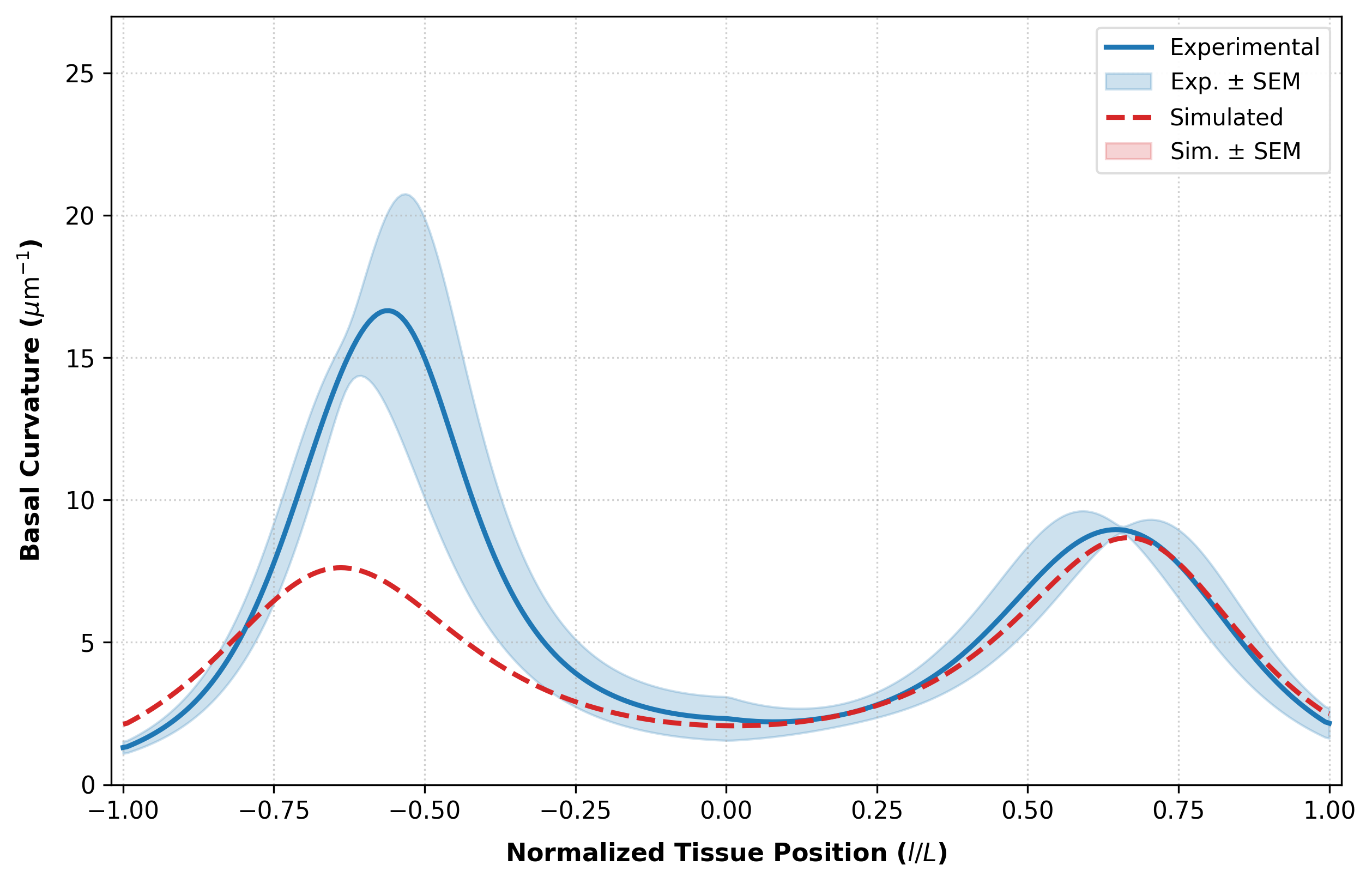

### AltwL3InitialShape.jpg

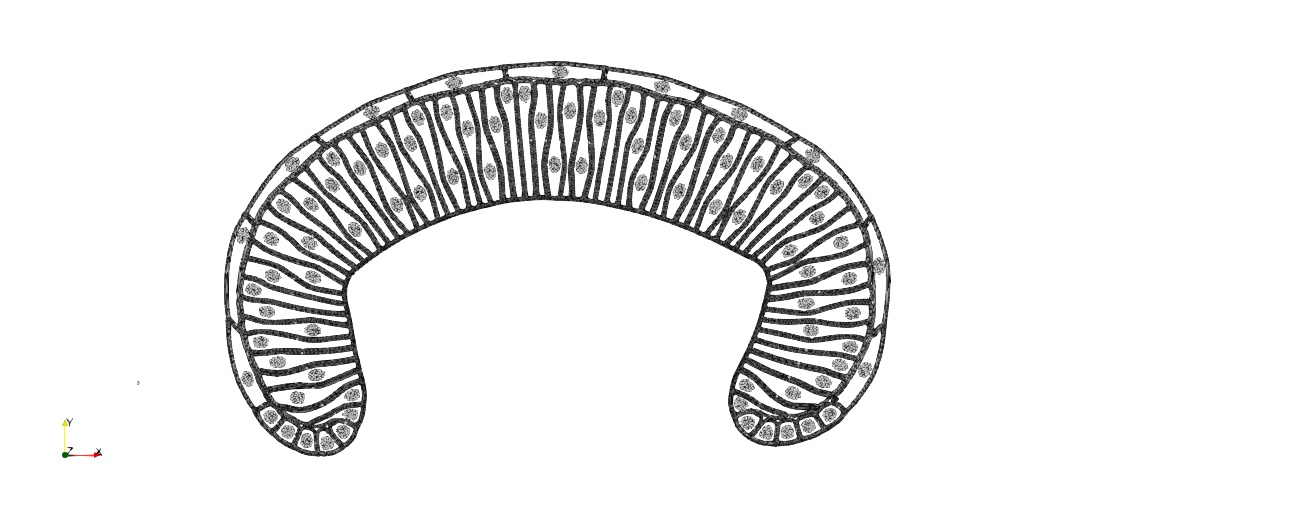

### Apical Surface Length 0hAPF.jpeg

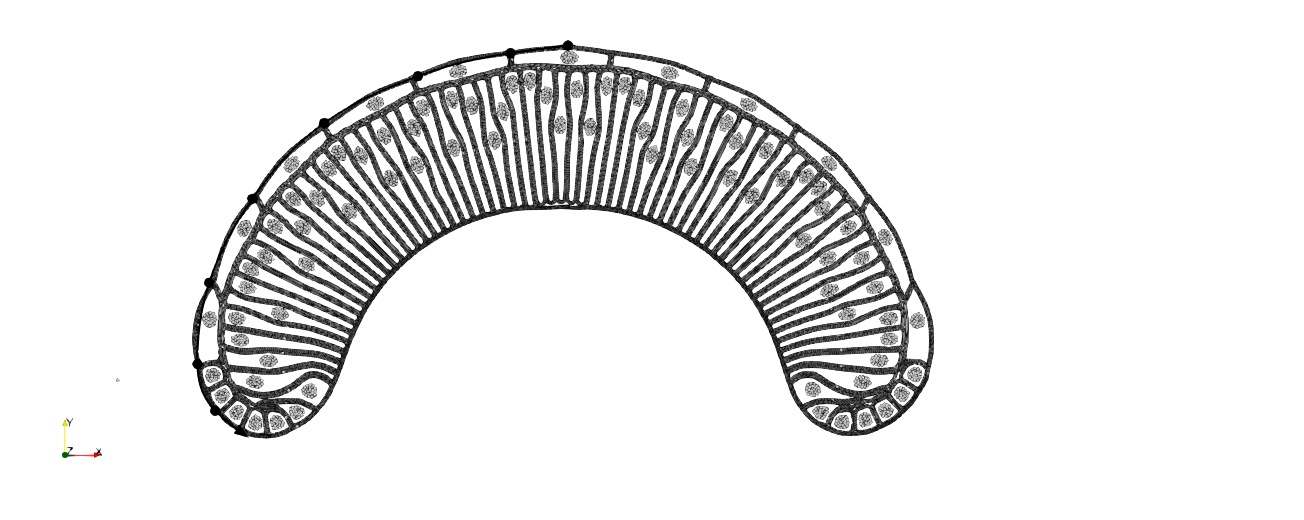

### Apical Surface Length 2hAPF.jpeg

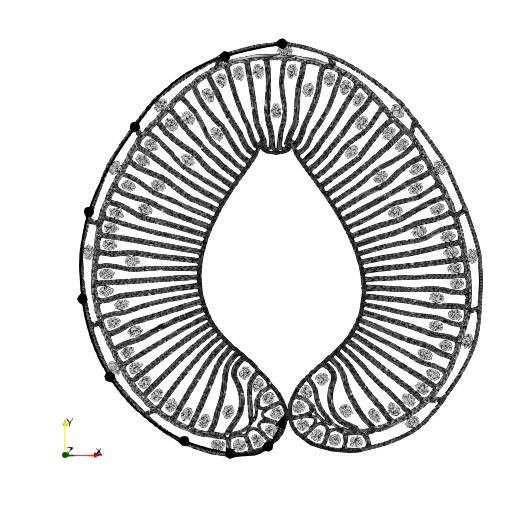

### Apical Surface Length 4hAPF.jpeg

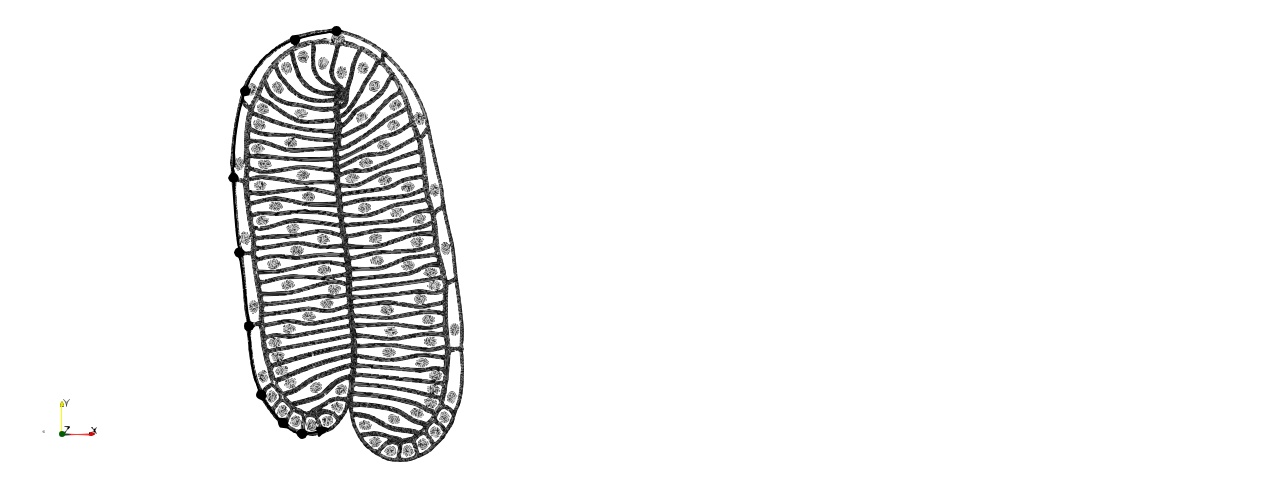

### Apical Surface Length wL3.jpeg

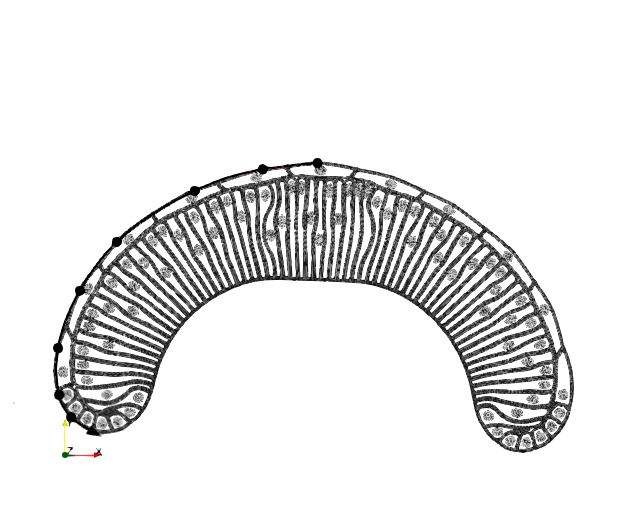

### Apical Surface Lengths Plotted.png

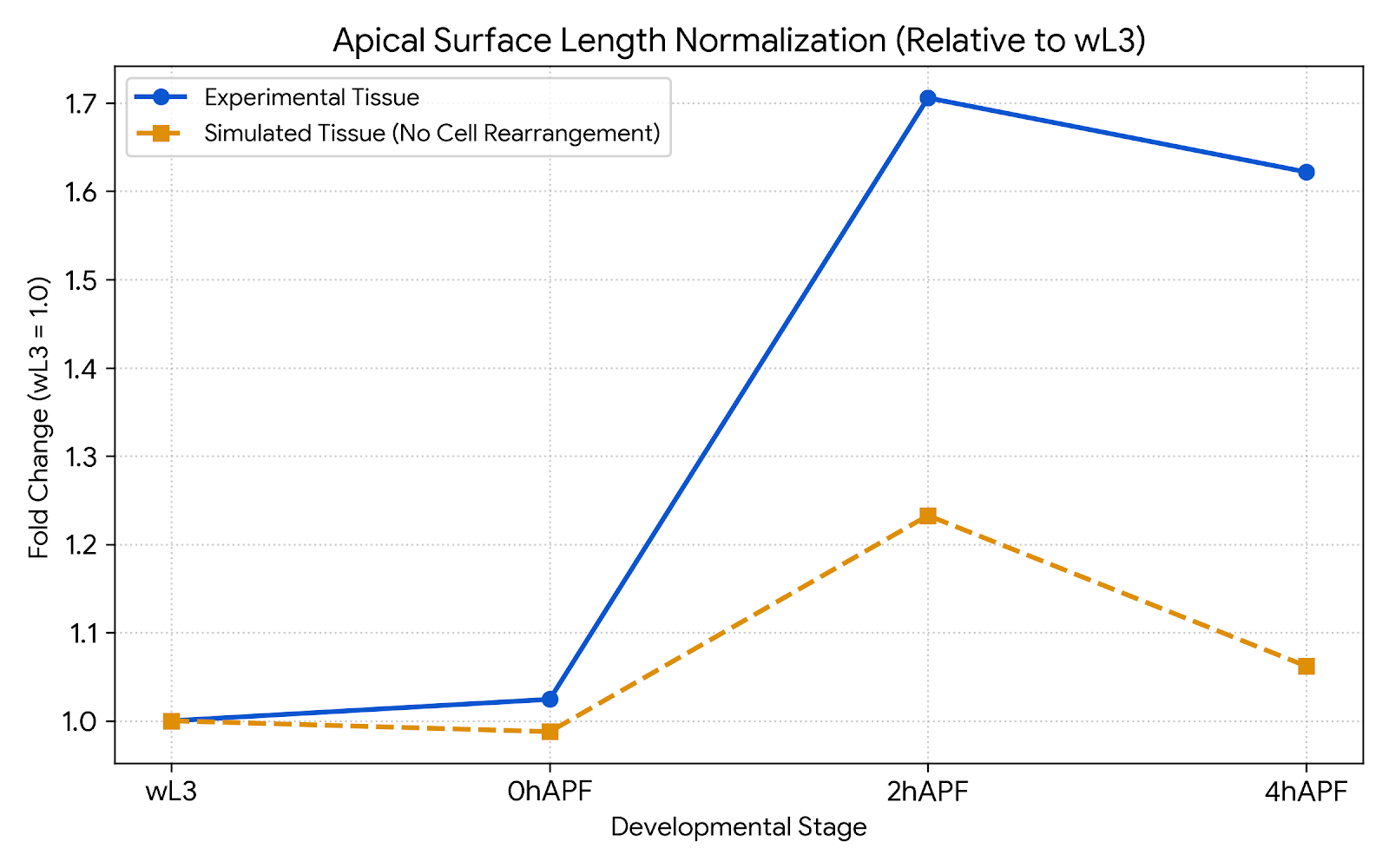

### apical_2hr_curvature_overlay.png

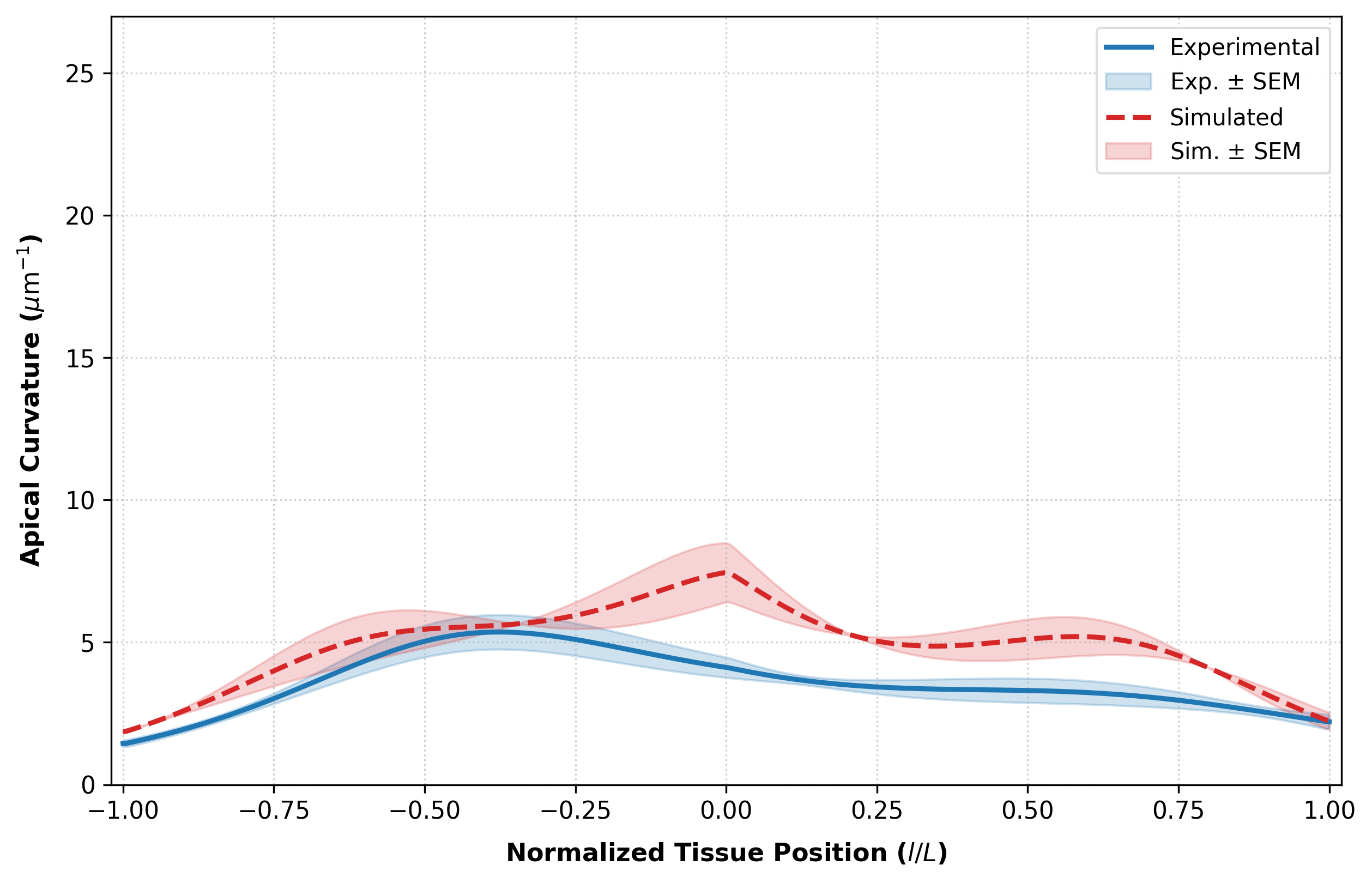

### apical_4hr_curvature_overlay.png

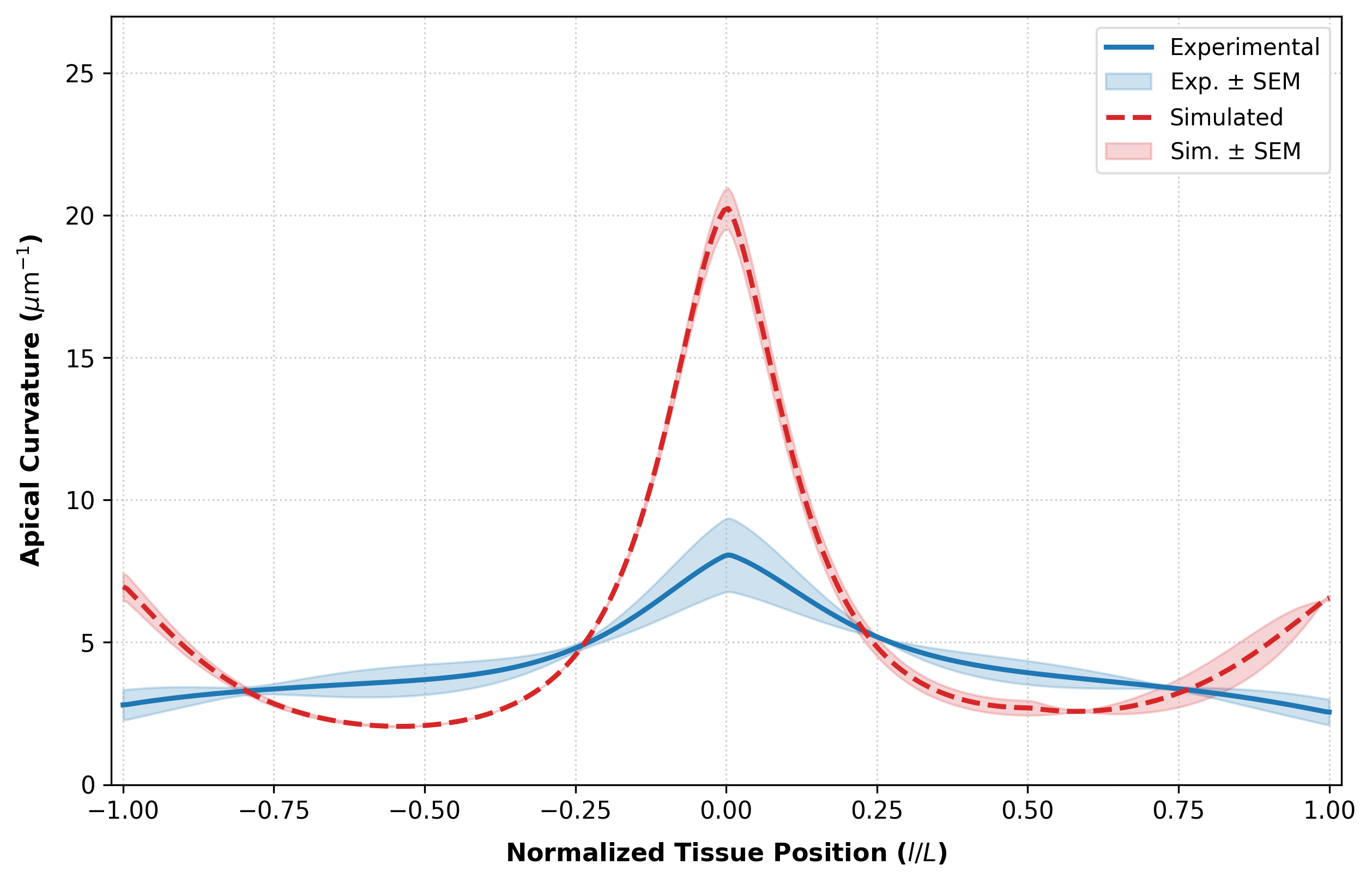

### apical_surface_lengths_normalized.png

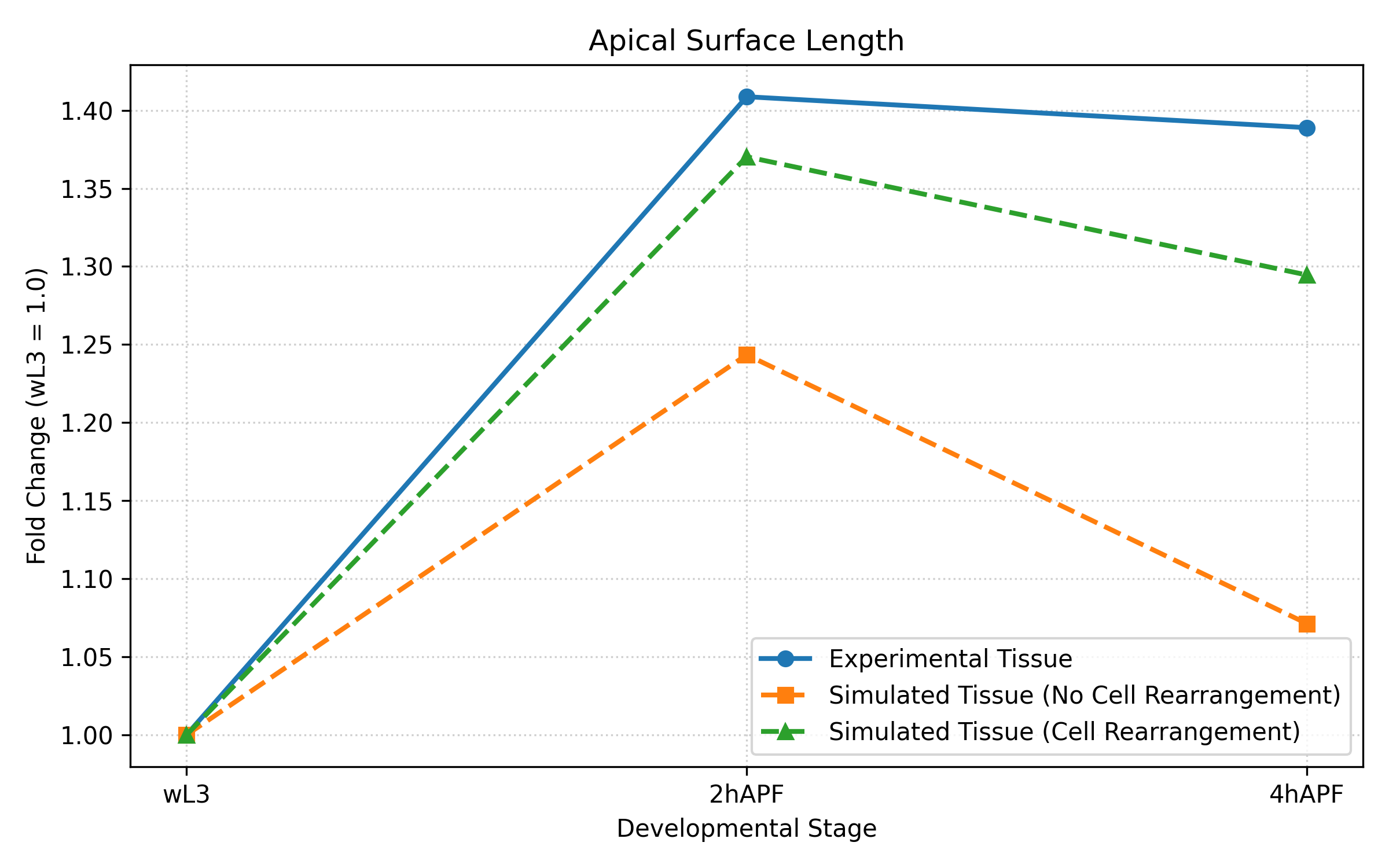

### apical_wL3_curvature_overlay.png

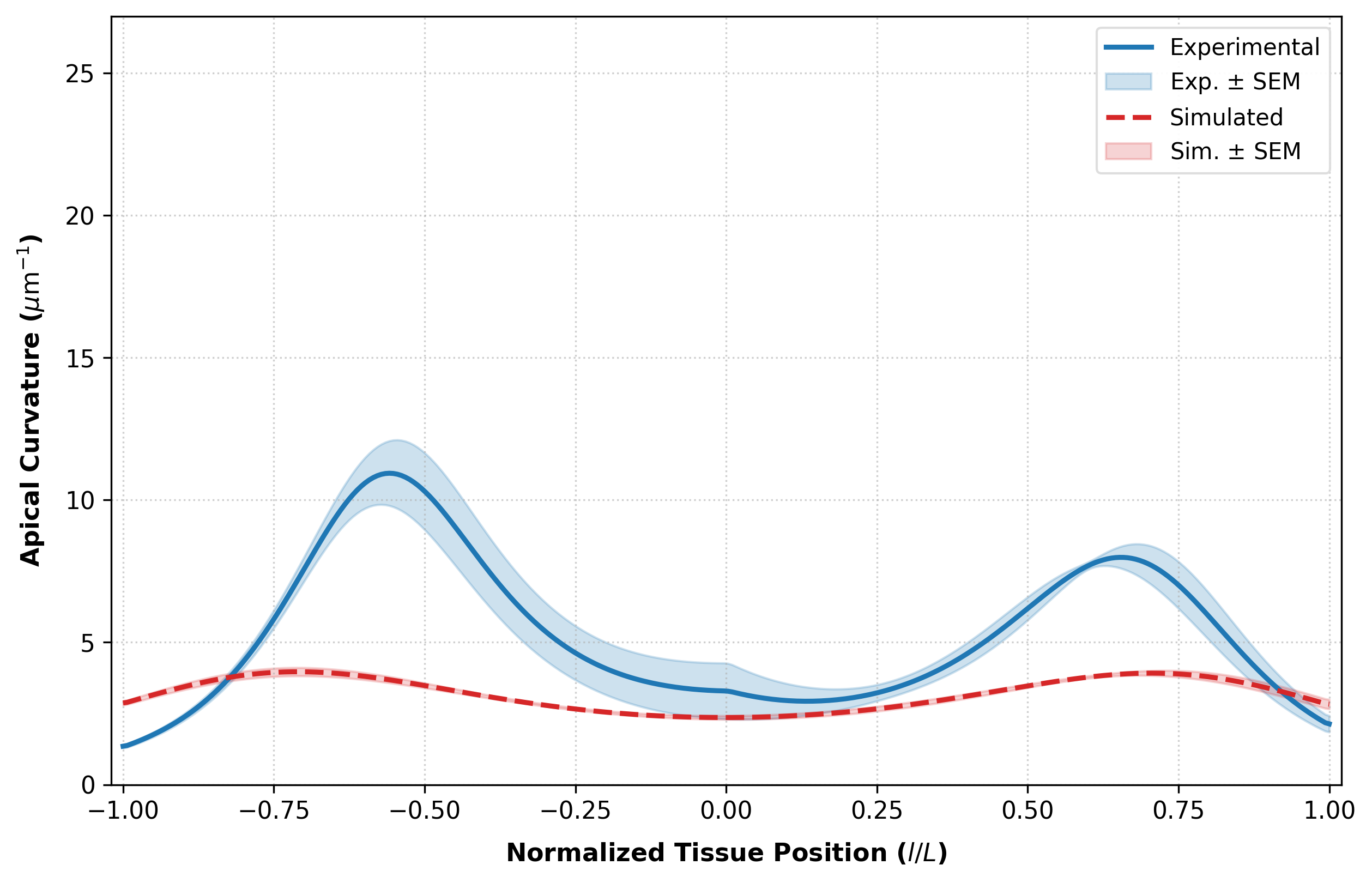

### basal_2hr_curvature_overlay.png

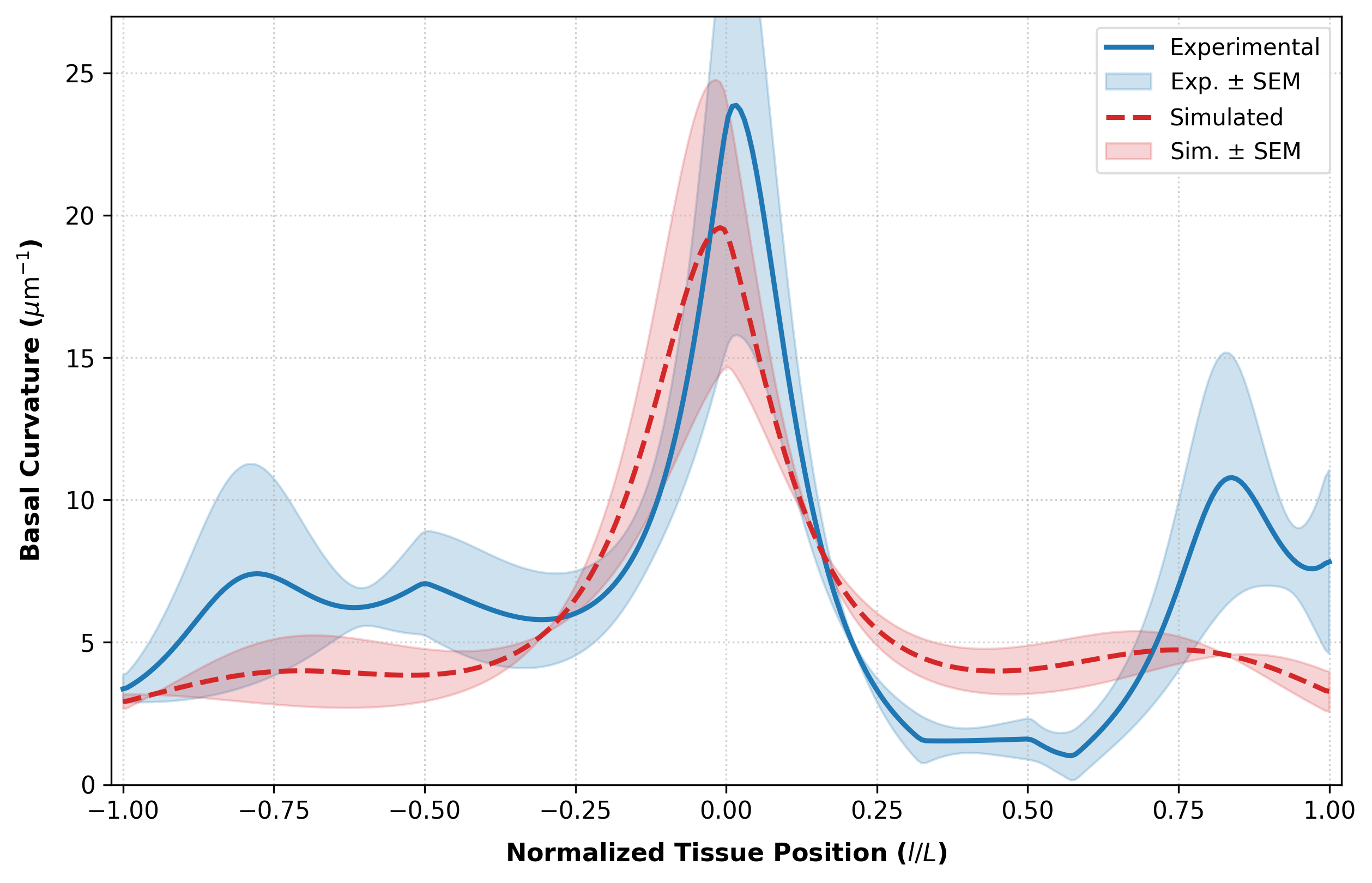

### basal_4hr_curvature_overlay.png

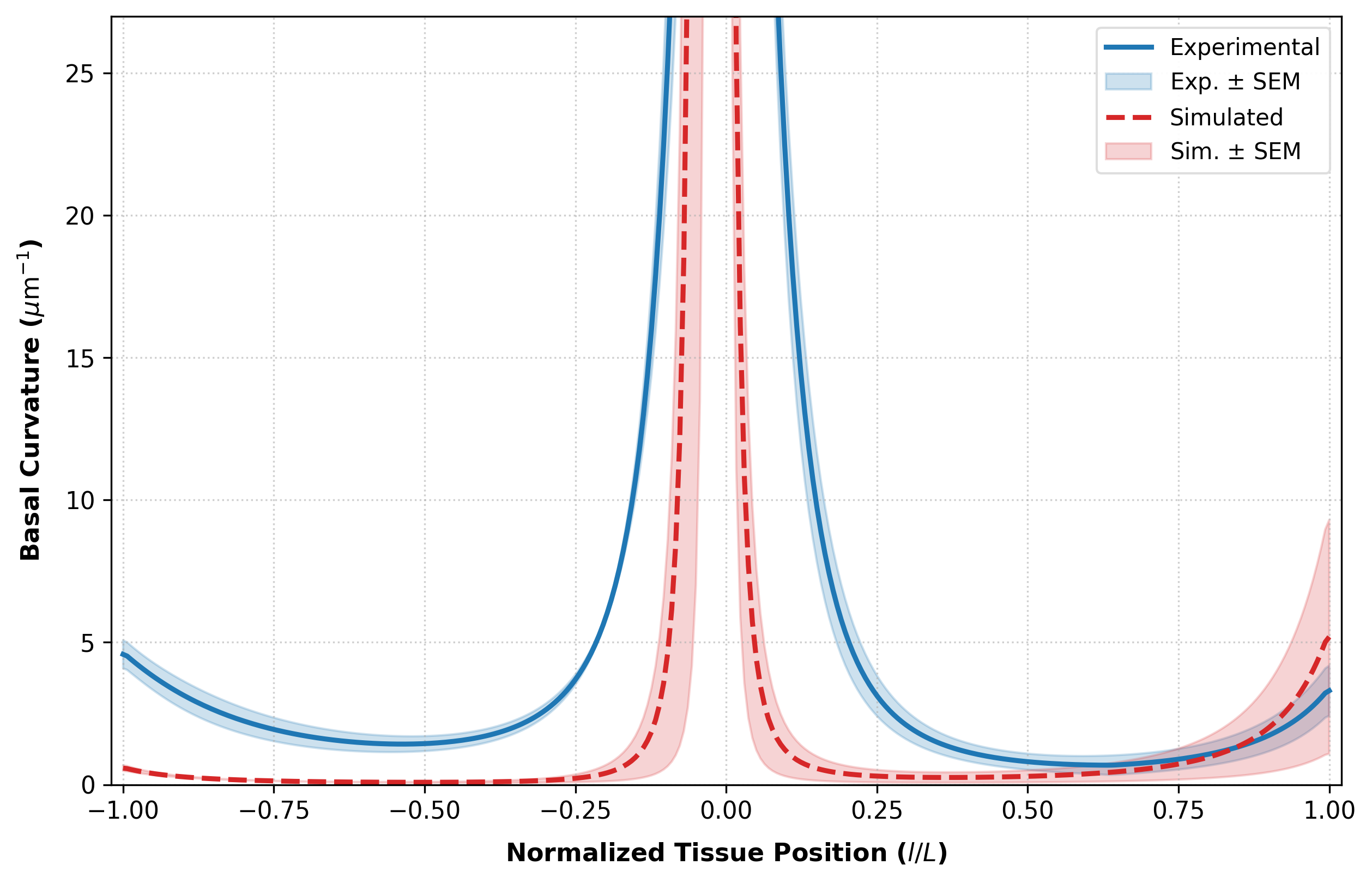

### basal_surface_lengths_normalized.png

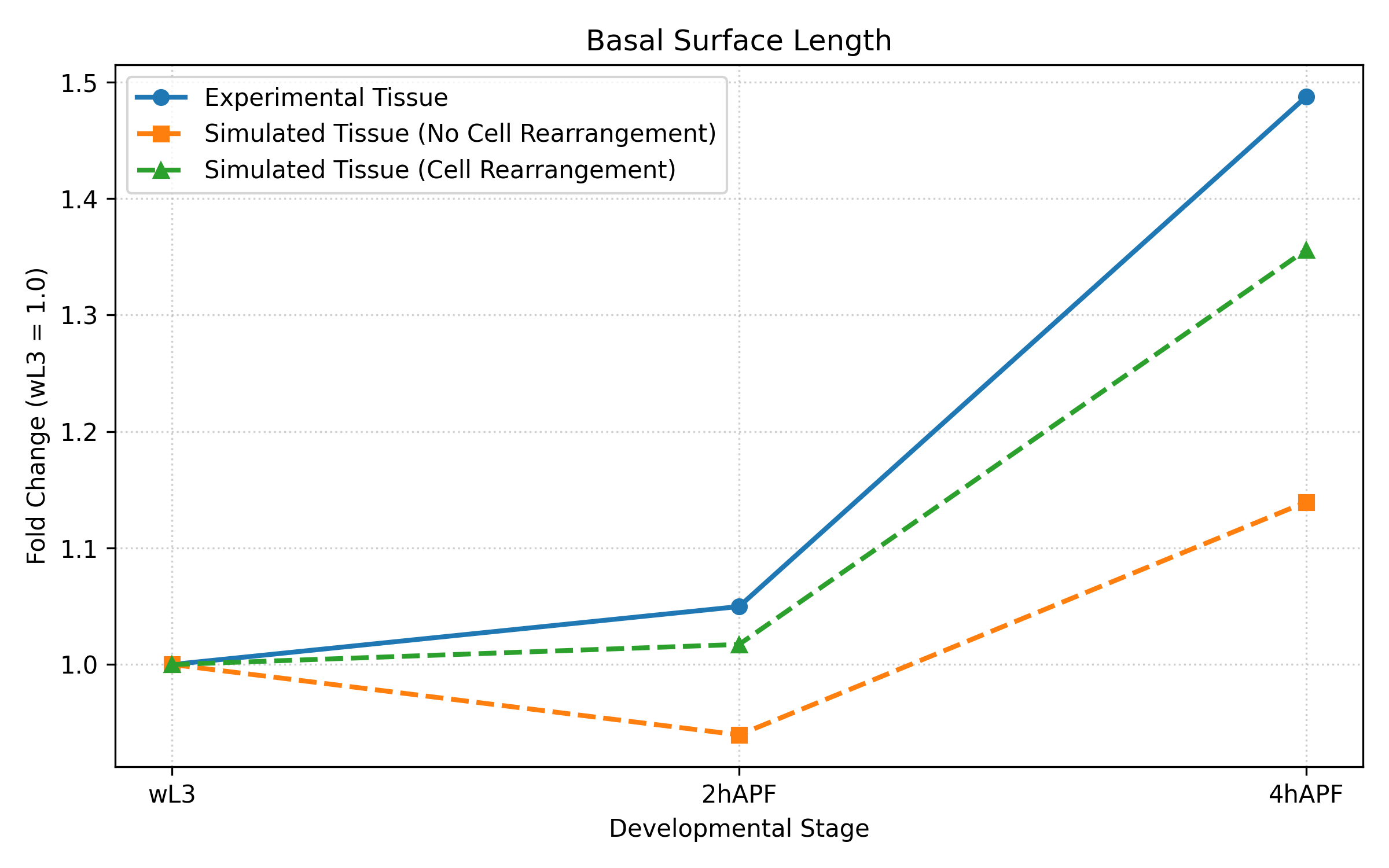

### basal_wL3_curvature_overlay.png

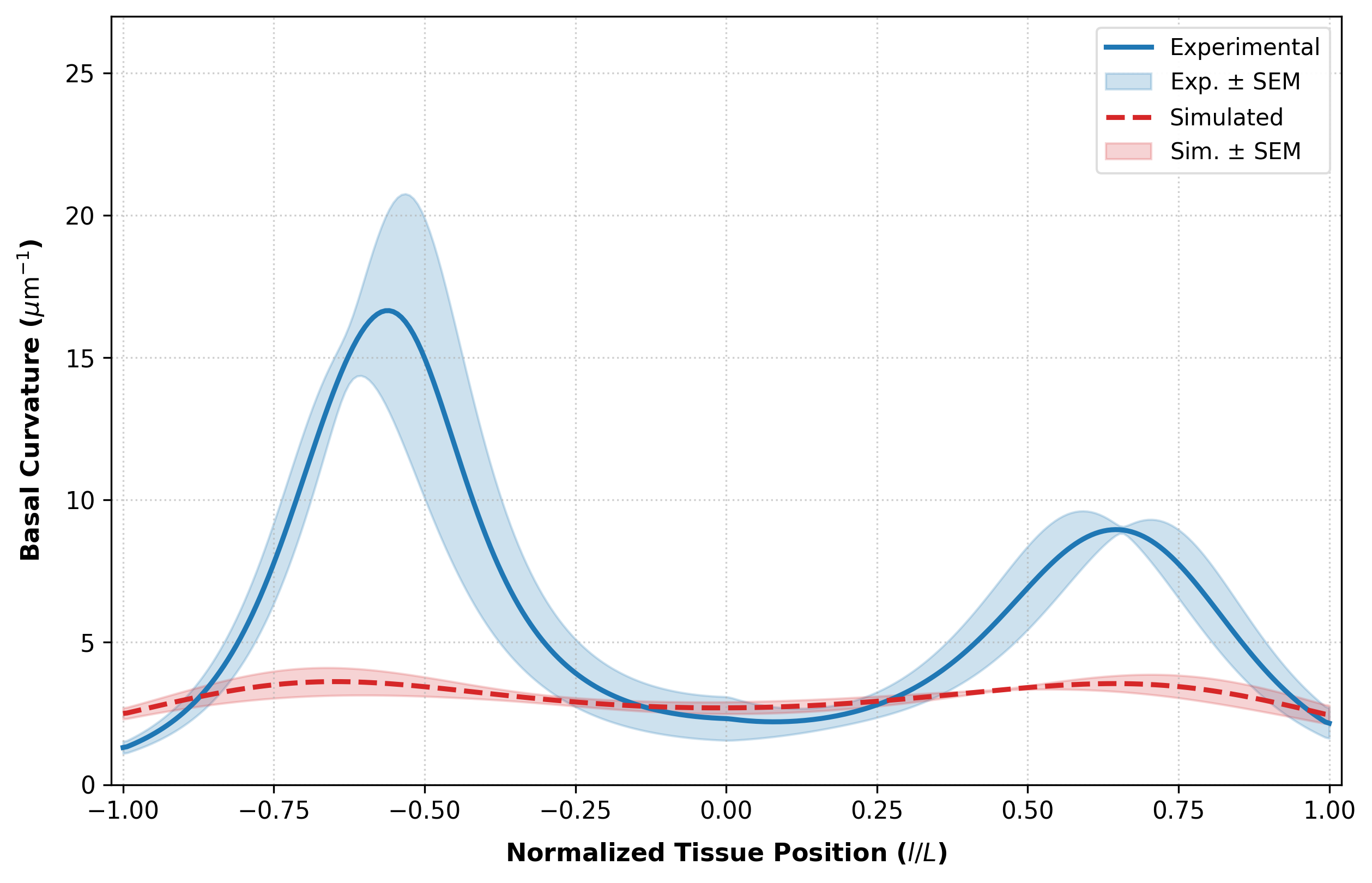

### D1.jpg

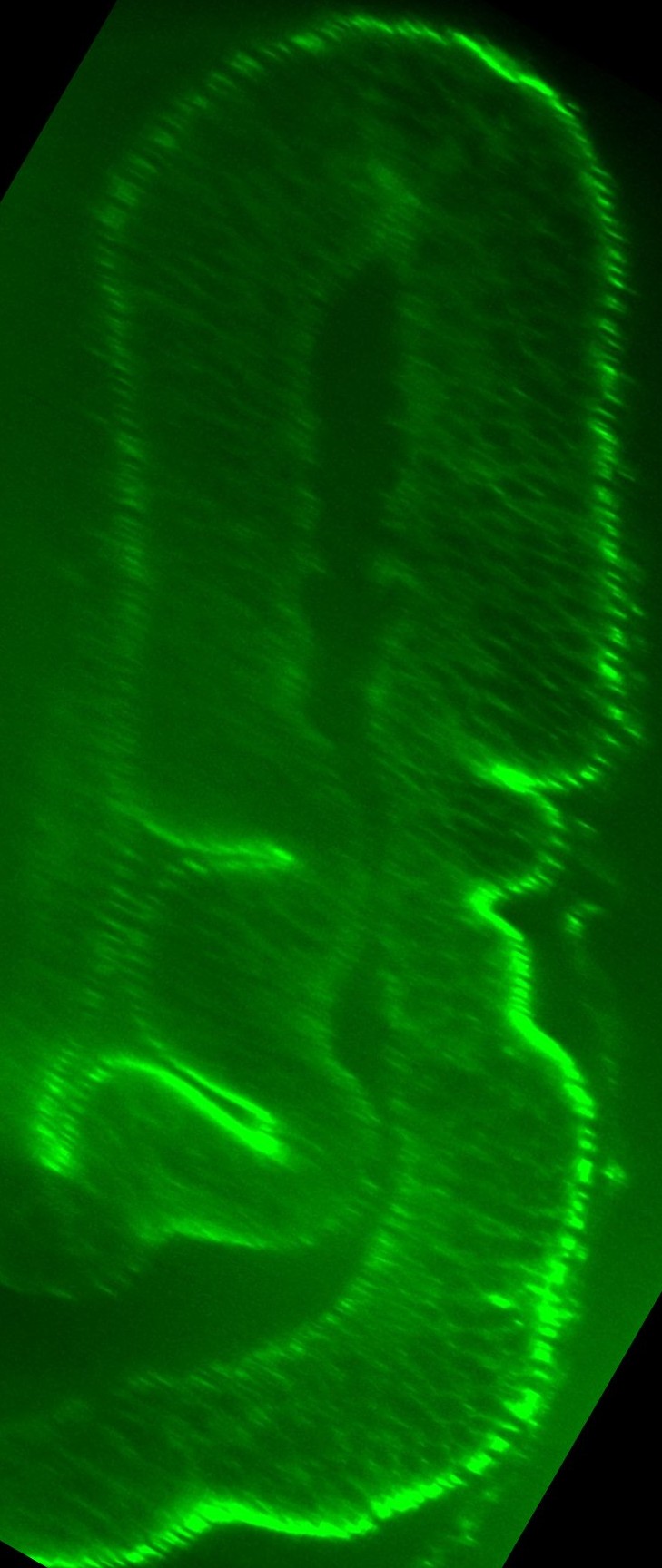

### D1.jpg

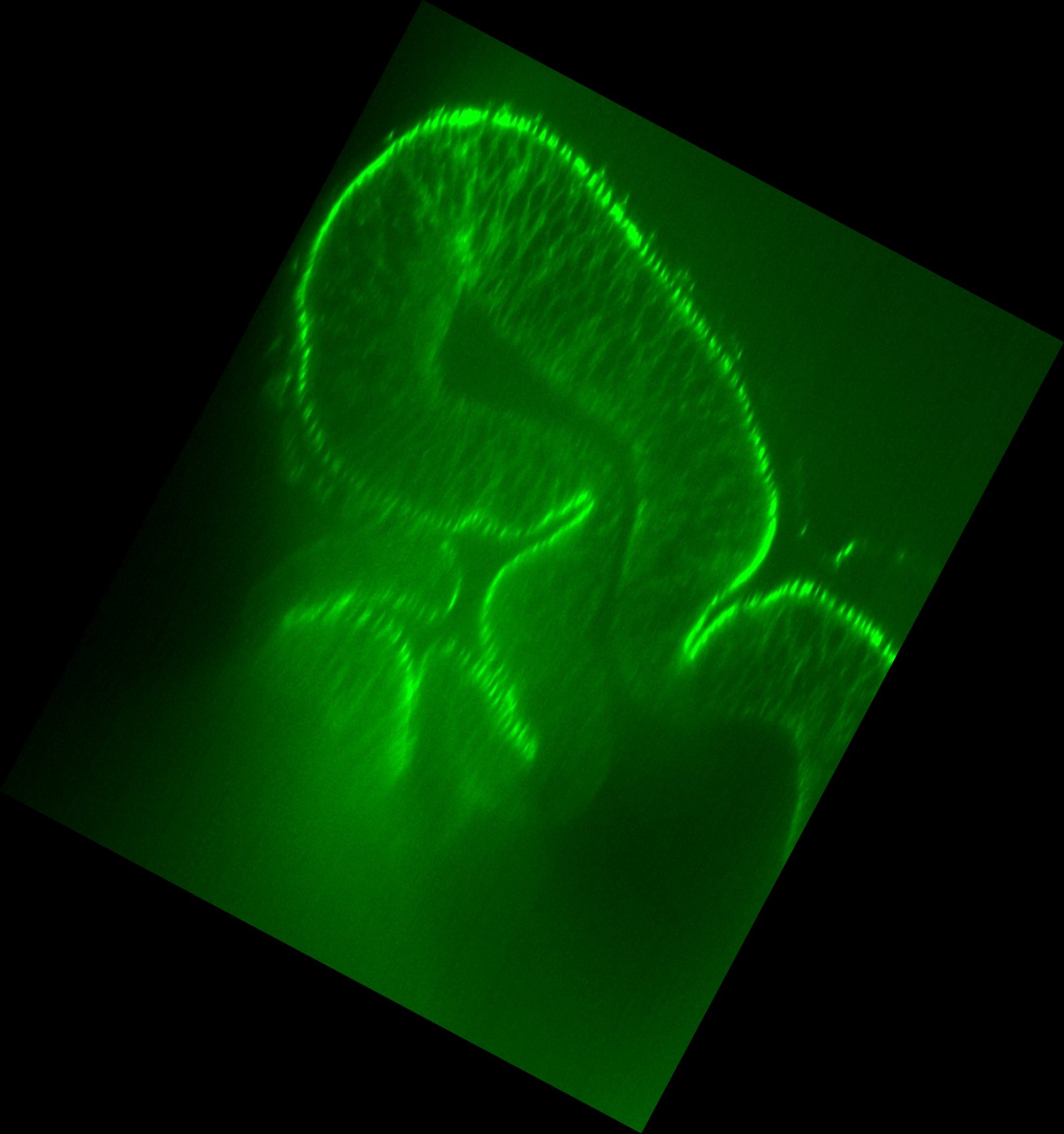

### D1.jpg

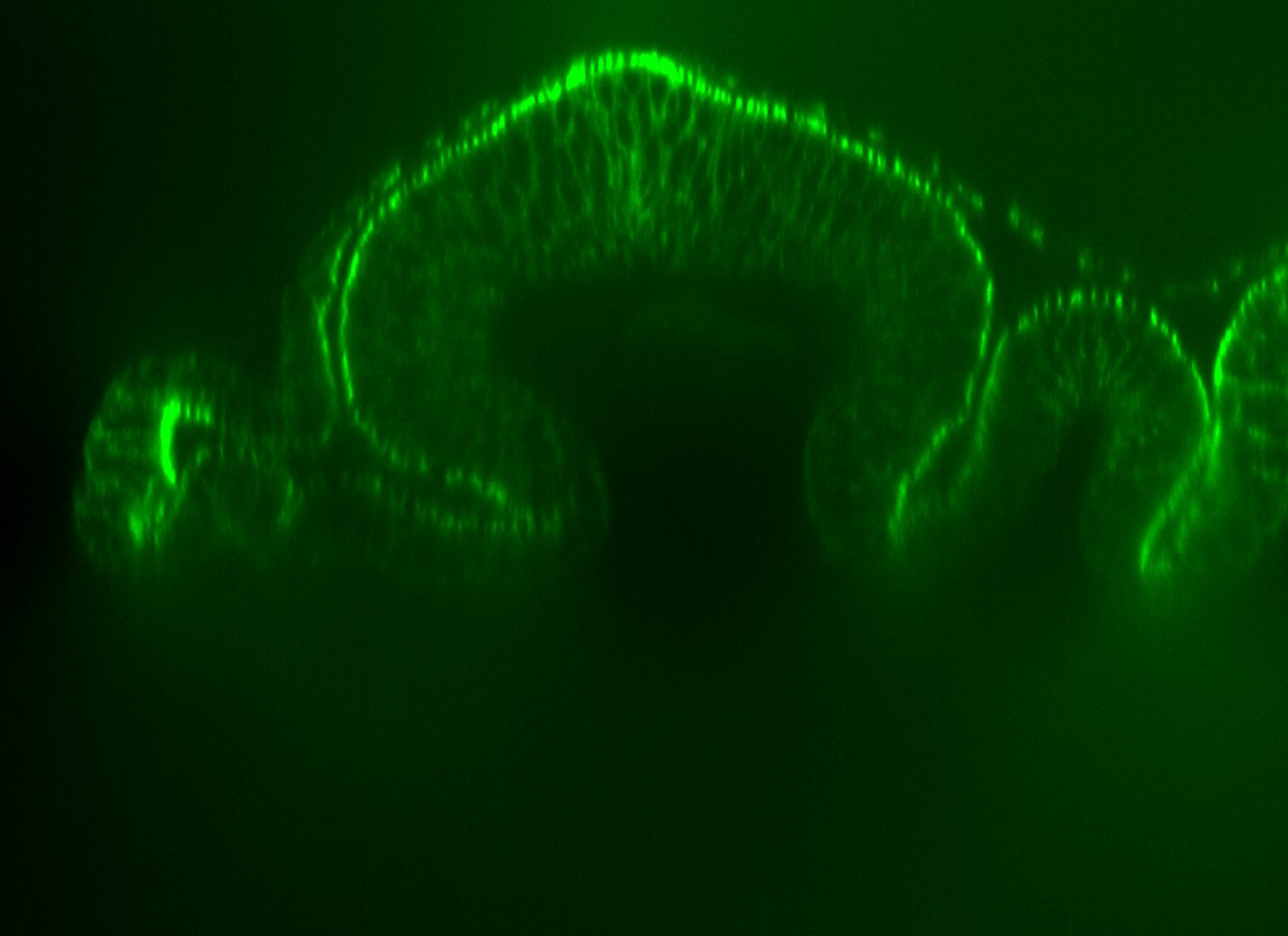

### D1.jpg

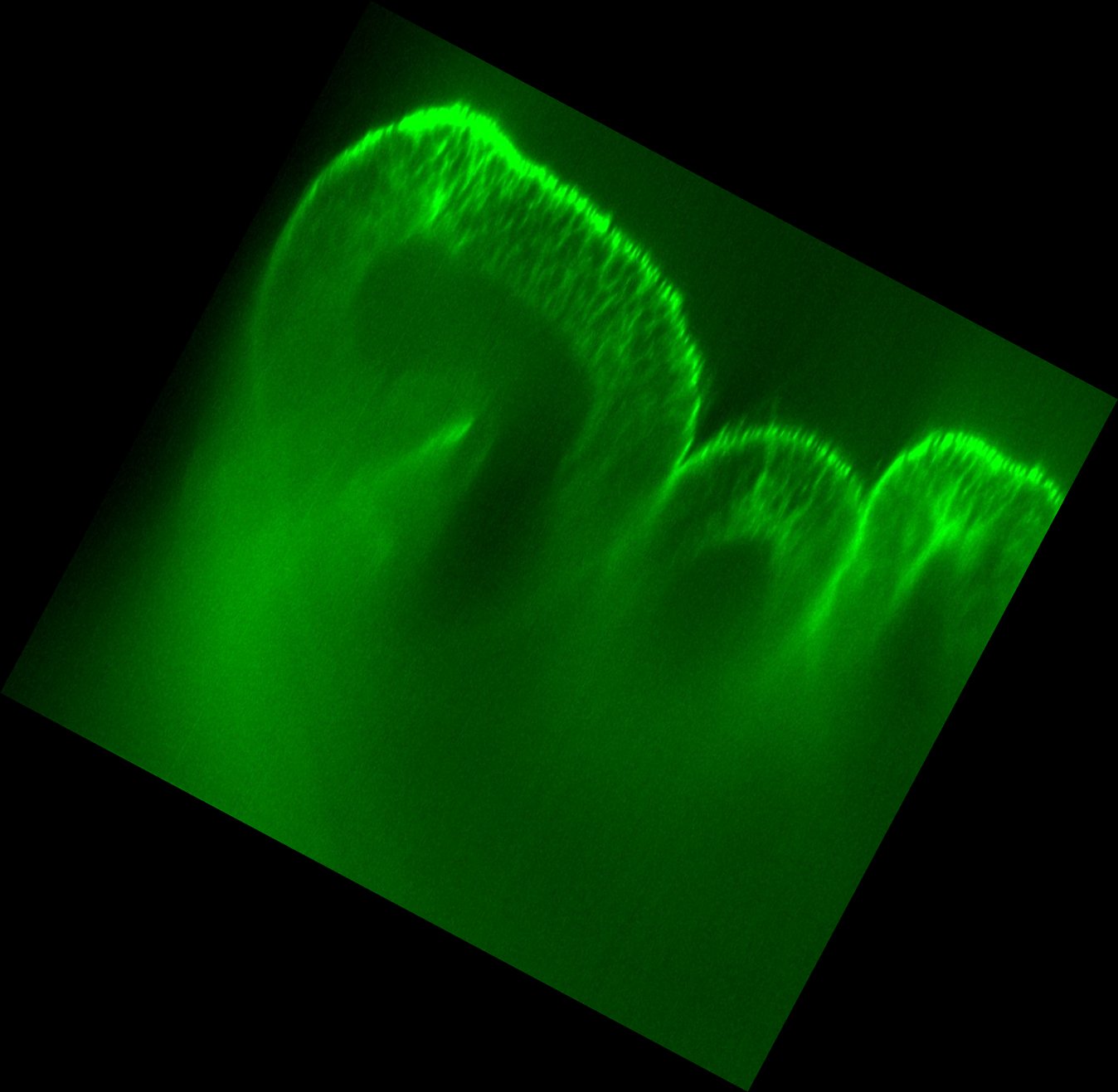

### D1_2.jpg

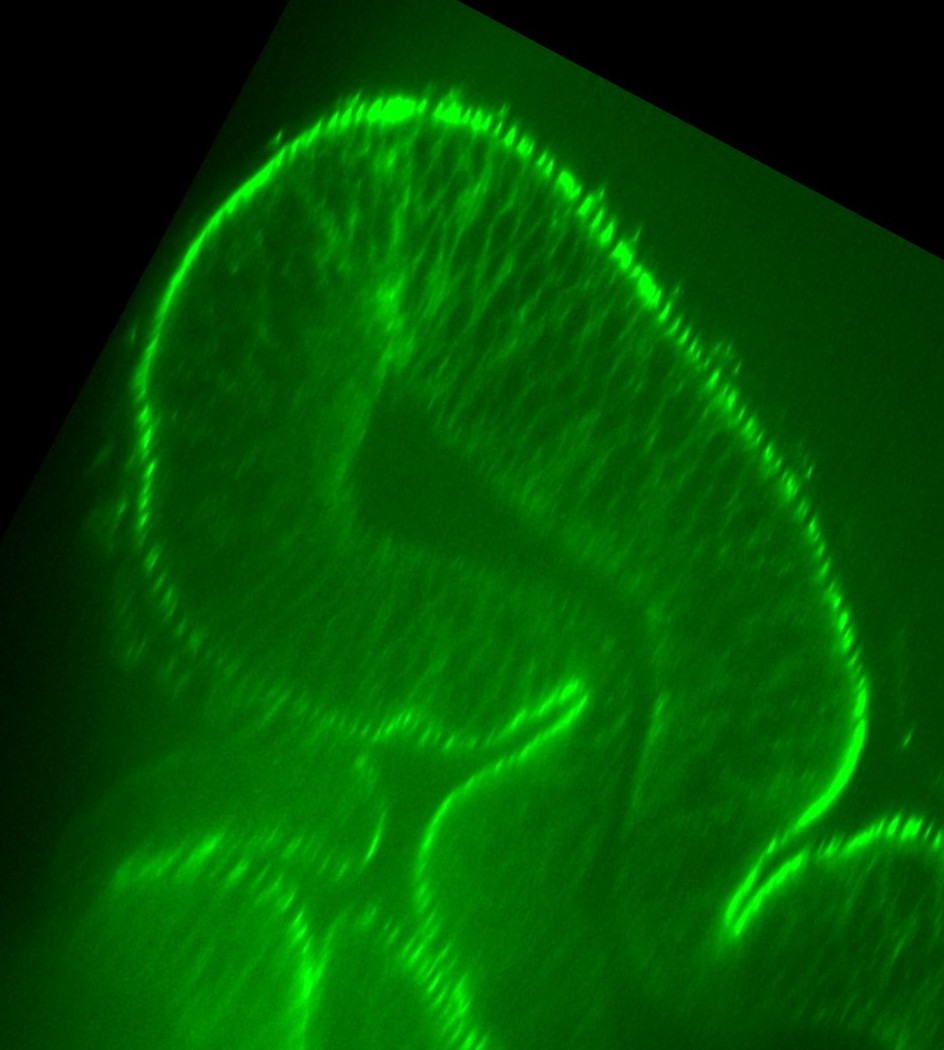

### D1Traced.jpg

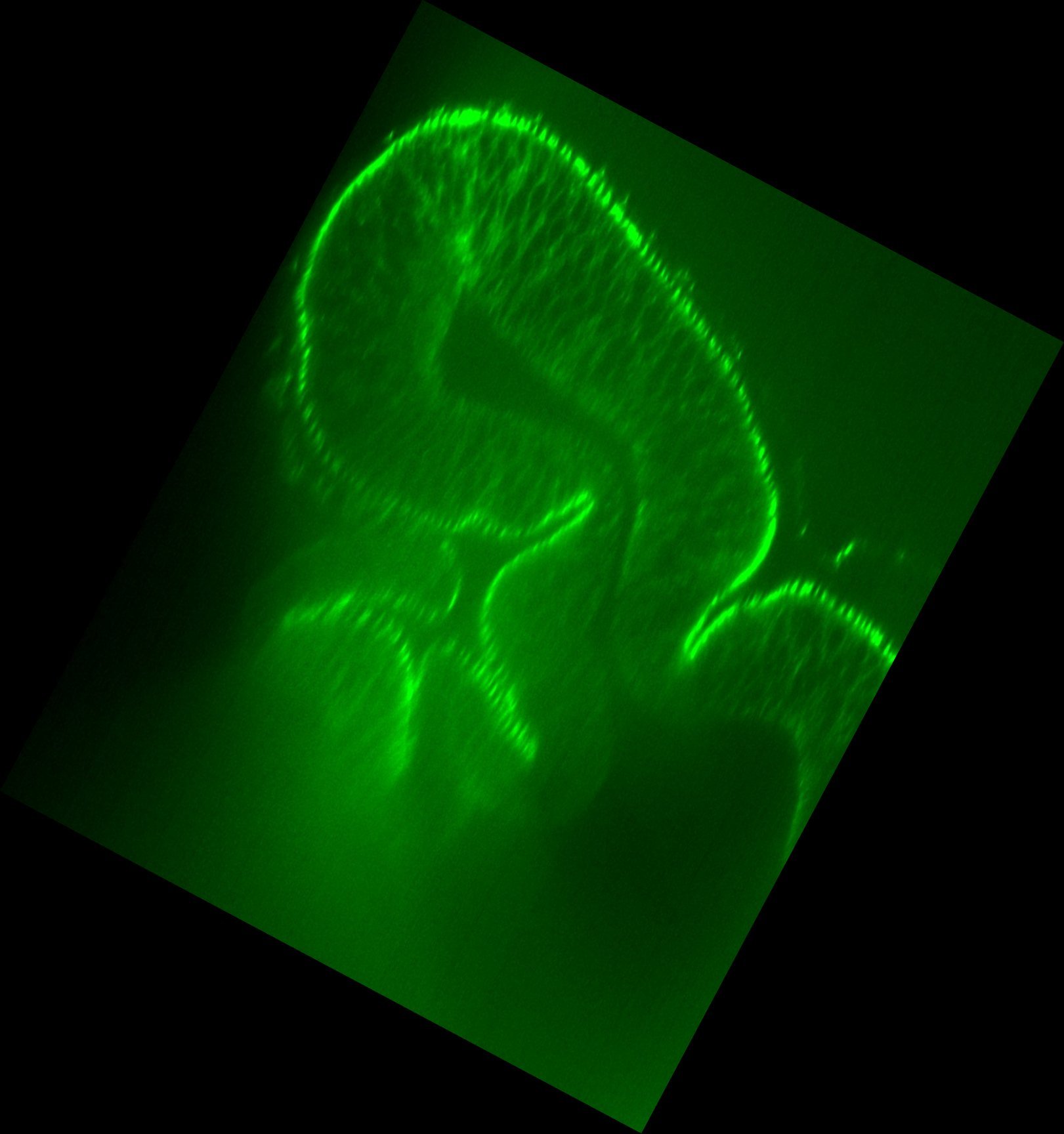

### D1Traced.jpg

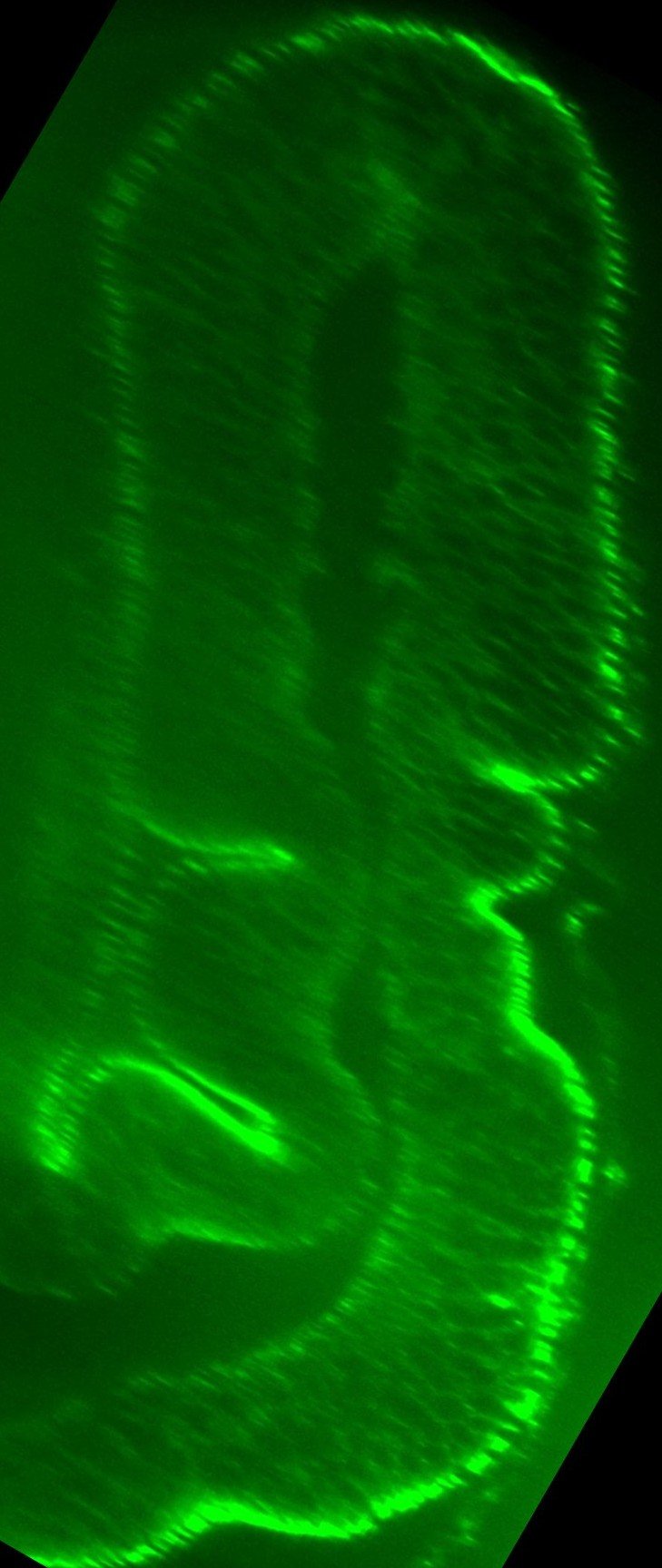

### D1Traced.jpg

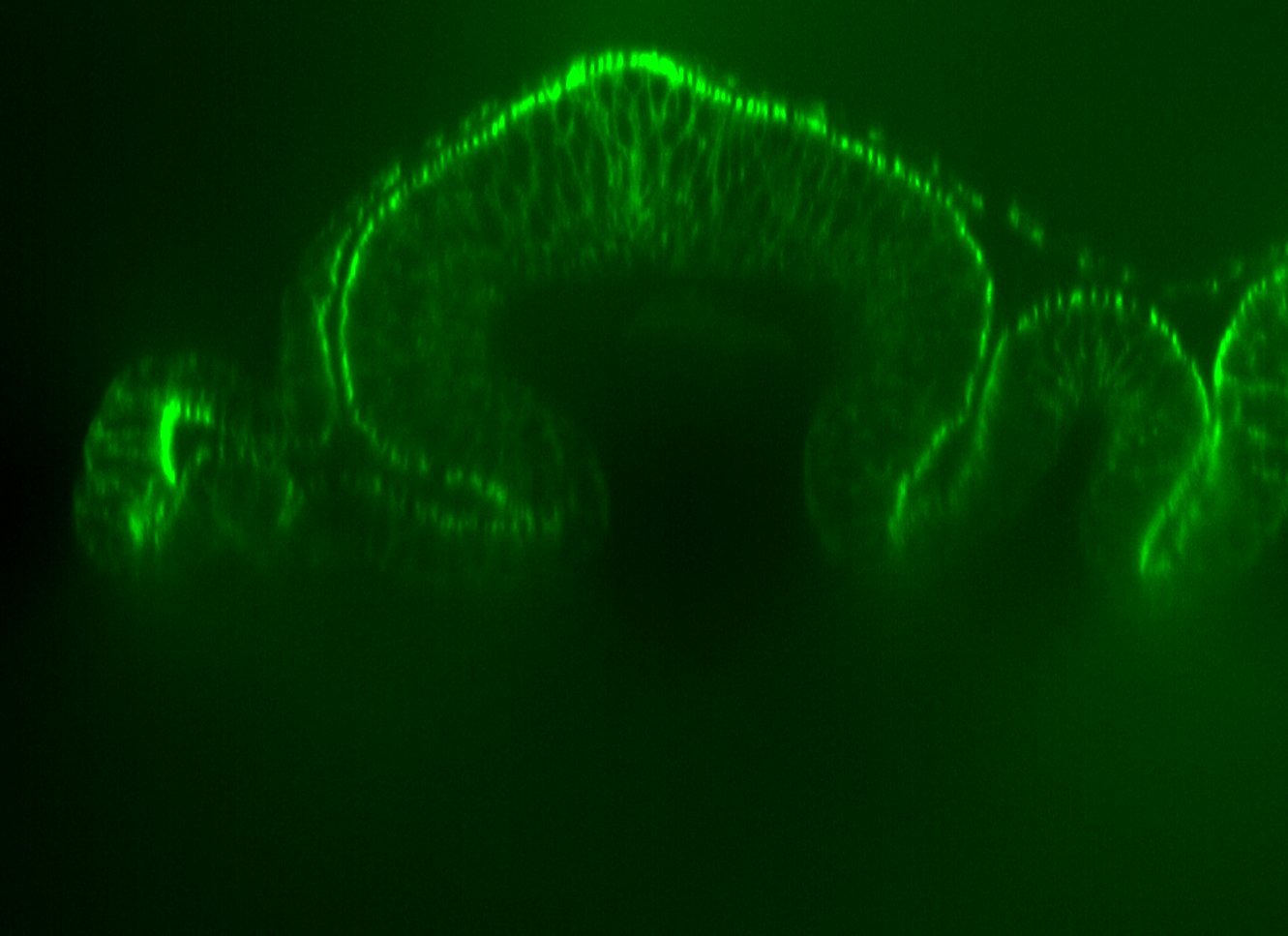

### D2.jpg

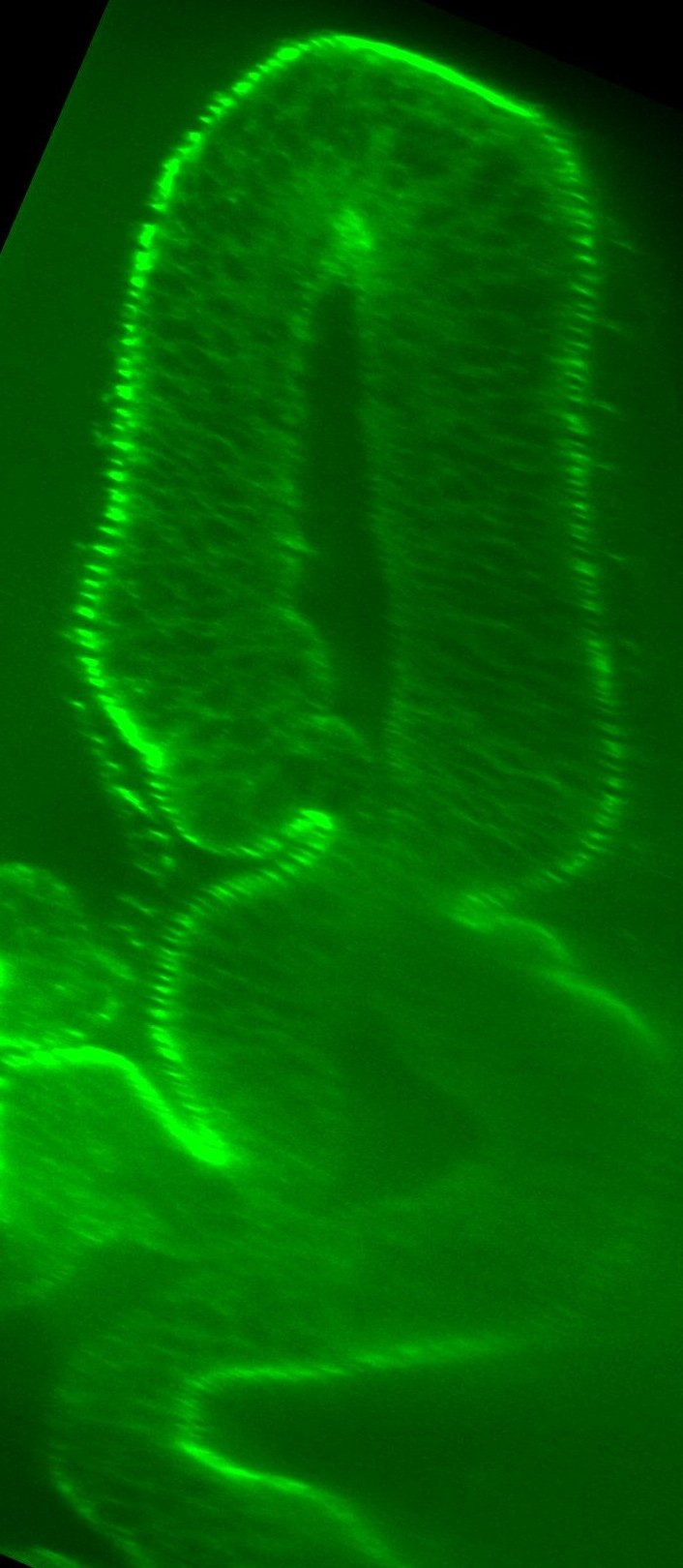

### D2.jpg

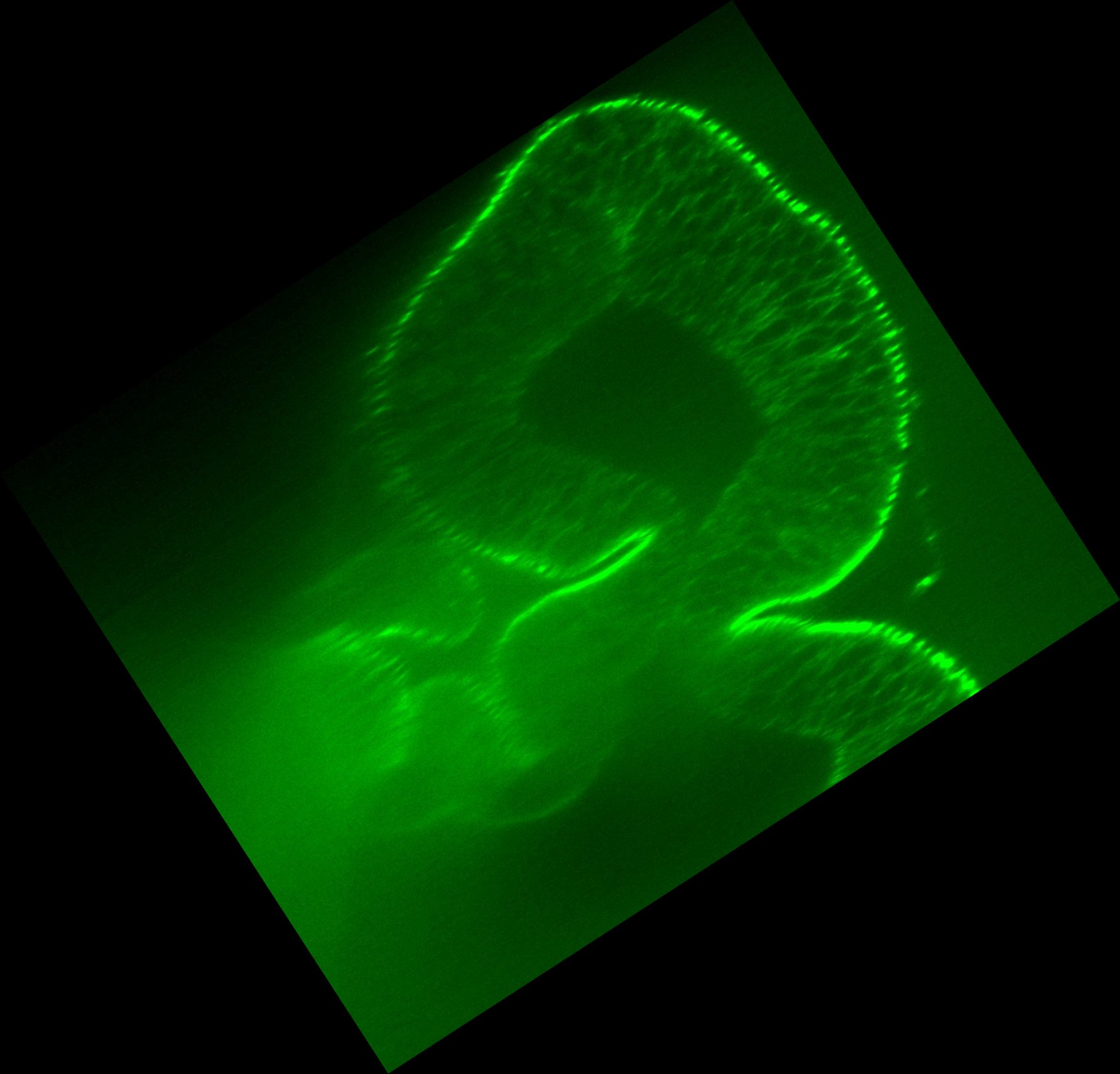

### D2.jpg

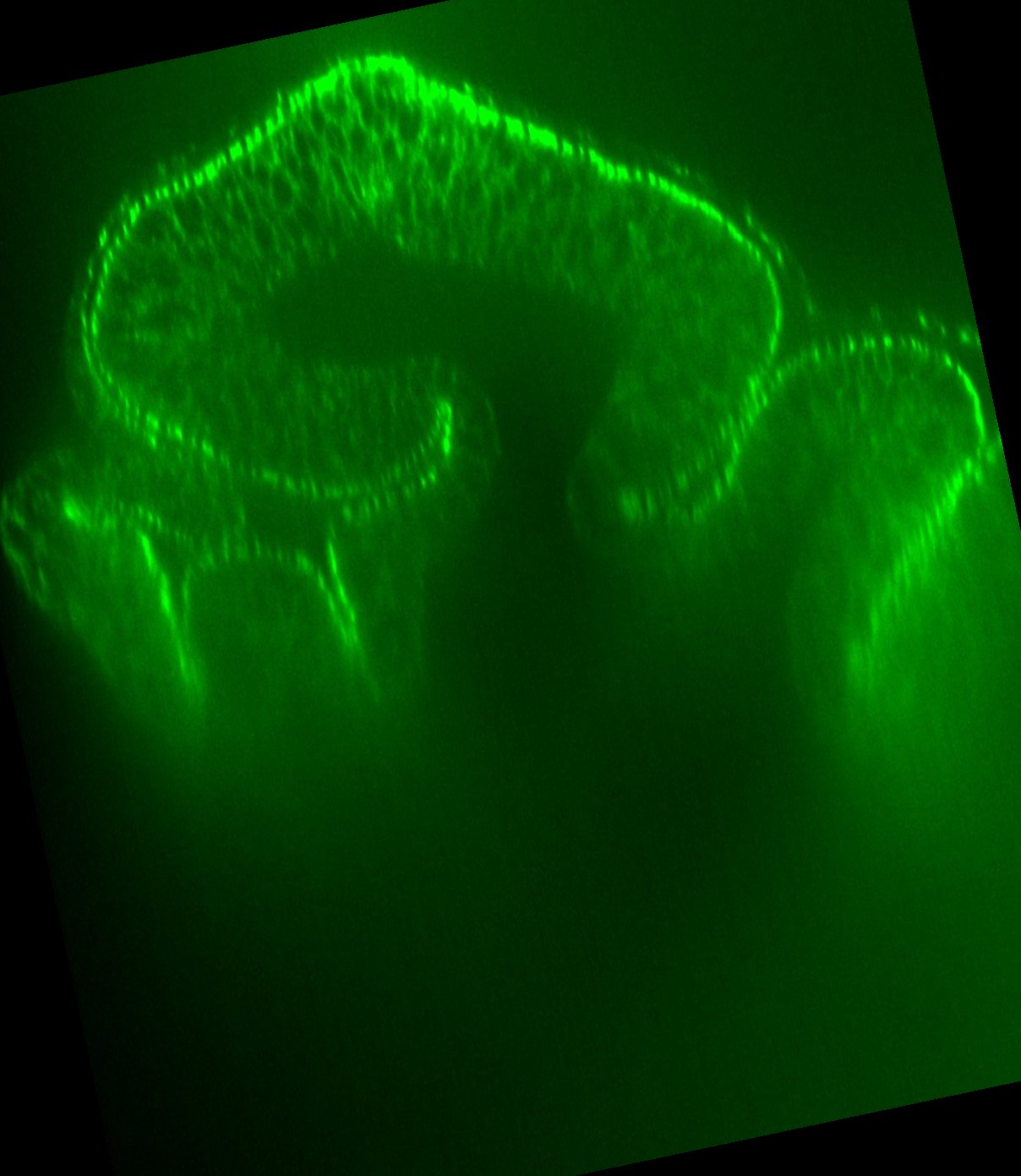

### D2.jpg

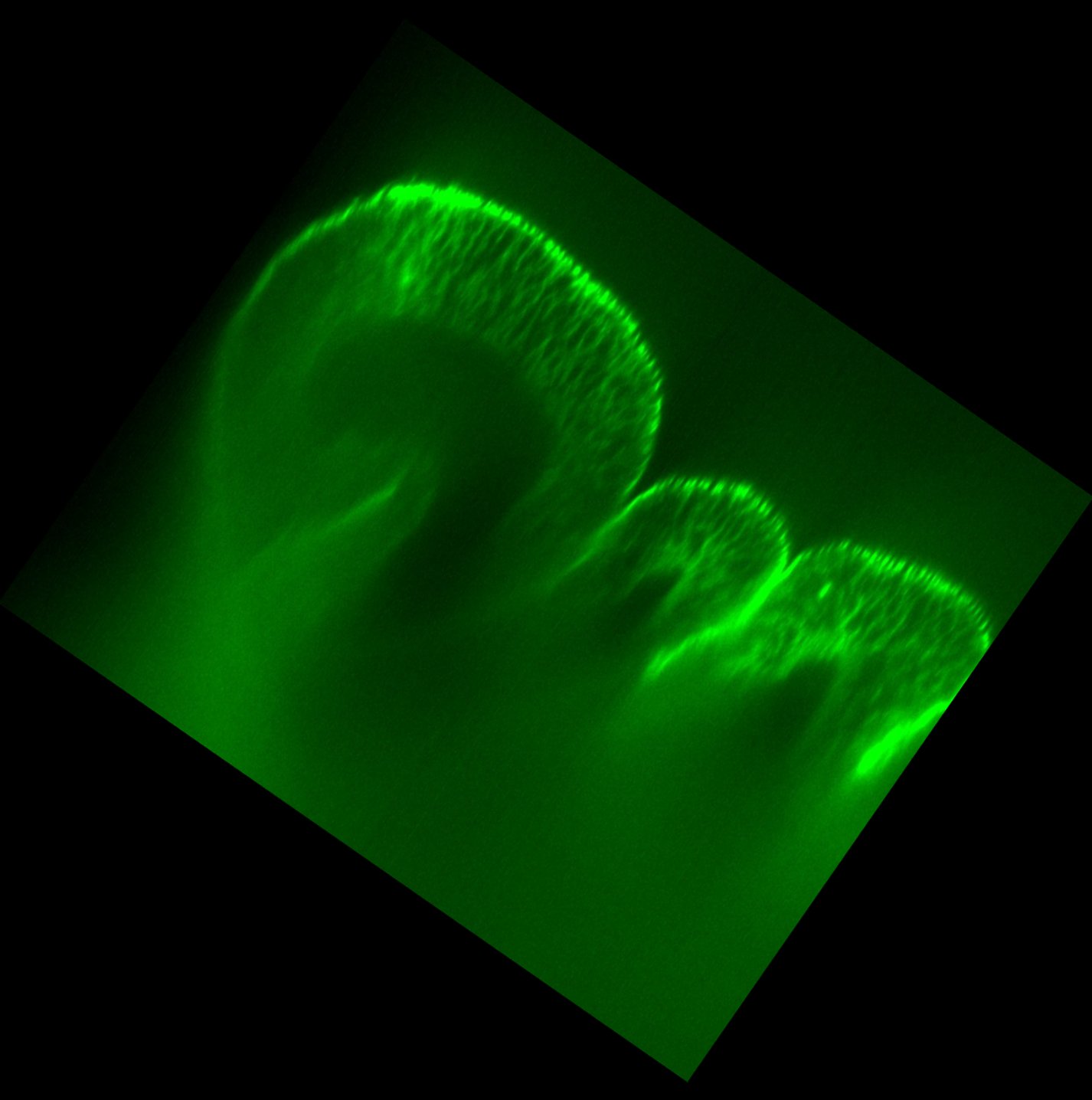

### D2_2.jpg

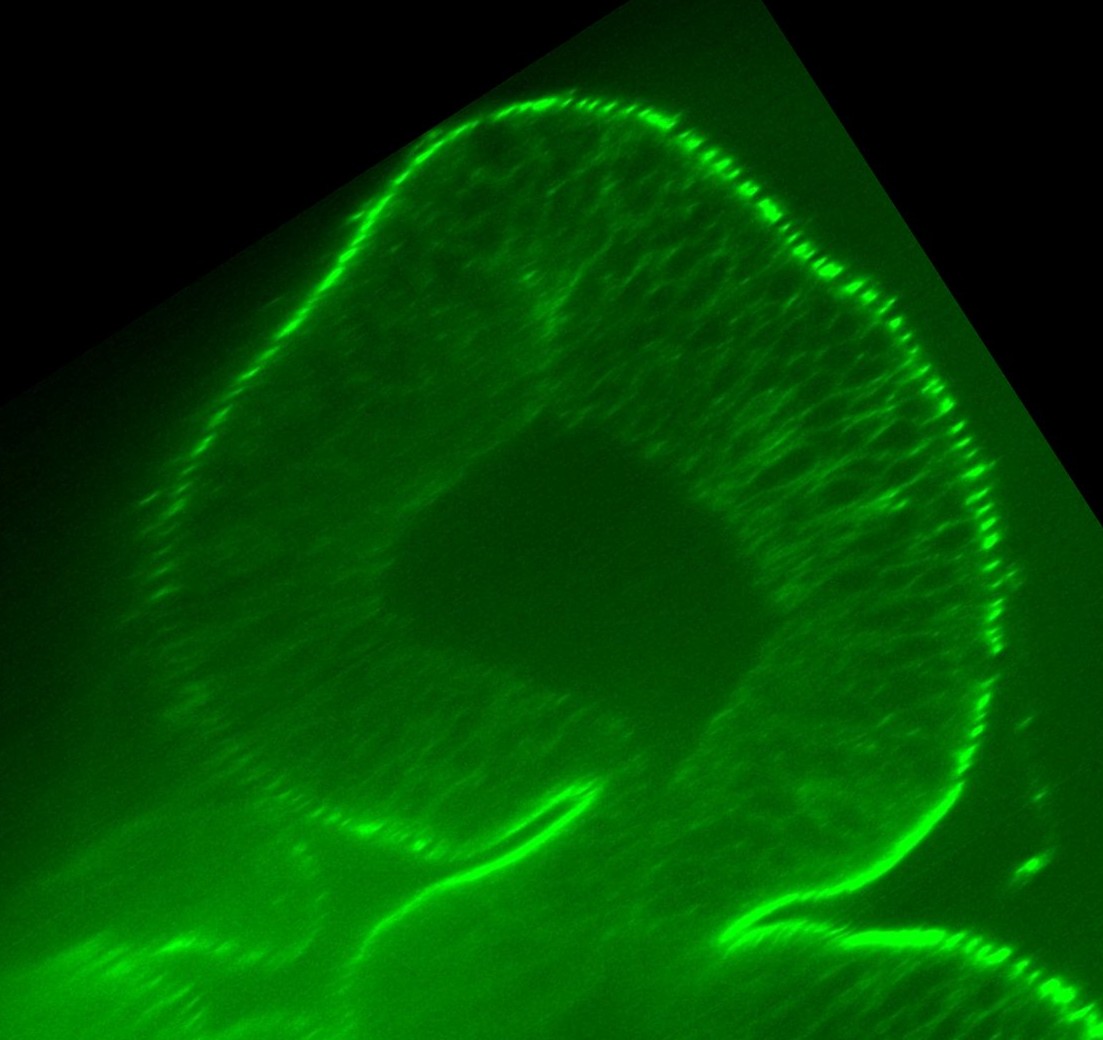

### D2Traced.jpg

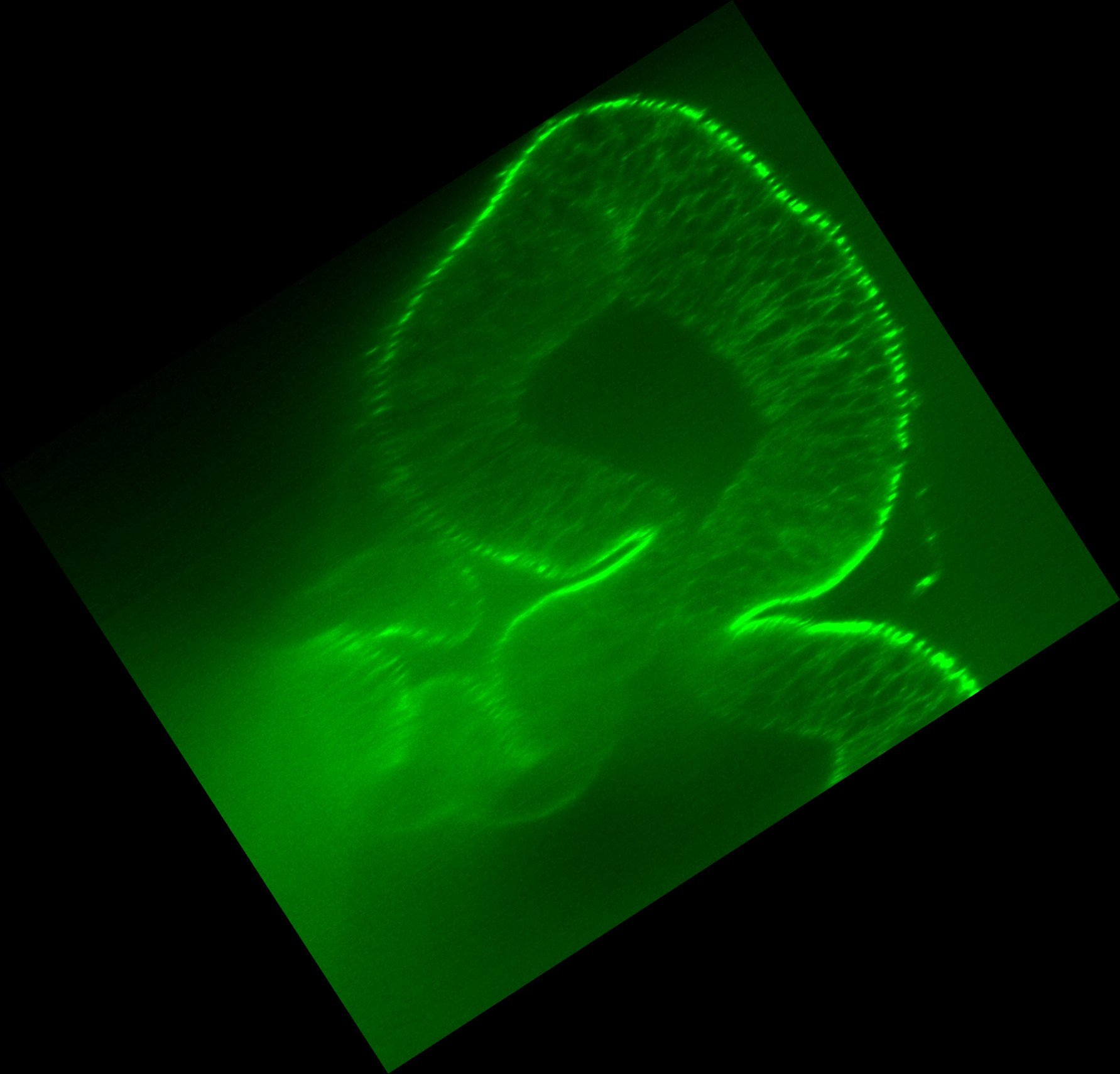

### D2Traced.jpg

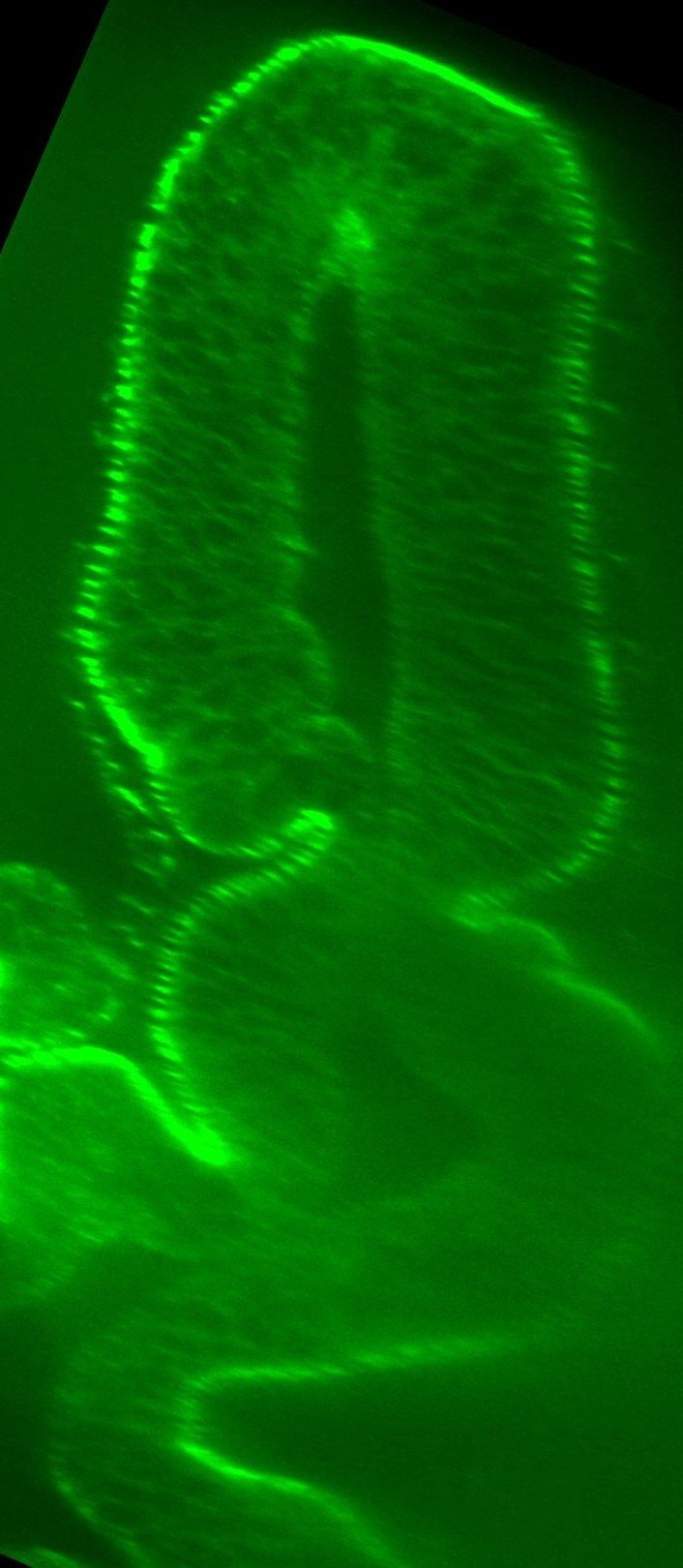

### D2Traced.jpg

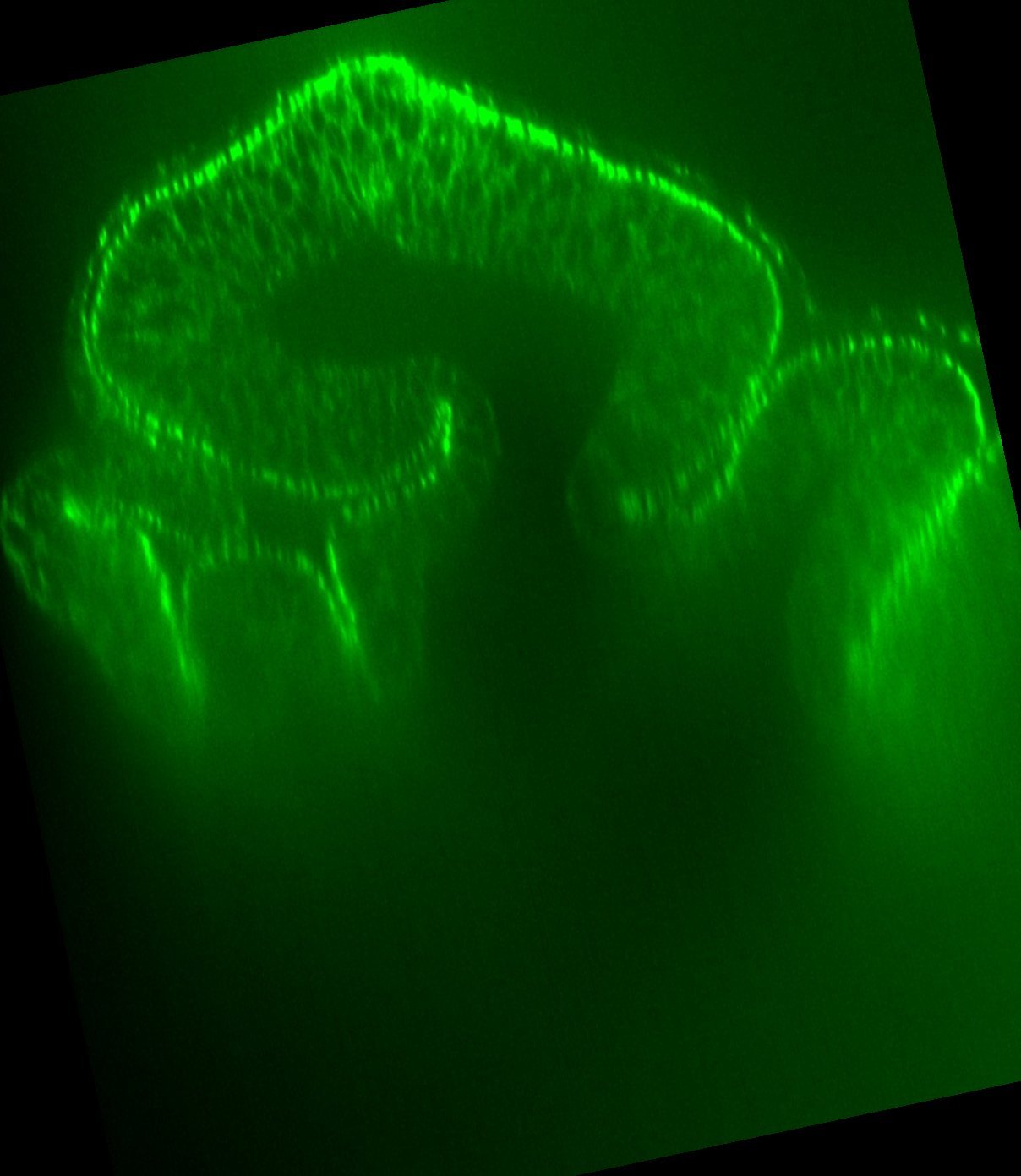

### D3.jpg

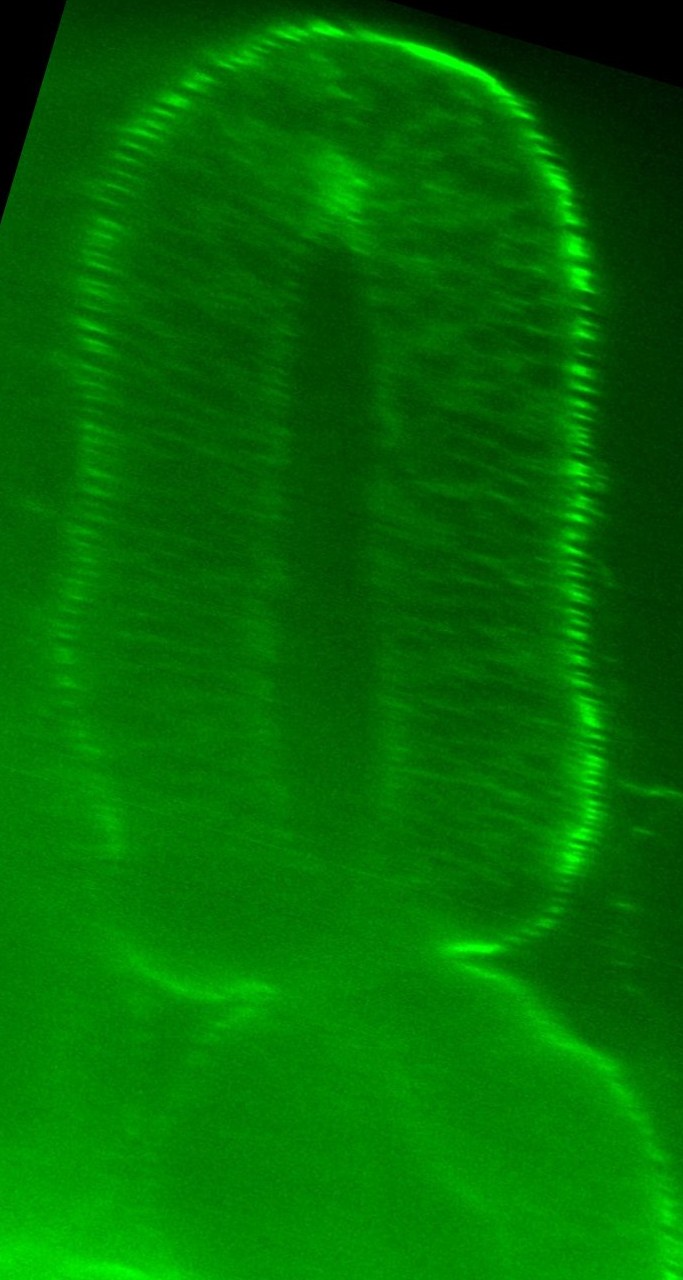

### D3.jpg

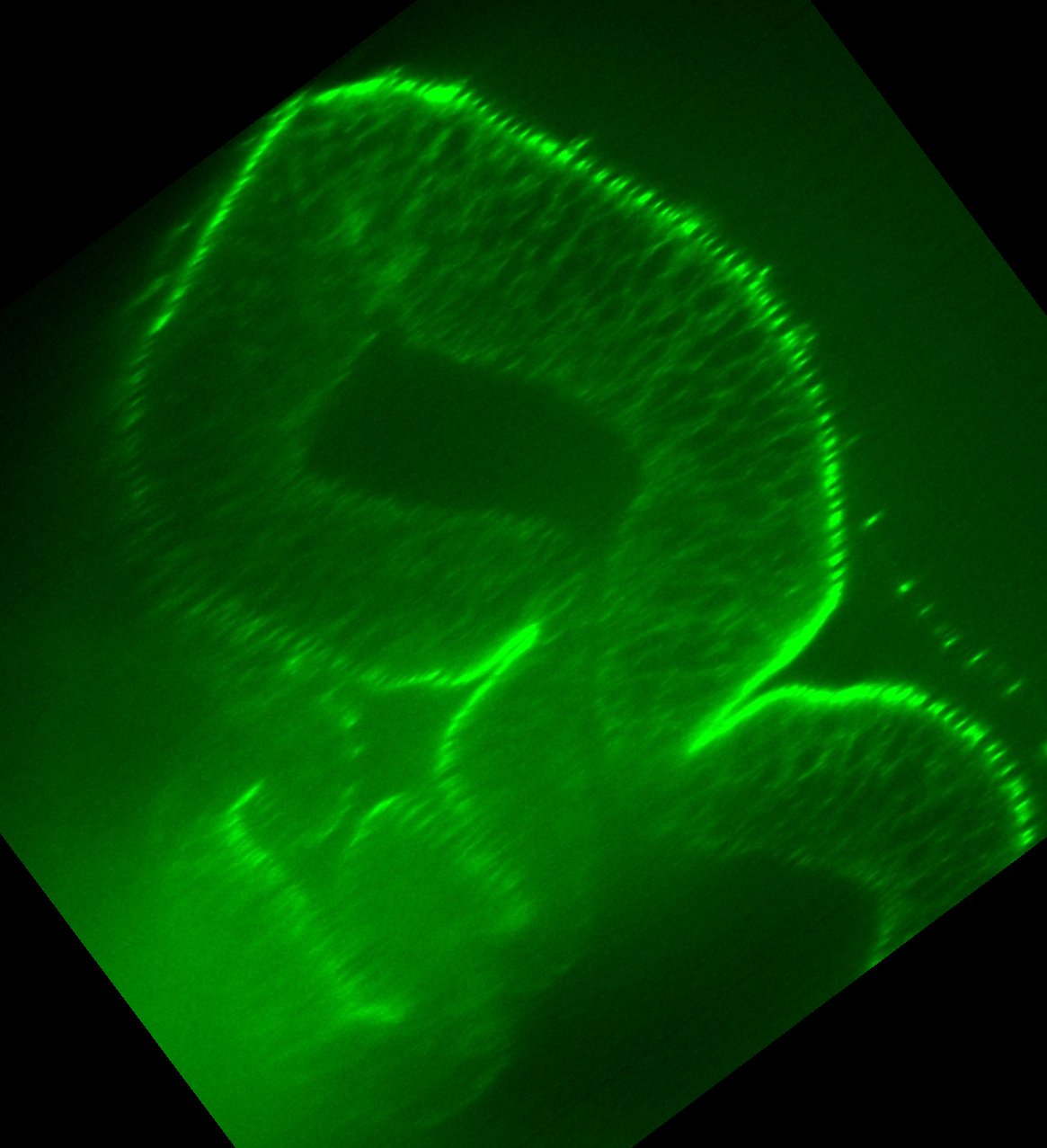

### D3.jpg

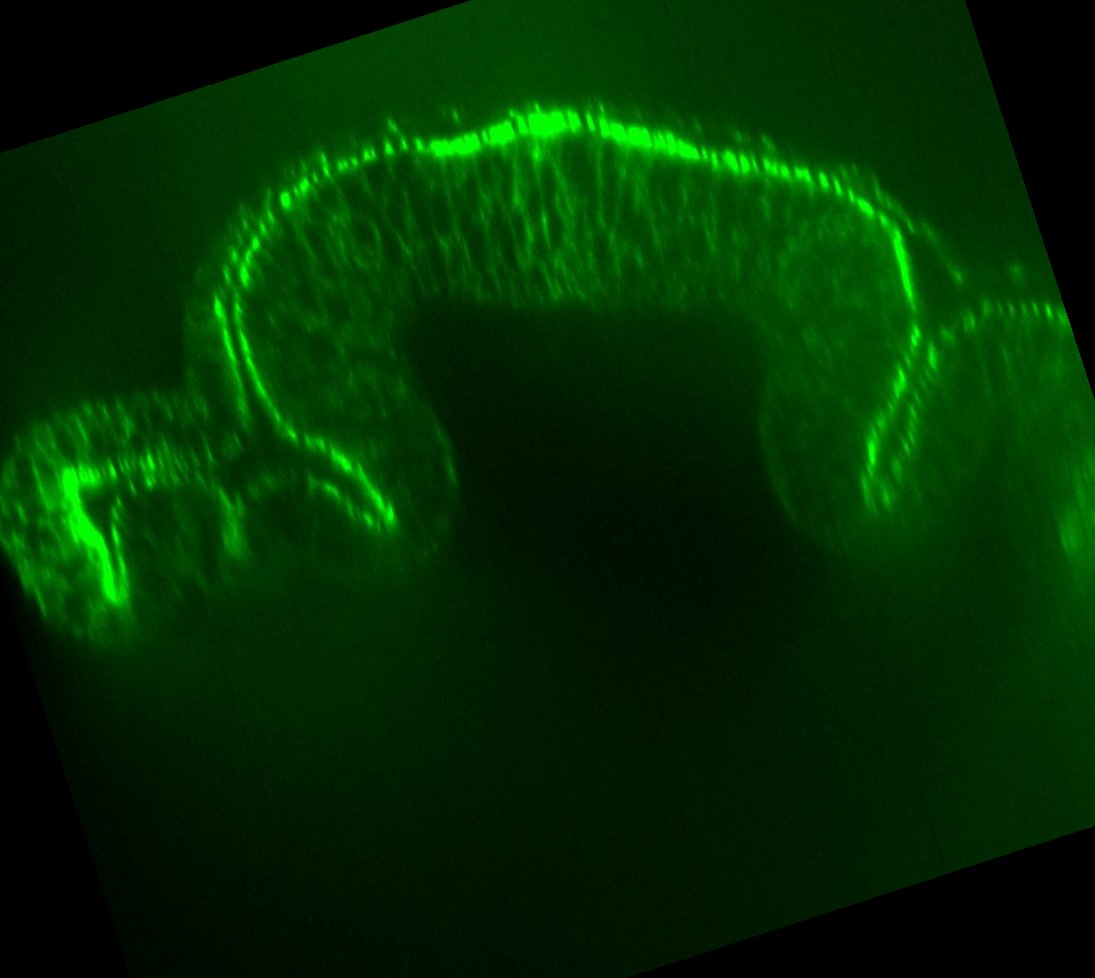

### D3.jpg

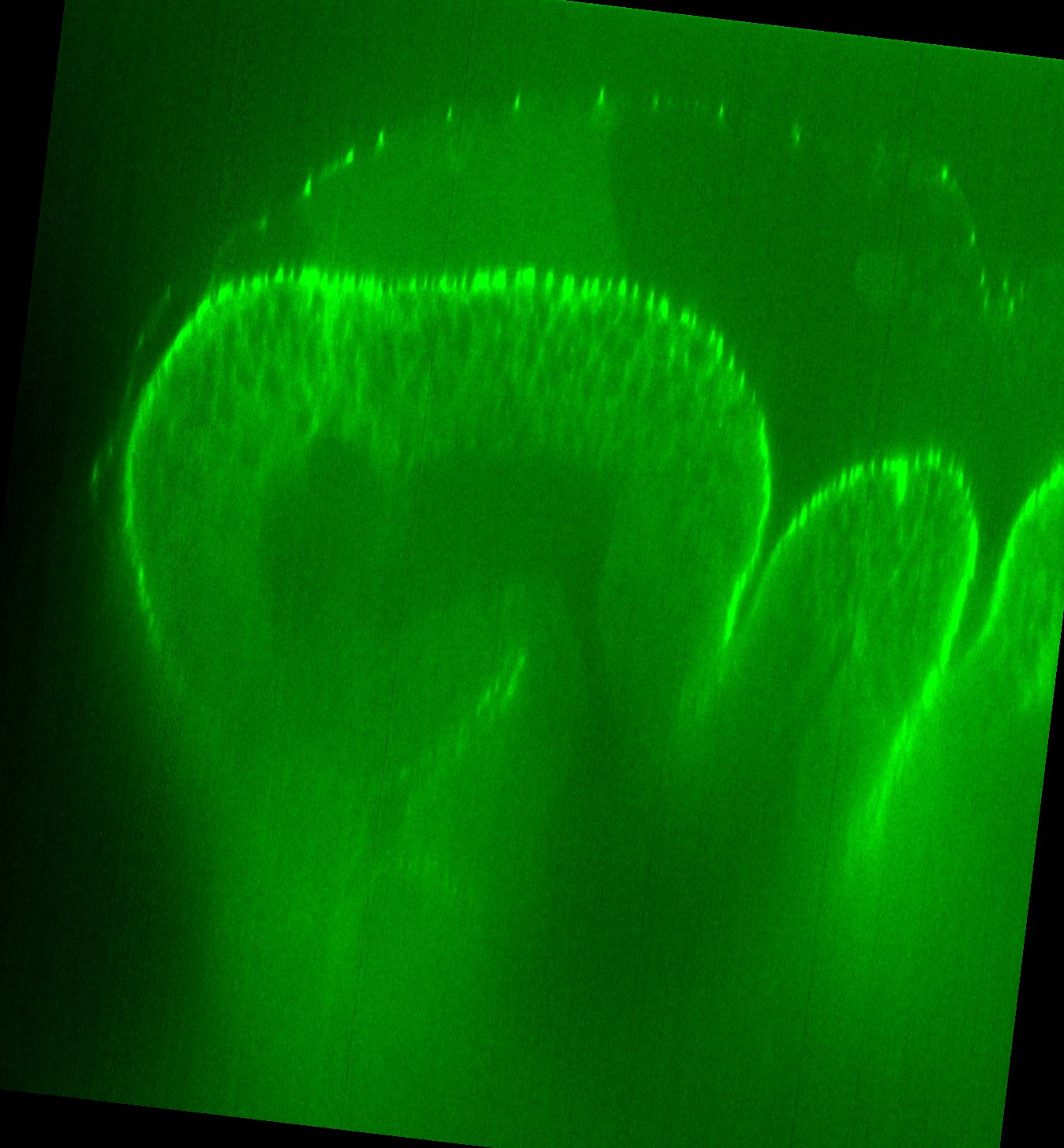

### D3_2.jpg

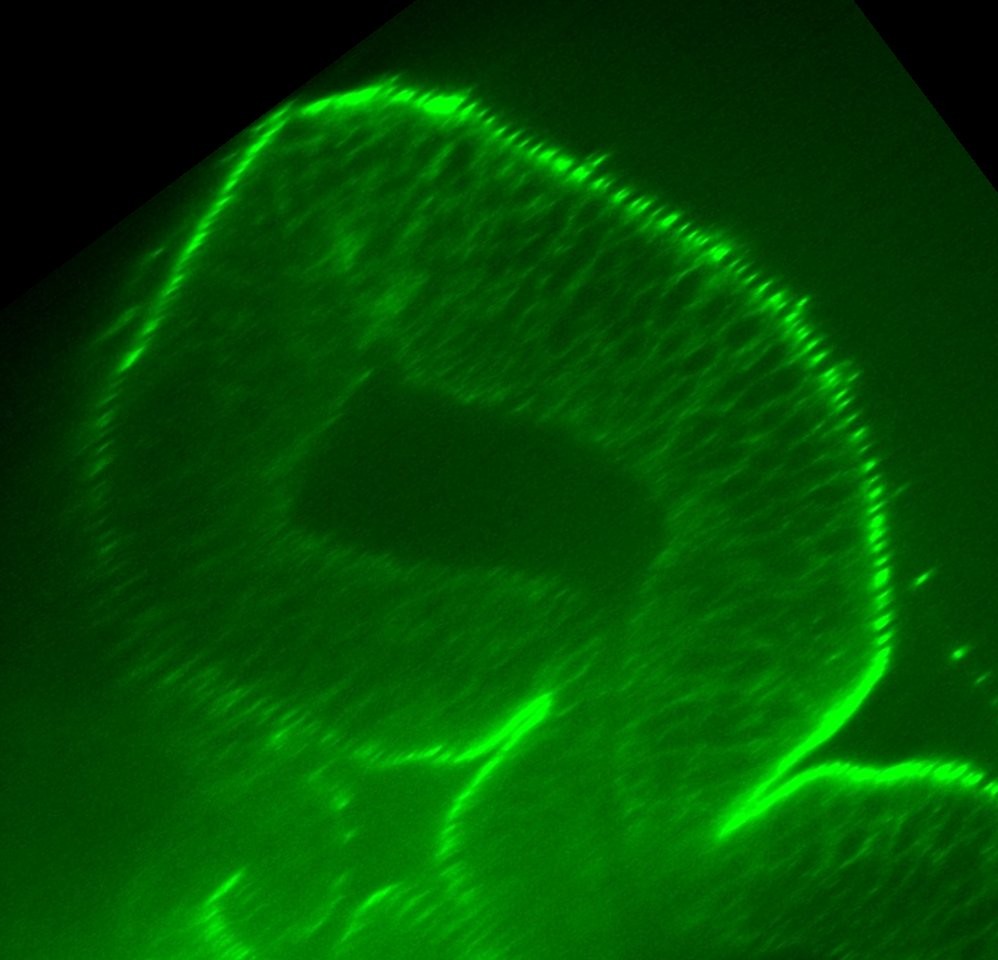

### D3Traced.jpg

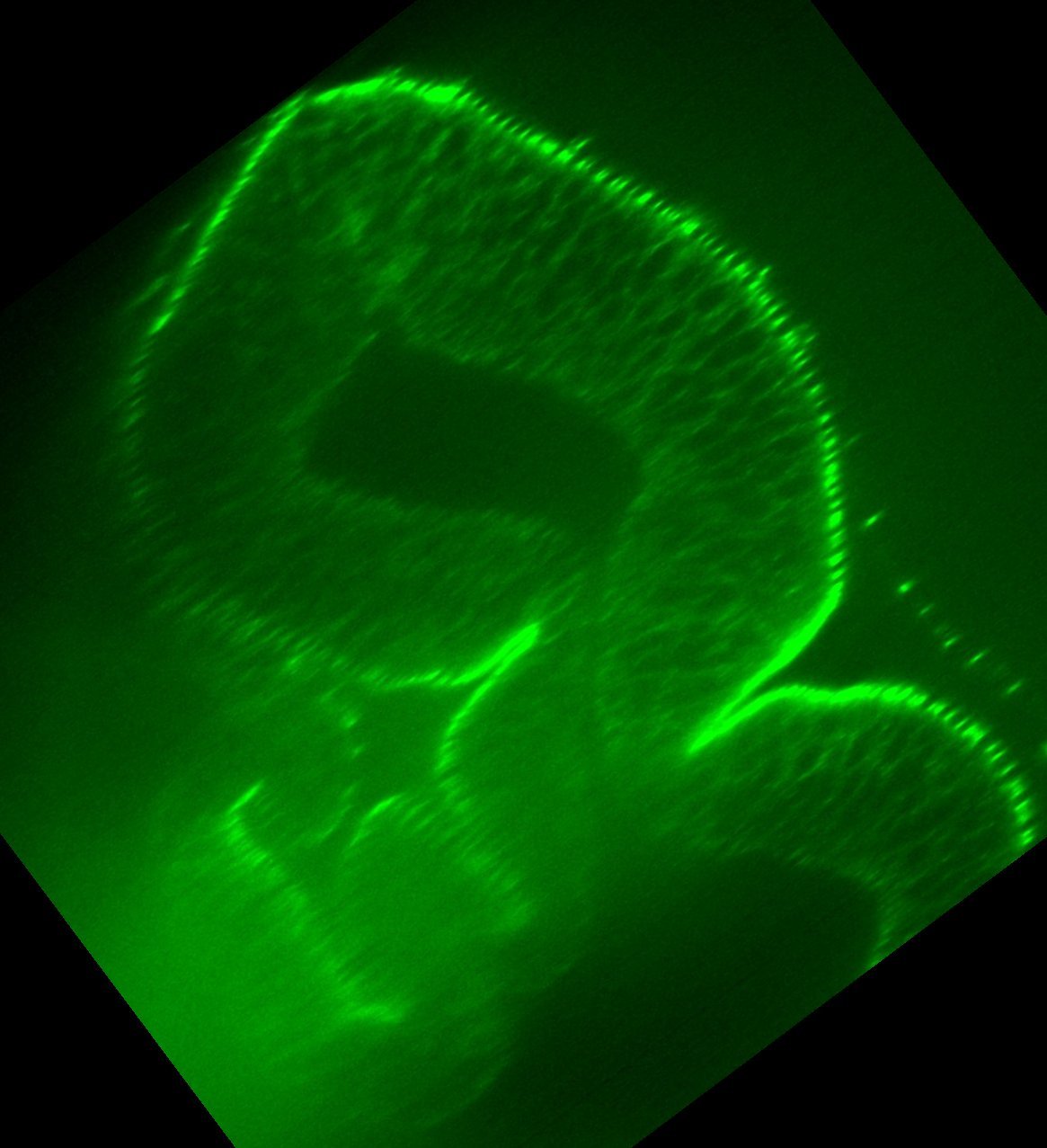

### D3Traced.jpg

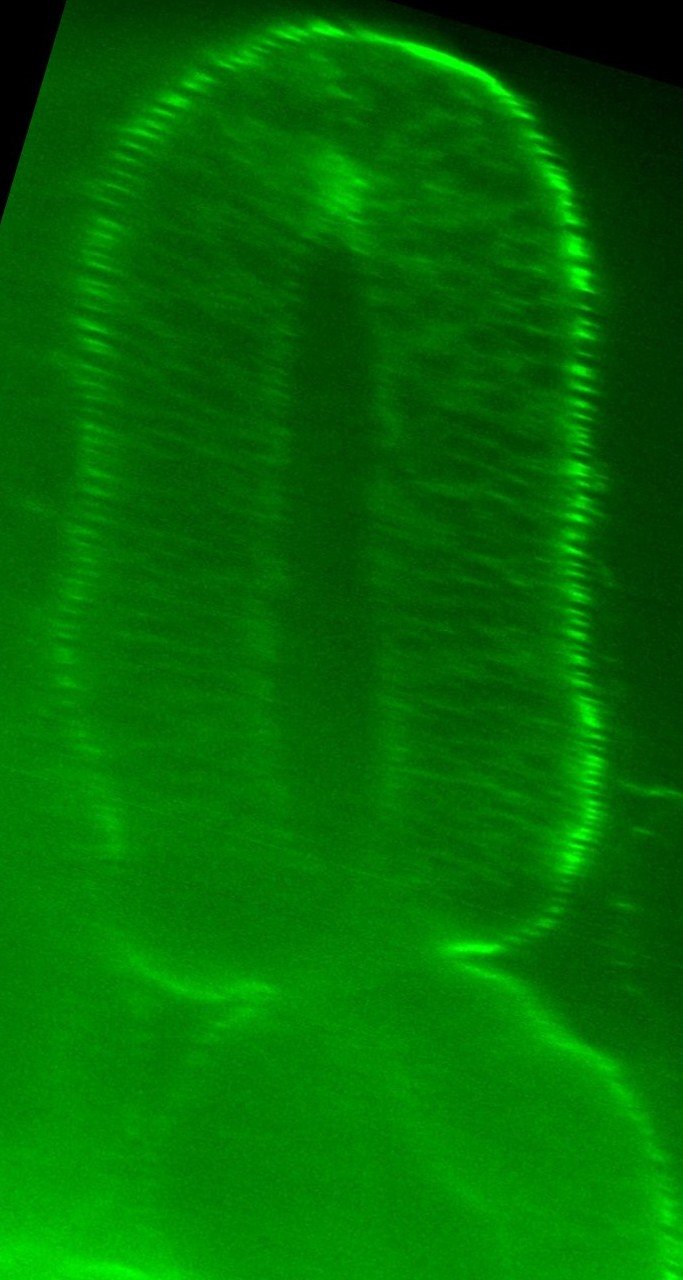

### D3Traced.jpg

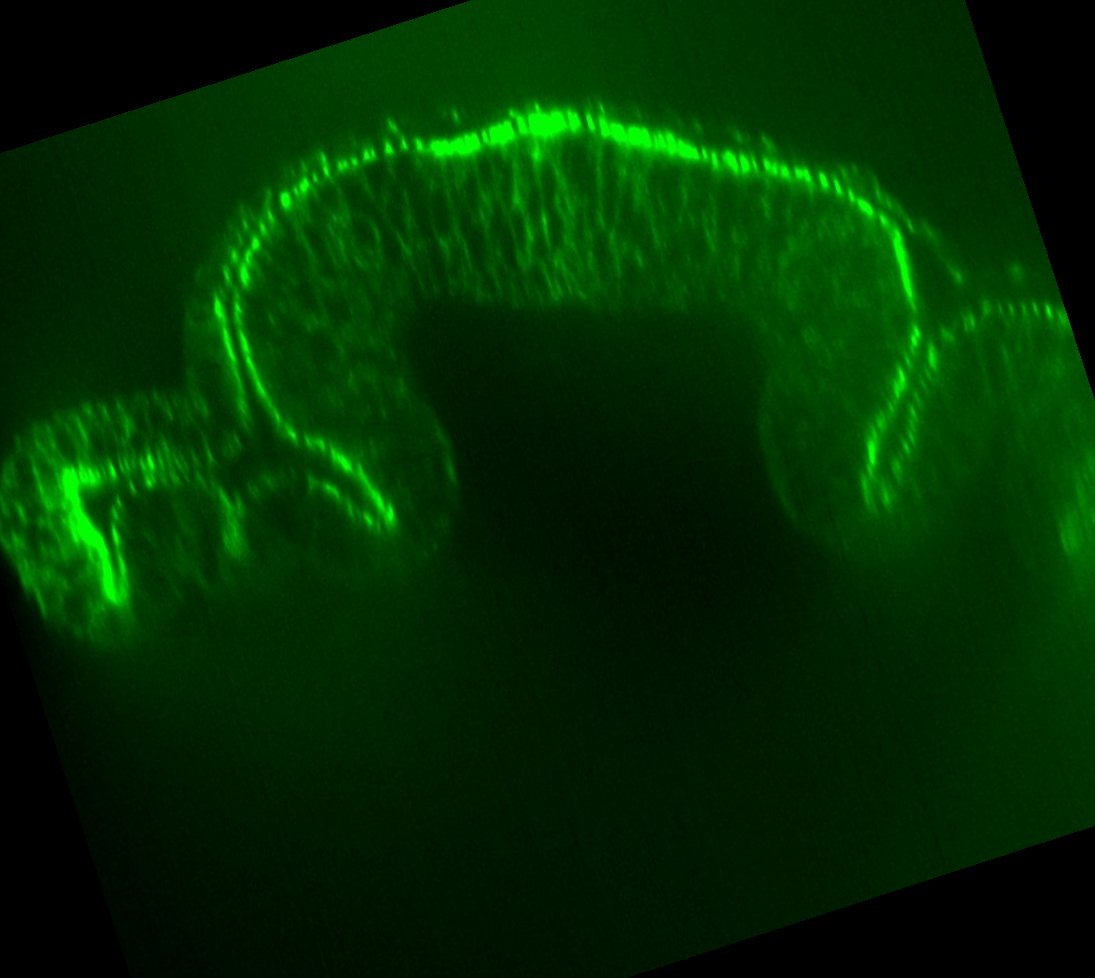

### No RA N1 Basal Surface Length 2hrAPF.jpeg

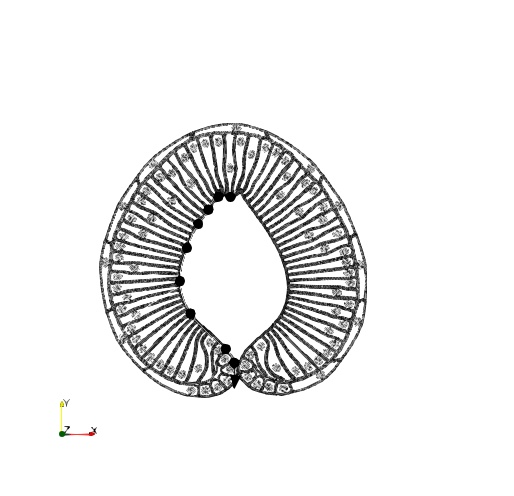

### No RA N1 Basal Surface Length 4hAPF.jpeg

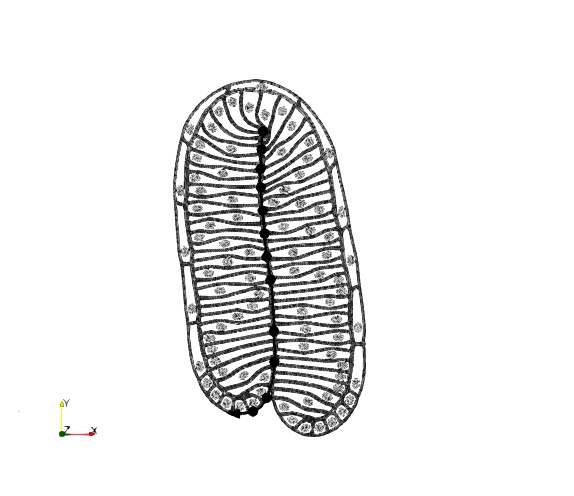

### No RA N1 Basal Surface Length wL3.jpeg

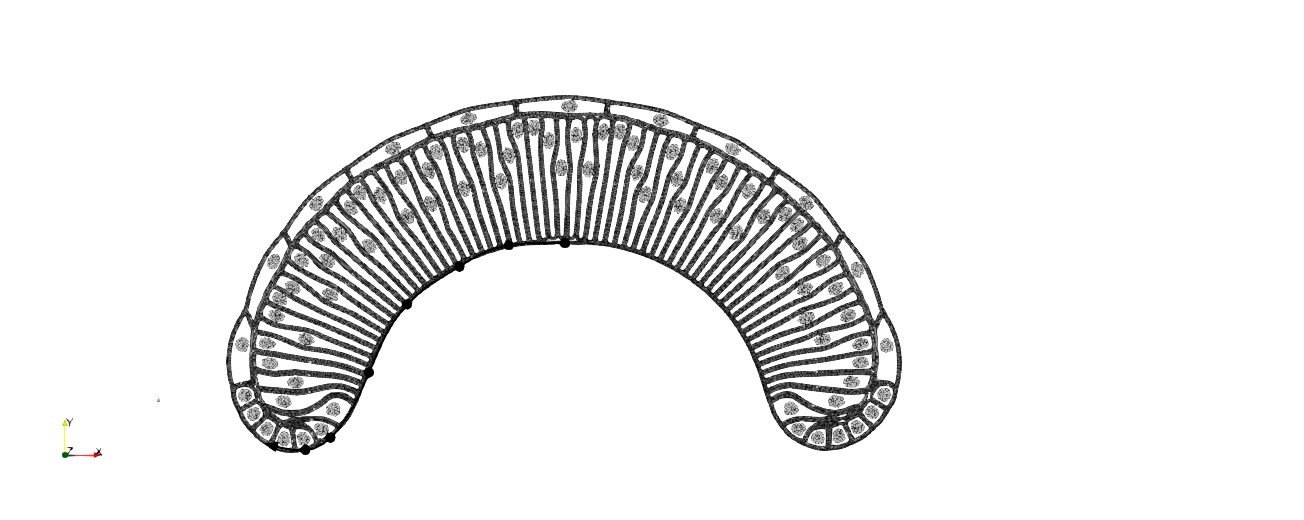

### V1aN020InitialCondition.jpg

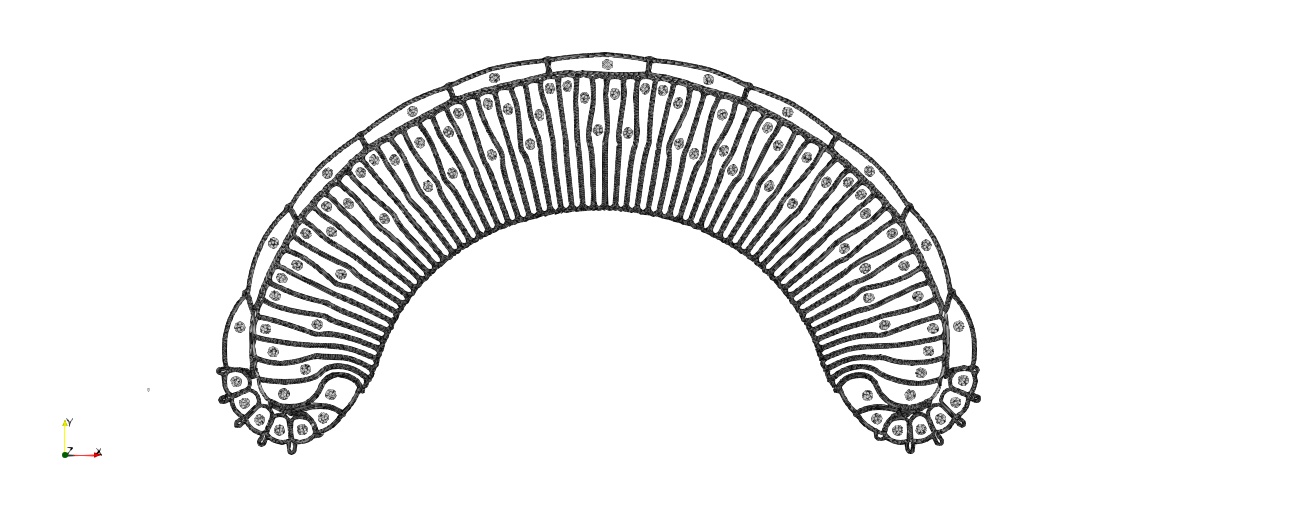

### V1awL3TissueShape.jpg

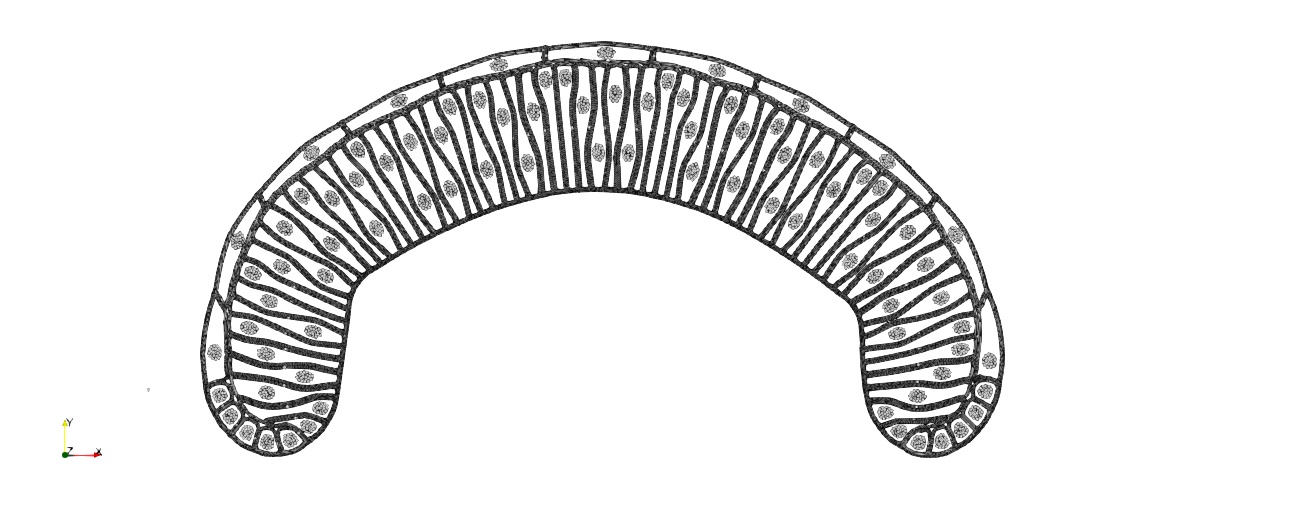

### wL3 Apical Surface.jpeg

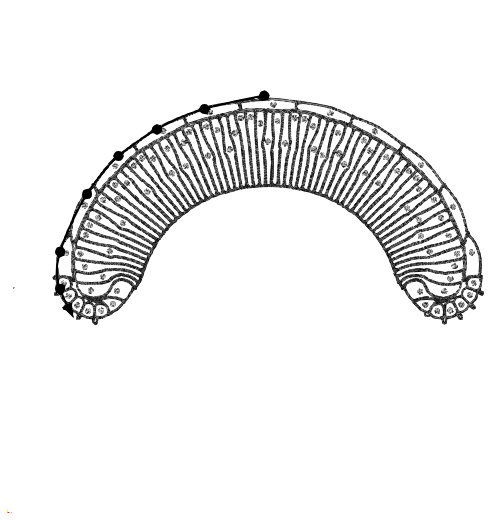

### WL3 ECAD Apical Surface.png

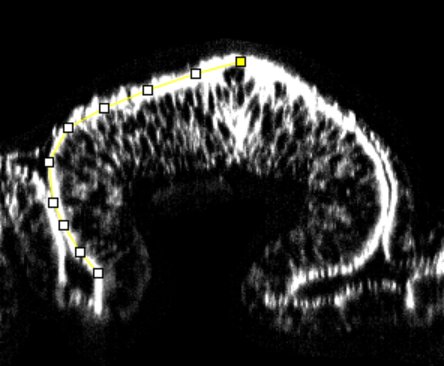

### WL3 ECAD Basal Surface.png

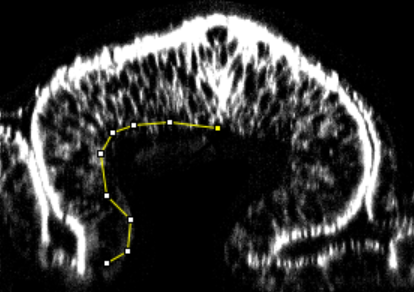

### wL3_height_overlay.png

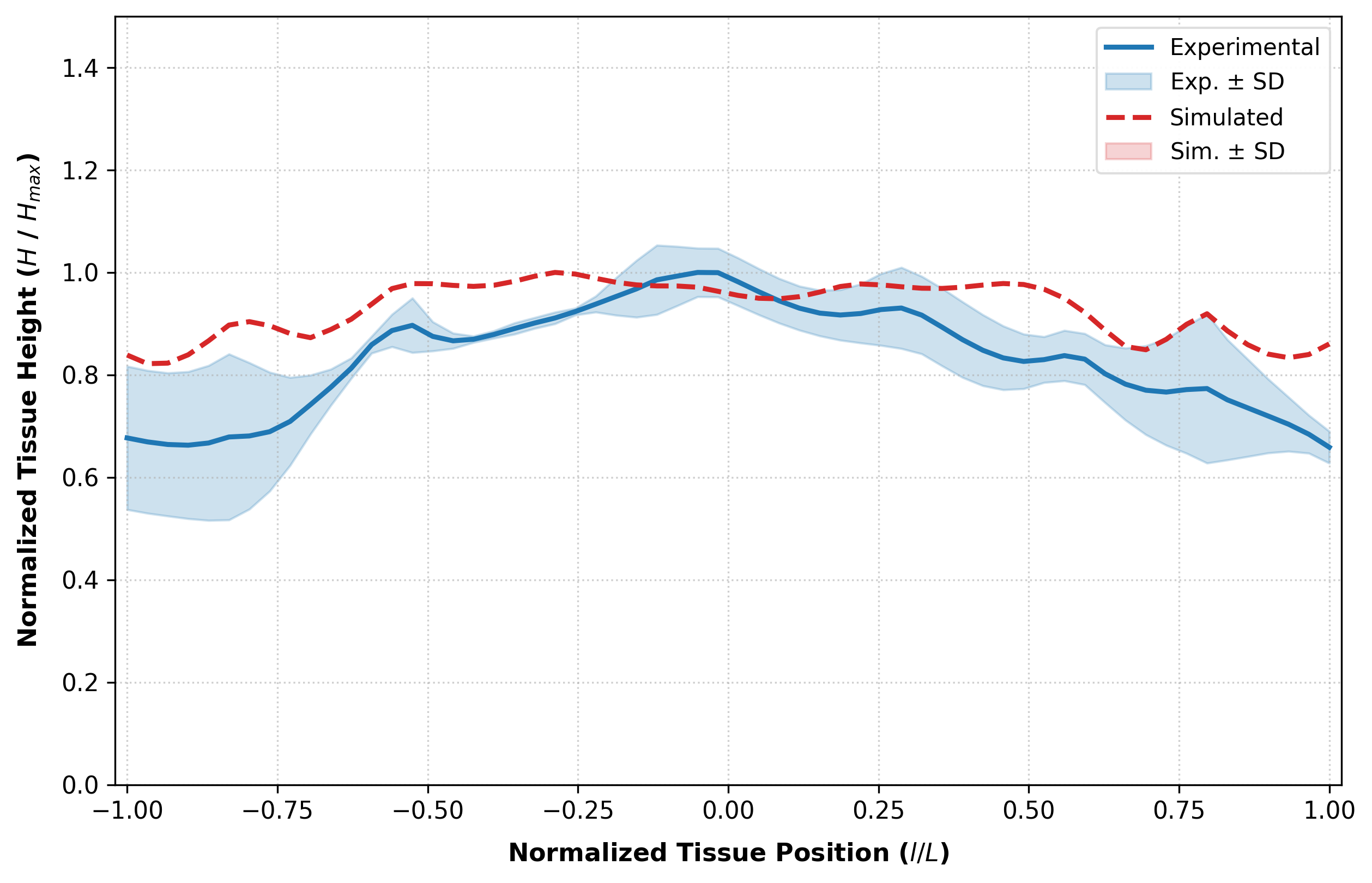

### wL3_height_overlay.png

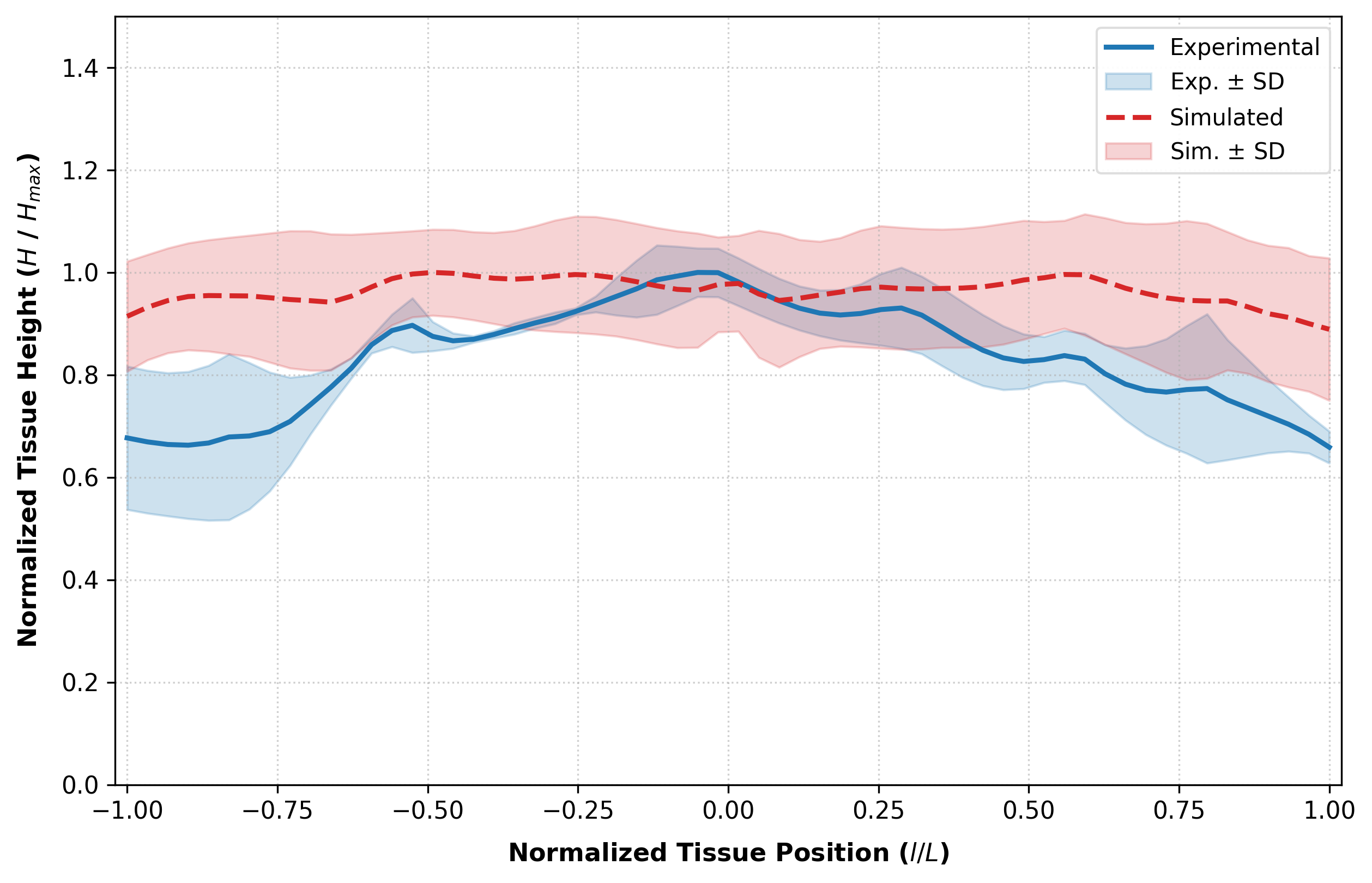

### wL3SimulatedShape.jpg

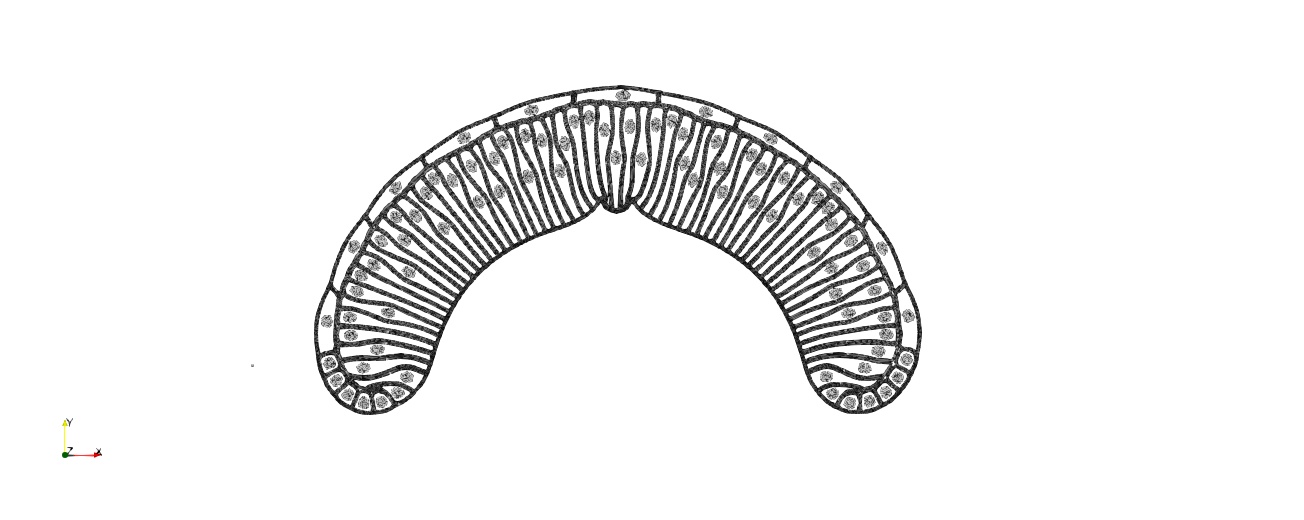
